# Exploring the specific predictive ability of multiple domains of spatial ability on STEM educational outcomes

**DOI:** 10.1101/2024.12.20.629833

**Authors:** Quan Zhou, Ziye Wang, Kaili Rimfeld, Andrea G. Allegrini, Robert Plomin, Margherita Malanchini

## Abstract

Research has implicated spatial ability as a robust predictor of aptitude, interest, and choice in STEM education and career pursuits. We address three under-explored questions regarding the role of spatial ability in STEM. First, can spatial ability consistently predict STEM success beyond other cognitive skills? Second, what aspects of spatial ability, if any, can predict success in STEM more accurately? Third, to what extent can genetic and environmental factors account for these predictions?

We addressed these questions by leveraging data from the Twins Early Development Study (N = 3,936; age range = 16-22) and using 16 tests that assessed three domains of spatial ability: navigation, object manipulation, and visualization. Results show that all three domains are highly predictive of STEM educational outcomes, especially STEM degree choice. These associations persisted after accounting for verbal and general cognitive abilities (*g*), albeit attenuated. Associations were strongest for tests of object manipulation (e.g., 2D and 3D drawing, pattern assembly and mental rotation). Genetic factors accounted for most of the observed associations between spatial ability and STEM outcomes (62% - 86%) —genetic variance was mostly shared with *g* (∼ 40%) and, to a lesser extent, verbal ability (∼ 25%).

Our findings highlight the potential utility of spatial ability as a specific predictor of success in STEM education and career choice beyond other cognitive abilities. Screening for and training spatial skills is likely useful for identifying potential, fostering talent, and improving outcomes in STEM.

**Significance Statement:** Leveraging a comprehensive battery of 16 spatial ability tests across multiple domains, we show that spatial ability has specific utility for predicting success in STEM education. Spatial skills predict success in STEM above and beyond other cognitive abilities, particularly when it comes to STEM engagement and pursuing further STEM education. Thus, spatial skills may be a fruitful target for policymakers, stakeholders, and industries looking to develop interventions, identify and foster talent, and reduce outcome disparities.

## Introduction

Science, technology, engineering, and mathematics (STEM) education is highly important in today’s technologically oriented world. At the individual level, success in STEM education is associated with positive life outcomes, including income, health, and longevity (1, 2). For governments worldwide, promoting student engagement in STEM is increasingly recognized as an investment in human capital development— an imperative for nations looking to remain competitive in the global economy (3–5). Thus, perhaps now more than ever, it is essential for social scientists to study how to foster engagement and talent in STEM to inform the development and implementation of evidence-based policies and educational programs.

To this end, research has investigated spatial ability as a predictor of aptitude, interest, and choice in STEM educational and career pursuits (6–8). Although theoretical discordances exist regarding the exact structure of the construct, spatial ability can be broadly defined as “the ability to generate, retain, retrieve, and transform well-structured visual images” (9). Typically, spatial ability tests assess individuals’ abilities to mentally rotate or reconfigure shapes or objects, visualize them from different perspectives, and make judgments about the operations of mechanical stimuli (10). Dating back to the earliest modern cognitive tests, spatial ability tests were and continue to be found in almost all batteries of general intelligence, historically and psychometrically situating it as a domain of general cognitive ability, or *g* (9, 11).

Much of the work on the association between spatial skills and STEM educational outcomes has conceptualized spatial ability within a simplified version of Carroll’s hierarchical model of *g*, in which tests are categorized within three content domains: mathematical, spatial, and verbal (12). Despite high inter-correlations between these three domains, longitudinal research in samples of gifted youth has consistently uncovered markedly different cognitive domain profiles between STEM and humanities students (6, 7, 13). That is, individuals scoring higher in the spatial domain relative to the verbal domain had a greater inclination towards STEM disciplines compared to humanities disciplines with respect to interest, degree choice, and occupation. In contrast, the opposite was true for those who scored higher in verbal relative to spatial (13, 14). Other studies have found that spatial ability added incremental predictive validity for STEM success beyond that explained by mathematics-verbal proxies of cognitive abilities (e.g., the SAT; (6, 13)). Altogether, this literature suggests that spatial ability, though putatively a domain of general cognitive ability, has specific relevance for STEM success.

The current study addresses three key gaps in the extant literature on the relationship between spatial ability and STEM. First, although spatial ability is generally acknowledged to contribute to *g*, the extent to which spatial ability predicts STEM success beyond other cognitive abilities remains unclear. To address this first aim, we tested the associations between spatial ability and STEM outcomes at three levels of granularity: first, we examined the link between spatial ability and STEM outcomes; second, we assessed their associations after accounting for verbal ability; and third, we tested whether their associations remained after considering general cognitive ability, the most stringent control. We compared the predictive utility of spatial ability across STEM and humanities subjects to unpack the specificity of the prediction further.

Second, measures of spatial ability used in research have largely focused on narrow conceptualizations of the construct (i.e., object-oriented abilities) while neglecting other domains of spatial skills (e.g., large-scale visualization and navigation abilities) that have traditionally been difficult to measure. Thus, little is known about the predictive role of navigational skills for STEM outcomes, nor whether differential predictive utility exists across different spatial domains and STEM subjects (i.e., are certain domains associated with certain STEM subjects more highly than others?). In the current study, we leverage previous work conducted by our research team, who investigated the structure of spatial ability measured with two innovative online batteries of 16 spatial ability tests (these batteries and their development are described in detail in the Methods and Materials section) (15, 16). Using exploratory and confirmatory factor analysis, our previous work uncovered a hierarchical structure of spatial ability in which three broad domains (navigation, object manipulation, and visualization) were largely captured by a higher-order general factor of spatial ability (16). In the present study, we explore the predictive ability of these three domains of spatial ability (as well as that of the general spatial ability factor) on STEM engagement and achievement, measured as subject choice, degree choice, and academic achievement. Our second aim was to investigate the extent to which these three domains of spatial ability differentially predict STEM engagement and achievement.

Finally, limited work has examined the contribution of genetic and environmental factors in explaining individual differences in spatial ability. To date, behavioral genetic studies of spatial ability have typically been limited by small sample sizes and measures that vary widely across studies (17, 18). As a result, estimates of etiological influence have varied widely (e.g., a meta-analysis on twin studies of “spatial reasoning” found heritability estimates ranging from 0.12 to 0.94 (17)), while efforts to consolidate these estimates are undermined by a lack of measurement invariance across studies. Our previous work examined the genetic and environmental influences underlying the variation (and covariation with *g*) in our comprehensive spatial ability batteries described above using the twin design to address this issue. Results suggested that spatial ability and its domains were partly heritable and, critically, displayed a large degree of genetic independence from *g* (45%) (16). Given this, our third aim in the present study was to examine the extent to which shared genetic and environmental factors accounted for the links between spatial ability and STEM outcomes before and after accounting for shared variance with *g* or verbal ability. Analyses were pre-registered with the Open Science Framework (https://osf.io/48kwa/).

## Results

### Creating the structure of spatial ability, verbal ability, and *g* through confirmatory factor analysis

Leveraging our previous work, we first conducted confirmatory factory analyses (19) on our measures of (a) spatial ability, (b) verbal ability, and (c) general cognitive ability (*g*) before extracting factor scores from each model (See Methods and Materials for additional details and SI Appendix, Figures S1-S4 for factor loadings, model fit statistics and correlations between factors). Descriptive statistics for each measure are displayed in SI Appendix, Tables S1-S3. All CFA models provided a good fit for the data, while extracted factor scores approaching normal distributions with acceptable values for skew and kurtosis (SI Appendix, Table S4).

### Spatial ability predicts STEM success beyond other cognitive abilities

We first examined the extent to which spatial ability predicted STEM and humanities educational outcomes. We explored these associations at three levels of granularity: first, we examined the direct link between spatial ability and educational outcomes; second, we examined their associations after accounting for verbal ability; and lastly, after accounting for *g*. Figure 1 presents the results of this first set of analyses. Full model statistics (as well as descriptive statistics for the outcome variables) are presented in SI Appendix, Tables S5-S10).

**Figure 1.**
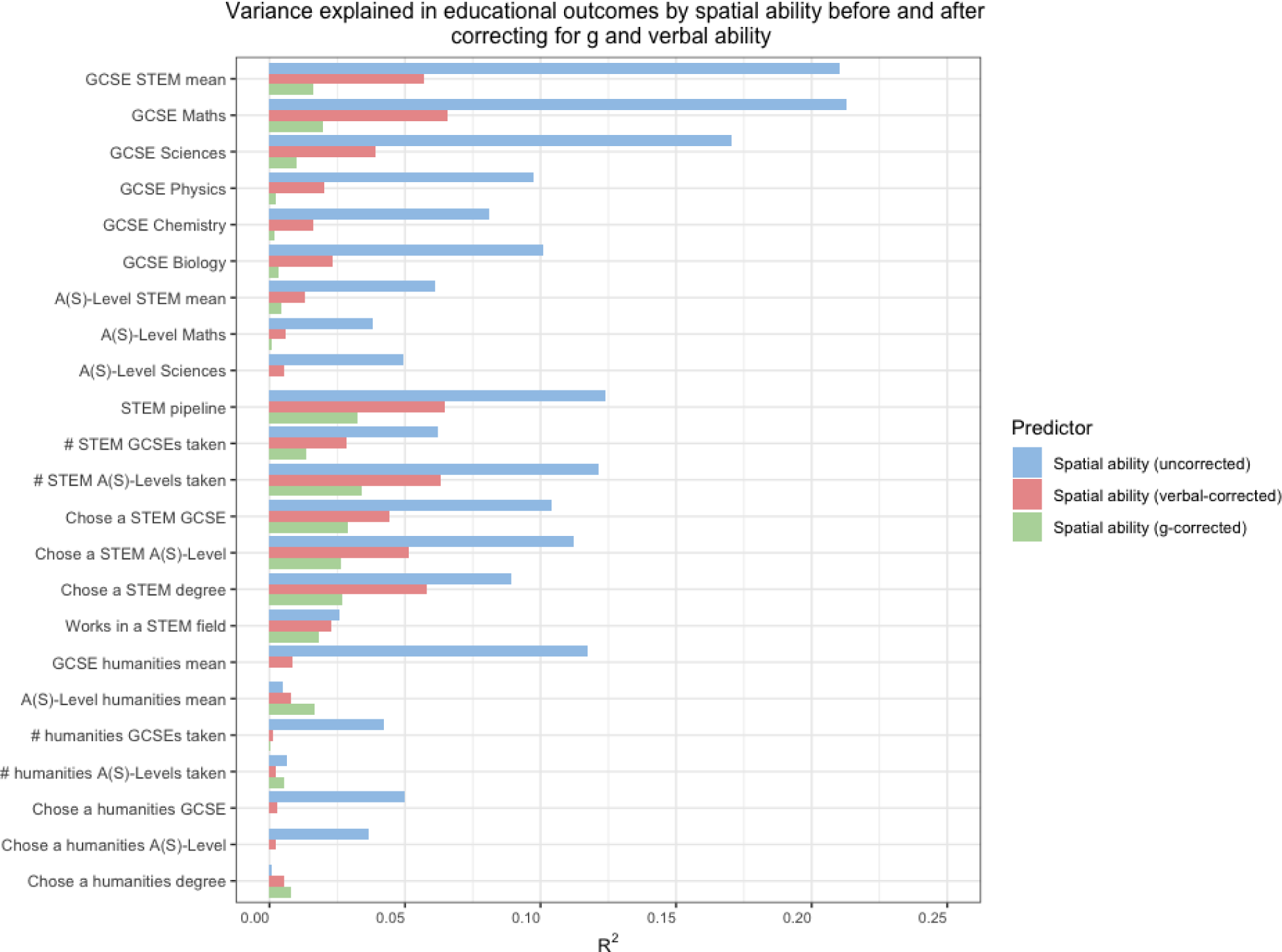
Variance explained in both continuous (R2) and binary (Nagelkerke R2) educational outcomes by the general factor of spatial ability uncorrected, corrected for verbal ability, and corrected for g.

The general factor of spatial ability predicted all educational outcomes, with the associations being significantly larger for STEM outcomes as compared to humanities outcomes. For continuous outcomes, the strongest effect size for the general factor of spatial ability was observed for *GCSE Mathematics grade* (β=0.46, 95% CI [0.42, 0.50], *R*^2^ = 0.23), while the weakest effect size was observed for the *number of humanities A-Levels taken* (β=0.09. 95% CI [0.03, 0.14], *R*^2^ = 0.01). For binary outcomes, the largest odds ratio for the general factor was observed for *choosing a non-compulsory STEM GCSE* (OR = 2.27, 95% CI [1.91, 2.73], Nagelkerke *R*^2^ = 0.11), while the smallest was observed for *choosing a humanities degree* (OR = 1.05, 95% CI [0.93, 1.18], Nagelkerke *R*^2^ = 0.00). See more details in SI Appendix, Tables S10.

Figure 1 also presents the verbal-corrected and *g*-corrected predictions for the same continuous and binary outcomes in terms of *R*^2^ and Nagelkerke *R*^2^, respectively. A general pattern of attenuation in variance was observed going from uncorrected to verbal-corrected to *g*-corrected models, indicating that both verbal ability and *g*—but especially *g*—index a large portion of phenotypic variance shared with spatial ability. Critically, however, corrected associations remained significant and non-negligible in terms of variance explained for many STEM outcomes, for example choosing a STEM subject in middle (GCSE) and high school (A levels), but not for humanities outcomes.

Interestingly, the largest associations after correcting for other cognitive abilities were observed for outcomes related to engagement and choice (e.g., STEM pipeline) rather than achievement scores (e.g., *GCSE Mathematics* and *GCSE STEM mean*). The largest of these corrected associations was observed for *STEM pipeline* (*R*^2^ = 0.04 after correcting for *g*), a 4-level ordinal variable we created that denotes how far students proceed in STEM education (see Methods and Materials for a detailed explanation of how this variable was computed). Similarly, associations remained for *choosing a non-compulsory STEM GCSE*, *choosing a STEM A(S)-Level*, and *choosing a STEM degree* (*R*^2^ = 0.03 after correcting for *g* for all), highlighting how spatial ability, independent of *g*, seems to have specific predictive utility for engagement and interest in STEM rather than achievement. Figure 2 highlights the distributions of the spatial ability score as a function of STEM pipeline score. Spatial abilities differed significantly at every stage of the STEM pipeline with the lowest scores on average observed for those students who left the STEM pipeline before middle school and the highest spatial ability scores observed for those students who chose a STEM degree.

**Figure 2.**
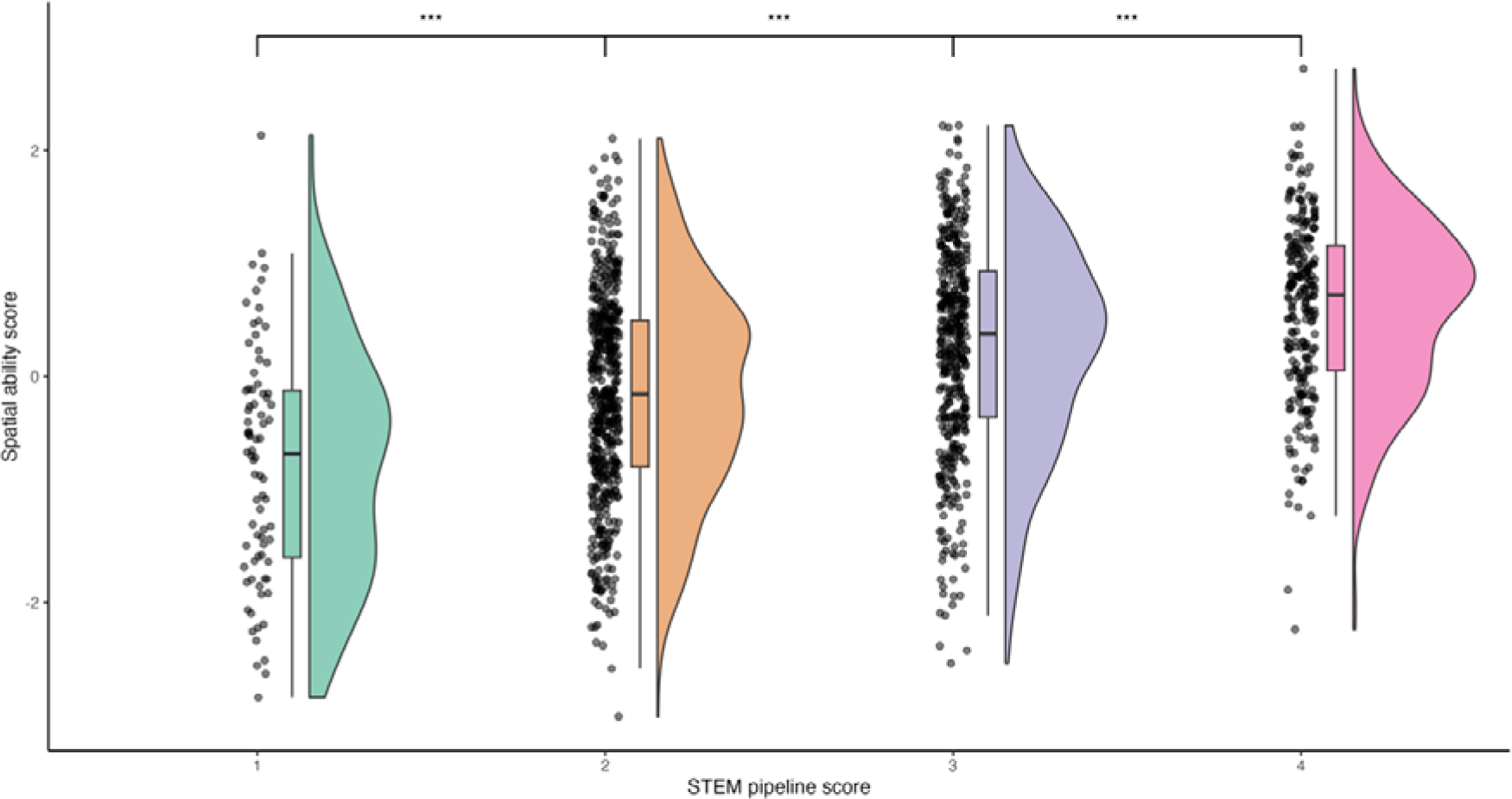
Rainbow cloud plot of spatial ability score as a function of STEM pipeline score, where 1 = did not choose any non-compulsory STEM GCSEs, 2 = chose at least one non-compulsory STEM GCSE, 3 = chose at least one STEM A(S)-Level, and 4 = chose to pursue a STEM Bachelor’s degree or higher. Box plots indicate median, interquartile range and min/max values. Asterisks indicate significant differences in spatial ability mean between groups at p < 0.05 level.

### Object manipulation abilities provide the most accurate prediction of STEM outcomes

When investigating potential differences in the associations across the three broad domains of spatial ability for different outcomes, we found limited evidence for differential predictions (Figure 3); the pattern of associations across the different domains was consistent across all outcomes. Notably, however, the associations for the object manipulation factor were slightly—but systematically—larger than those observed for the other three factors.

**Figure 3.**
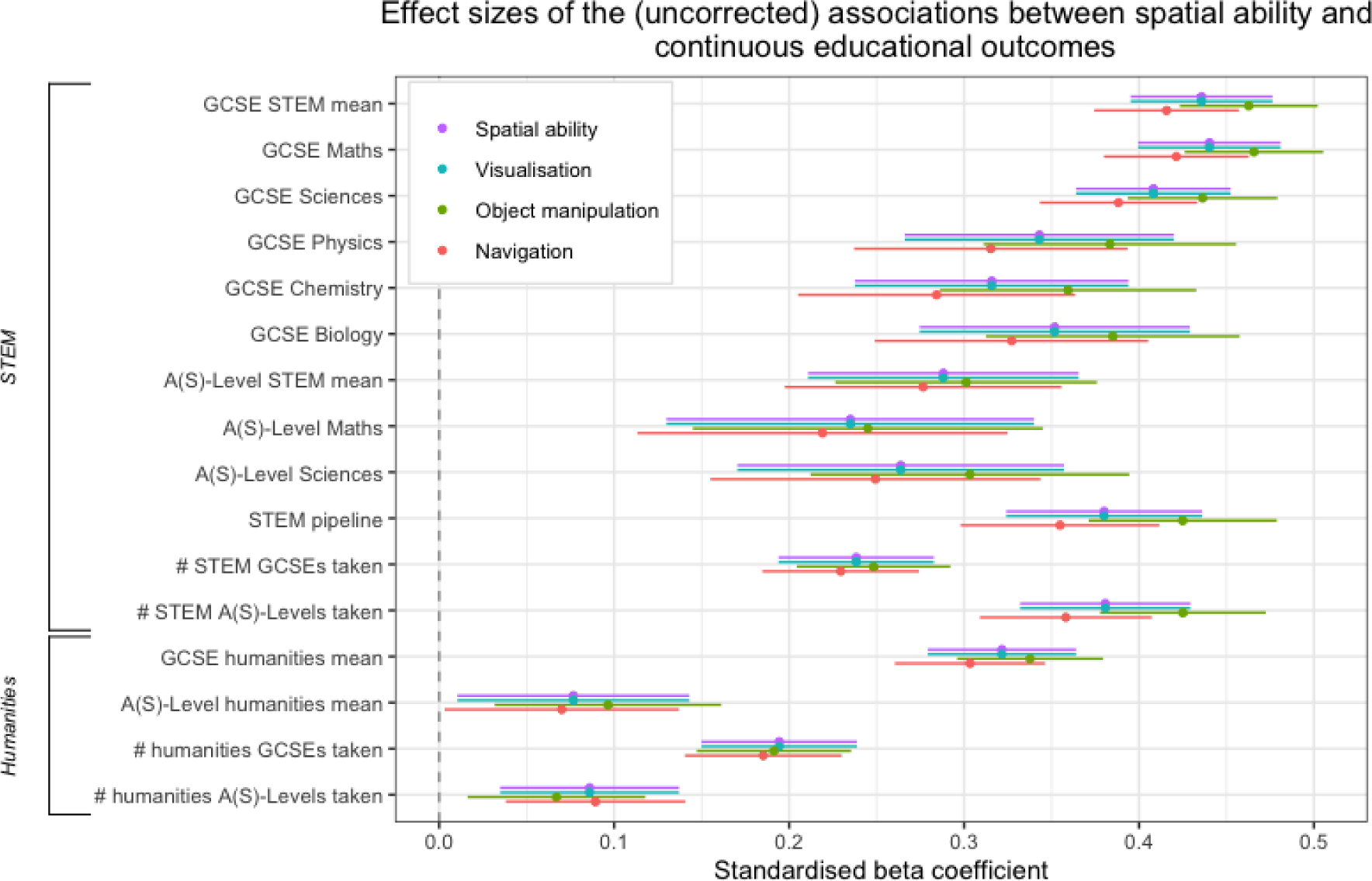
Standardized beta coefficients from linear regression analyses examining the uncorrected associations between spatial ability (and its domains) and two selected educational outcomes: mean GCSE grade in STEM subjects and remaining in the STEM pipeline for longer. Error bars represent 95% confidence intervals.

We, therefore, ran post-hoc analyses associating all 16 individual spatial ability items with two core outcomes that emerged from our previous analyses: *GCSE STEM mean grade* and *STEM pipeline*. We found that tests of object manipulation ability consistently resulted in greater associations with STEM outcomes than individual visualization or navigation ability tests. As shown in Figure 4, the most predictive tests were *2D drawing* (β=0.39. 95% CI [0.35, 0.44], *R*^2^ = 0.19) and *paper folding* (β=0.39. 95% CI [0.33, 0.45], *R*^2^ = 0.14), while the least predictive were scanning (β=0.06. 95% CI [-0.01, 0.13], *R*^2^ = 0.00) and map reading (β=0.08. 95% CI [0, 0.16], *R*^2^ = 0.01). Full model statistics are shown in SI Appendix, Table S9.

**Figure 4.**
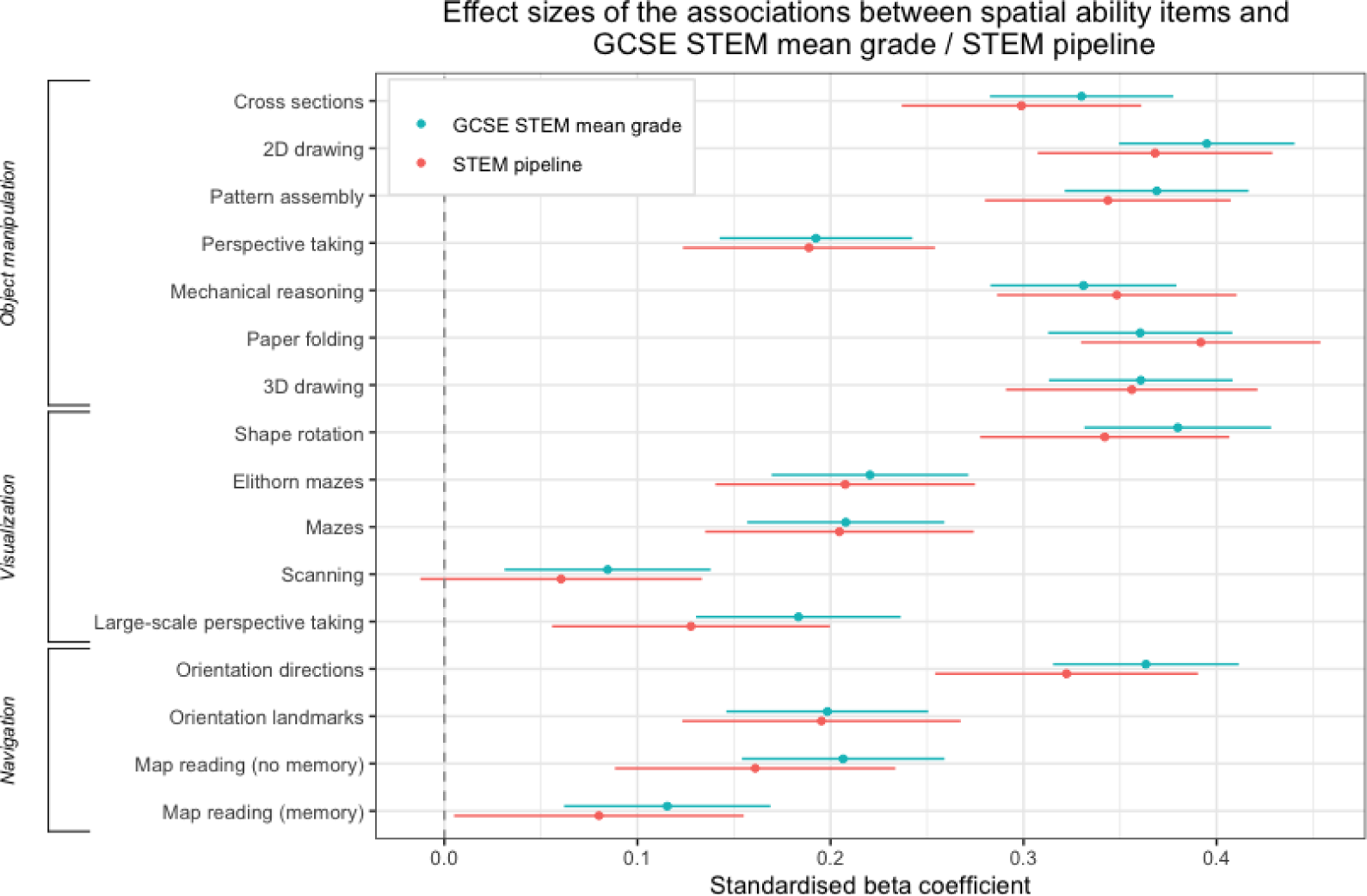
Effect sizes of the associations between individual spatial ability items and STEM GCSE mean grade and STEM pipeline. Error bars represent 95% confidence intervals.

### The link between spatial ability and STEM outcomes is largely accounted for by genetic factors

We conducted a series of ACE twin models (see Methods) to examine the extent to which genetic and environmental factors accounted for the associations between spatial ability and STEM outcomes.

Univariate analyses showed that, consistent with previous literature, all cognitive and educational variables were substantially heritable (*h*^2^ = 0.36 to 0.72). Spatial ability factors and some educational outcomes showed no evidence for shared environmental influence, while the remaining outcomes showed modest shared and unique environmental influences. Full univariate statistics can be found in SI Appendix, Table S11.

Then, we conducted bivariate correlated factors twin models to explore the shared etiology underpinning the phenotypic associations between spatial ability and educational outcomes. Due to the lack of domain specificity observed in our phenotypic analyses, we limited these analyses to our general spatial ability factor. We observed moderate to strong genetic correlations across almost all pairings of spatial ability and STEM educational outcomes (*r*A = 0.15 to 0.71) and, accordingly, high bivariate heritability estimates (bivariate *h*^2^ = 0.62 to 0.86), indicating that shared genetic factors accounted for the largest proportion of the phenotypic correlation between the two variables, between 62 and 86%. Moreover, these estimates were substantially higher for spatial ability-STEM pairings than for spatial ability-humanities pairings (*r*A = 0.04 to 0.40; bivariate *h*^2^ = 0.26 to 0.65), indicating that genetic overlap with spatial abilities was greater for most STEM subjects than humanities subjects.

Figure 5 depicts the genetic overlap between the general factor of spatial ability and STEM and humanities outcomes. Results for the shared environment were more varied, with some outcomes (for both STEM and humanities) showing substantial or even complete overlap. In contrast, non-shared environmental overlap was zero or near-zero for all outcomes (SI Figures S6-S7). Visualizations for these ACE components are displayed in SI Appendix, Figures S8 and S9, while full model statistics are presented in SI Appendix, Table S12.

**Figure 5.**
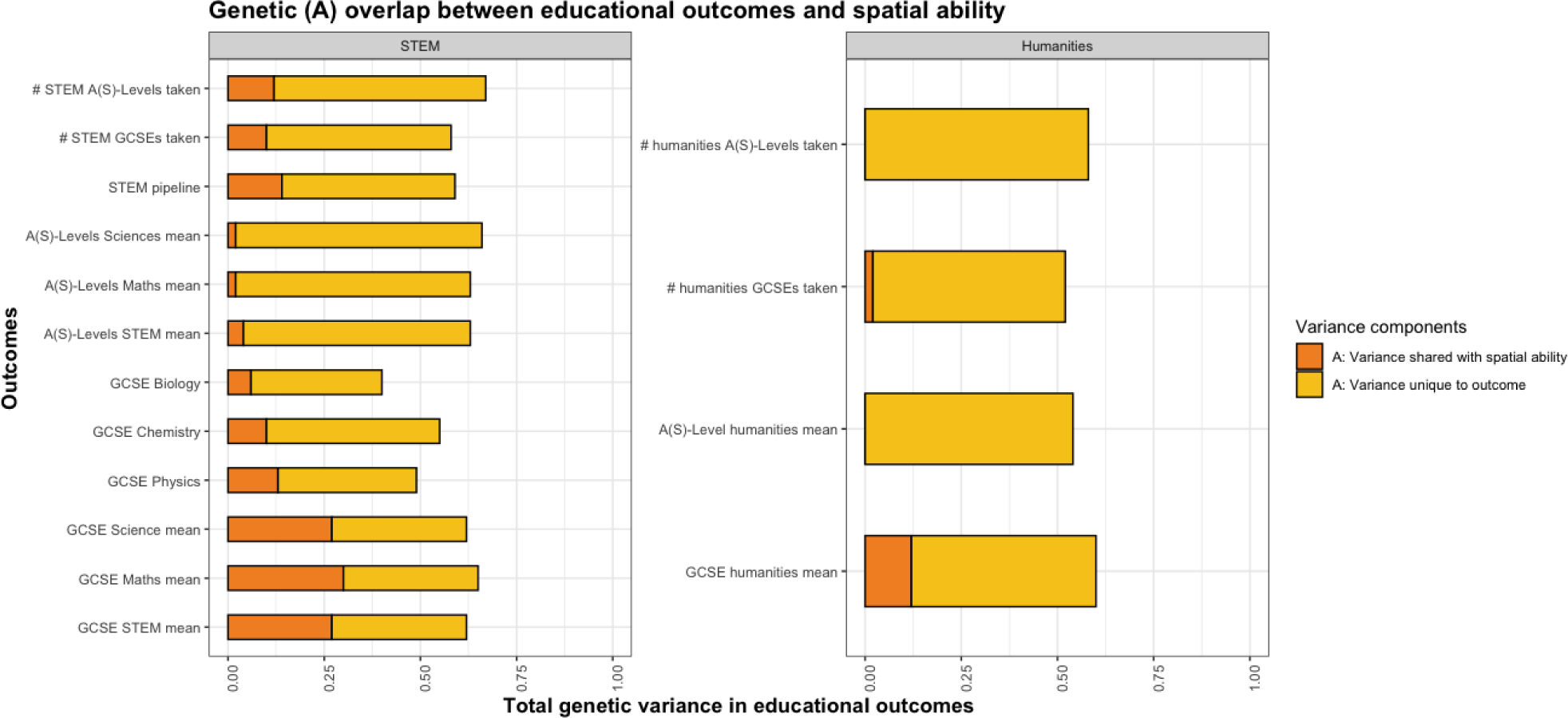
Standardized squared path estimates from bivariate Cholesky decompositions examining the etiological overlap between spatial ability and educational outcomes (only the genetic component is shown here; for the shared environmental and unique environmental components, refer to SI Appendix, Figures S6-S7). The total length of each bar represents the total genetic contribution to variance in the outcome (i.e., its heritability), which is decomposed into that unique to the outcome (in yellow) and that shared with spatial ability (in orange).

Finally, we conducted trivariate Cholesky decompositions to explore this shared aetiology after accounting for verbal ability or *g*. Mirroring our phenotypic analyses, we observed greater shared genetic variance for *g*-corrected models compared to verbal-corrected models. In other words, the specific shared genetic influence between spatial ability and educational outcomes were lower when accounting for *g* than when accounting for verbal ability. Visualizations for the general factor of spatial ability and educational outcome models accounting for verbal ability and *g* are displayed in SI Appendix, Figure S8 and S9, respectively, while full model statistics are presented in SI Appendix, Table S13.

## Discussion

Leveraging a unique battery of spatial ability tests across multiple domains, we explored three research questions. First, does spatial ability consistently predict STEM success above and beyond other cognitive abilities? Second, which domains of spatial ability, if any, are more predictive of STEM success? And third, to what extent do shared genetic and environmental factors account for the specific associations between spatial ability and STEM success?

Our results show that spatial ability is highly predictive of STEM success. These associations remained significant even after correcting for verbal ability or *g*, with the largest associations observed for STEM engagement rather than achievement variables. At a more granular level, object manipulation is a more accurate predictor than other spatial domains for almost all outcomes. At the etiological level, shared genetic factors consistently account for the largest proportion of the phenotypic associations between spatial ability and STEM success, though much of this genetic variance is also shared with general cognitive ability.

### The associations between spatial ability and STEM success persist after correcting for other cognitive abilities

We first examined the direct association between spatial ability (and its domains) and STEM success without controlling for other cognitive abilities. As expected, we found that spatial ability was highly predictive of STEM outcomes and to a much larger degree than humanities outcomes. Across all spatial ability domains, the largest direct predictions were observed for *GCSE Mathematics* and *GCSE STEM mean grades* (computed as the average of GCSE Mathematics, Statistics, Technology and all science subject grades (i.e., Statistics, Information and Communication Technology (ICT), Physics, Chemistry and Biology)). In comparison, the prediction for *GCSE humanities mean grade* (computed as the average of GCSE English Language, English Literature, all language subjects, and all other humanities subject grades (i.e., English, French, Spanish, German and History)) was roughly half of the *GCSE STEM mean grade*.

We also found that predictions for individual science subject grades (e.g., *GCSE Biology grade*) were much lower. This finding is likely explained by smaller sample sizes and reduced variance. At the time GCSE examinations were taken in our sample, only individuals pursuing the most advanced science qualifications were awarded individual subject grade GCSEs (others received either a “Core Science” GCSE or an “Additional Science” GCSE). Thus, individuals with data for *GCSE Biology grade, GCSE Chemistry grade* and *GCSE Physics grade* comprise only a small portion of the overall sample. Moreover, ostensible talent-based self-selection in this subsample would likely have reduced variance in predictor and outcome variables. Both explanations also apply to the relatively low A-Level outcome predictions, given that choosing to pursue A-Levels is non-compulsory.

Next, we examined how the associations between spatial ability and STEM outcomes persisted after accounting for verbal ability or *g*. We found that many predictions across the four spatial ability factors, though attenuated in effect size, remained highly significant for STEM outcomes but not humanities outcomes. This suggests that spatial ability is associated with humanities outcomes almost entirely due to the former’s shared variance with other cognitive abilities, while some unique variance accounts for its associations with STEM outcomes. Moreover, the attenuation in predictions was greater when correcting for *g* than for verbal ability, reflecting that *g* is perhaps an overly stringent control.

Most notable amongst our corrected analyses was the finding that the highest predictions across all spatial ability factors were for variables related to STEM engagement rather than achievement. The highest was for our *STEM pipeline* variable, a 4-level ordinal variable indicating how far an individual proceeds in STEM education, followed closely by the binary choice variables that were used to compute this variable (*chose a non-compulsory STEM GCSE*, *chose a STEM A(S)-Level*, and *chose a STEM degree*). Considering the relatively low corrected associations for STEM *achievement* variables, spatial ability may play a particularly important predictive role in STEM *interest* and *engagement*. This has profound implications for improving overall STEM outcomes, given that persistence and retention are necessary antecedents to achievement (20). However, we acknowledge that choice, persistence and retention are not independent of performance.

### Object manipulation skills are the strongest predictor of STEM success

The second question we aimed to explore in our phenotypic analyses was the extent to which we would observe differential predictions between our three domains. Our deep phenotyping allowed us to explore whether certain aspects of spatial ability were more predictive than others. we observed that object manipulation systematically explained more variance in STEM outcomes than the other factors, although the differences were only significant when looking at individual tests. In fact, when we examined these associations at the level of each spatial ability test, we found that several of the most predictive items that comprised the object manipulation factor (e.g., “2D drawing”, “pattern assembly”, and “paper folding”) *were* significantly more highly associated with STEM outcomes than the items that comprised the navigation and visualization factors. This suggests that honing in on developing these specific skills may have particular utility for prediction and intervention.

### Genetic and environmental associations

Finally, at the etiological level, we explored the extent to which shared latent genetic (A), shared environmental (C), and unique environmental factors (E) contributed to the phenotypic correlations between spatial ability and educational outcomes, both before and after accounting for shared etiological variance with *g* and verbal ability. We first conducted univariate twin models on all of our cognitive and educational outcome measures, observing, as expected, high heritability estimates for all of them (i.e., phenotypic variance in the trait was largely accounted for by genetic variance). We then conducted bivariate twin models pairing each educational outcome with spatial ability to decompose this genetic variance into that unique to the outcome and that shared with spatial ability. Therein, as seen in Fig. 5, we observed a higher degree of genetic overlap between spatial ability and STEM outcomes compared to spatial ability and humanities outcomes. However, when we added *g* to the model (i.e., extending it from a bivariate to a trivariate twin model), we found that the majority of the shared genetic variance between spatial ability and the outcome was already accounted for by genetic variance shared with *g*. Much like our findings from our phenotypic analyses, this attenuation in unique genetic variance contributed by spatial ability was less pronounced when we controlled for verbal ability rather than g. Given that previous work on our measure of spatial ability in the same sample found that 45% of its genetic influence was independent of genetic influence on g (21), the near-total genetic overlap with g in our trivariate models was somewhat unexpected, though it can be reconciled by considering that the remaining 55% of its genetic influence (i.e., the portion that is shared with g) may be the portion contributing to its phenotypic association with educational outcomes.

## Limitations

The present study has several limitations. First, our correlational analyses prohibit causal interpretations of spatial ability’s effects on STEM success. This is compounded by the fact that our spatial ability data was collected in our sample after GCSEs were taken. As such, our findings do not preclude the possibility that early STEM success may also influence the development of stronger spatial ability skills, especially when considering extant literature on the reciprocal relationship between general intelligence and educational achievement across development (22, 23).

Future work can address this weakness by employing a cross-lagged model on multiple measurements of both spatial ability and STEM achievement across development.

As previously discussed, our work is further limited by the variance and sample size restriction for individual STEM discipline data. As a result, we could not detect potential differential predictions for different STEM subjects (e.g., spatial ability being more predictive of, say, Physics achievement than Biology achievement). Given that extant literature has uncovered considerable differences in spatial-mathematics- verbal cognitive profiles between gifted degree holders in life science STEM fields (verbal > spatial) compared to physical science and engineering STEM degree holders (spatial > verbal) (24), future work should aim to corroborate this level of granularity in prediction.

Finally, regarding our genetic analyses, the usual limitations of the twin model apply. These include the lack of generalization as we only include British sample; and the potential for non-random mating, which can bias estimates of heritability in both directions. These limitations are described in detail elsewhere (25–27).

## Implications

The implications of our findings fall into two categories: intervention (i.e., improving outcomes, particularly in disadvantaged groups) and identification (i.e., identifying and nurturing highly gifted individuals primarily for human capital development).

With regards to intervention, several recent studies and meta-analyses have suggested that spatial ability, in contrast to *g*, appears to be malleable to long-term improvement when subject to early childhood training and activity (28–30). Given our finding that spatial ability—and, in particular, object manipulation ability—appears to have particular utility as a specific predictor of STEM interest and choice rather than achievement, policy makers looking to increase engagement and retention in the so-called STEM pipeline may find it particularly prudent to focus on intervening in the development of the interest and choice of these skills at a young age. It is important to note here that our finding of high heritability does not preclude the possibility of intervention. Heritability is a statistic derived from decomposing the observed variance in a given sample under given environmental conditions—not a static measure of immutability or determinism. Regarding identification, spatial ability may be a useful indicator of potential STEM talent in educational and professional settings. Whether it be schools looking to funnel precocious youth down advanced STEM educational pathways or businesses looking to increase productive output, tests of spatial ability may be beneficial for their predictive value in addition to that provided by general tests of cognitive ability (i.e., by realizing its extra predictive ability above and beyond *g*).

## Conclusions

In conclusion, the present study investigated the specific predictive utility of multiple domains of spatial ability on STEM success and the etiological underpinnings of this relationship. We found that spatial ability (e.g., object manipulation ability) was highly associated with STEM outcomes; these associations persisted even after correcting for other cognitive abilities, particularly for outcomes related to continued engagement in STEM education, highlighting the importance of interest and choice of students; and that shared genetic factors largely drove these associations. Our work underscores the distinctive nature of spatial ability and demonstrates its utility as a specific predictor of STEM educational outcomes, particularly regarding interest and choice.

## Methods and Materials

### Participants

Participants were part of the Twins Early Development Study (TEDS), a longitudinal study of more than 10,000 twin pairs representative of the UK population born in England and Wales between 1994 and 1996. The TEDS sample has been assessed over a dozen times from infancy through early adulthood on a wide range of measures. See Rimfeld et al. (2019) (31) for more details on the TEDS sample. The present study focuses on a subsample of TEDS twins who completed two spatial ability batteries at the ages of 19-22. For phenotypic analyses (i.e., linear and logistic regressions), one twin was randomly selected from each pair to account for relatedness, while full twin pairs were used for our genetic analyses. The maximum N was 1,968 individuals for phenotypic analyses and 3,936 individuals for genetic analyses. See SI Appendix, Tables S1-S11 for full descriptive statistics and model statistics for each variable and pairwise comparison. All individuals with major medical, genetic or neurodevelopmental disorders were excluded from the dataset. These included twins with ASD, cerebral palsy, Downs syndrome, chromosomal or single-gene disorders, organic brain problems, profound deafness and developmental delay.

### Measures

#### Spatial ability

Our spatial ability measures included 16 tests across two online batteries administered when our sample was aged 19-22. The first battery, called “King’s Challenge”, consisted of 10 tests spanning a range of object manipulation and visualization tasks: (1) a mazes task (searching for a way through a 2D maze in a speeded task); (2) 2D drawing (sketching a 2D layout of a 3D object from a specified viewpoint); (3) Elithorn mazes (joining together as many dots as possible from an array); (4) pattern assembly (visually combining pieces of objects to make a whole); (5) mechanical reasoning (multiple choice naïve physics questions); (6) paper folding (visualizing where the holes are situated after a piece of paper is folded and a hole is punched through it); (7) 3D drawing (sketching a 3D drawing from a 2D diagram); (8) mental rotation (mentally rotating objects); (9) perspective-taking (visualizing objects from a different perspective), and (10) cross-sections (visualizing cross-sections of objects). More details on the development of this battery and its tests are available elsewhere (15).

The second battery, “Spatial Spy,” consisted of 6 spatial tests within a gamified, virtual environment. Completed on a web browser, participants were asked to solve a mystery by collecting clues while navigating a virtual environment. Four abilities not traditionally assessed by object-oriented psychometric tests were measured in this battery: (1) navigating when reading a map; (2) navigating based on a previously memorized map or route; (3) navigating following directions (e.g., cardinal points), and (4) navigating using reference landmarks. Additionally, two traditional psychometric tests were adapted to assess spatial ability in a large-scale environment: perspective-taking and scanning. More details on this battery are outlined elsewhere (16).

Descriptive statistics for all items are displayed in SI Appendix, Table S1.

#### General cognitive ability (g) and verbal ability

Our measures for our cognitive ability controls spanned 7 measurements across development at ages 7, 9, 10, 12, 14, 16, and 21. Measurements varied in administration format (e.g., phone, booklet, online) and content, but systematically included verbal and non-verbal items from which verbal, non-verbal and general (i.e., g) mean composites were computed at each age. For comprehensive details on each measure and the derivation of composites, refer to the TEDS data dictionary here: https://www.teds.ac.uk/datadictionary/home.htm. Descriptive statistics for each measure are displayed in SI Appendix, Table S2.

#### Educational outcome measures

We examined a total of 46 available educational outcome measures (6 binary measures and 40 continuous measures), data for which were collected in our sample across development at ages 16, 18, and 21 (see https://www.teds.ac.uk/datadictionary/home.htm for details).

Our binary outcomes were comprised of yes (1) or no (0) variables relating to choice of non-compulsory STEM and humanities GCSEs, A-Levels, and degree.

Descriptive statistics for all outcome variables are displayed in SI Appendix, Tables S5-S7. Our continuous measures were comprised of individual grade scores, composite grade scores, the number of STEM/humanities GCSEs/A-Levels taken, and a “STEM pipeline” variable which is described below. Composite scores were computed as the mean of pre-existing composites in the TEDS data dictionary (e.g., “GCSE STEM mean” was derived from pre-existing variables for mathematics GCSE mean, science GCSE mean, and technology GCSE mean), while variables for number of STEM/humanities GCSEs and A-Levels taken were computed by summing the total number of individual subjects taken. The STEM pipeline was computed as a 4-level ordinal variable where one’s score represents the termination of their STEM education: 1 indicates that the individual did not choose any non-compulsory STEM GCSEs, 2 indicates that the individual chose at least one non-compulsory STEM GCSE, 3 indicates that the individual chose at least one STEM A-Level, and 4 indicates that the individual chose to pursue a STEM Bachelor’s degree or higher. To ensure that our pipeline variable accurately captured retention and drop-out as opposed to study attrition, we restricted our sample to only those individuals with available data at all 4 stages of the pipeline. All continuous measures were age- and sex-regressed prior to further analyses.

### Analyses

#### Confirmatory factor analysis for spatial ability, *g*, and verbal ability

Using the lavaan package in R (32), we first conducted confirmatory factor analysis (19) on our measures of spatial ability, *g* and verbal ability before extracting factor scores for each.

For spatial ability, we aimed to reproduce the best-fitting factor structure previously reported by Malanchini et al. (2020) in the same dataset, specifying a second-order model in which the three domains of object manipulation, navigation, and visualization load onto a higher-order spatial ability factor. Full information maximum likelihood estimation was used to account for missingness. This model provided a good fit to the data (χ2=649.319 (102), p<0.001, CFI=0.925, TLI=0.912, RMSEA=0.052, SMRS=0.063), revealing factor loadings on the general spatial ability factor of 0.86, 0.91 and 1.00 for object manipulation, navigation, and visualization, respectively. Full CFA model statistics for spatial ability (as well as for *g* and verbal ability, described below) are presented in SI Appendix, Figures S1-S3.

We then extracted factor scores for object manipulation, navigation, visualization, and the general factor of spatial ability from this model to use as independent variables in subsequent regression and twin analyses. All factor scores showed roughly normal distributions with acceptable values for skew (<+/-2) and kurtosis (<+/-7). All variables were age- and sex-regressed prior to factor analysis. The factor scores were standardized again prior to our phenotypic and genetic analyses.

Full descriptive statistics for our spatial ability factor scores (as well as for *g* and verbal ability) are shown in SI Appendix, Table S4.

For *g*, we specified a unifactorial model in which measures at the 7 ages loaded onto a single *g* factor. This model provided an excellent fit for the data (χ2=112.285 (14), p<0.001, CFI=0.963, TLI=0.944, RMSEA=0.070, SMRS=0.036), with factor loadings ranging from 0.60 to 0.84. Factor scores extracted from the *g* factor were normally distributed with acceptable values for skew and kurtosis. Again, variables were age- and sex-regressed prior to factor analysis, and the factor scores were standardized.

For verbal ability, we similarly specified a unifactorial model for verbal ability across the 7 ages. This model also provided an excellent fit to the data (χ2= 75.195 (14), p<0.001, CFI=0.97, TLI=0.954, RMSEA=0.055, SMRS=0.034), with factor loadings ranging from 0.60 to 0.80. Factor scores extracted from this verbal ability factor were normally distributed with acceptable values for skew (<+/-2) and kurtosis (<+/-7). As above, variables were age- and sex-regressed prior to factor analysis, and the factor scores were standardized. Correlational analyses, visualized in SI Appendix, Figure S4, confirmed strong phenotypic associations between all four factors of spatial ability, *g* and verbal ability.

#### Regression analyses

We first regressed our four spatial ability predictors (i.e., navigation, visualization, object manipulation, and the general factor of spatial ability) on *g* and verbal ability to create 12 total predictor variables (e.g., navigation, navigation corrected for g, and navigation corrected for verbal ability). We then conducted 480 linear regressions (i.e., all permutations for our 40 continuous outcomes on the 12 predictors) and adjusted for multiple comparisons using Benjamini-Hochberg correction. We then conducted 72 logistic regressions (i.e., all permutations for our 6 binary outcomes on the 12 predictors), again adjusting for multiple comparisons using Benjamini-Hochberg correction. We calculated Nagelkerke *R*^2^ values for our logistic regression models. Full model statistics are presented in SI Appendix, Table S8 and Table S9.

#### Genetic analyses

Using the OpenMX package in R (33), we conducted univariate, bivariate and trivariate twin models to examine *a)* the etiology underlying our cognitive and educational measures, *b)* the shared etiology underlying the phenotypic associations between spatial ability and educational outcomes, and *c)* the unique shared etiology underlying the phenotypic associations between spatial ability and educational outcomes after accounting for shared etiology with *g* or verbal ability. Full model statistics and visualizations are presented in SI Appendix, Tables S11-S13; Figure 5; and SI Appendix, Figures S6-S9.

#### Univariate twin model

The classical univariate ACE twin method is a design that capitalizes on the difference in genetic relatedness between MZ twins, who share 100% of their DNA, and DZ twins, who share on average 50% of their segregating genes, to decompose individual differences in an outcome into genetic and environmental sources of variance. By assuming that the shared family environment of both MZ and DZ twins are equal, a difference in intraclass correlations in trait outcome (i.e., a greater MZ correlation than DZ correlation) can be inferred as the effect of additive genetic influence (commonly denoted as A, or heritability) on the variance in the trait.

The environmental factors which make twins more similar is inferred as the effect of the shared environment (or C). The unique or non-shared environment (or E) effects, which are the individual-specific environmental influences that make family members different from each other (34).

#### Bivariate and trivariate twin models

The multivariate twin method extends the logic of the univariate twin method by decomposing the covariance between two or more traits into A, C and E components, thus providing information on the extent to which a phenotypic correlation between two or more traits is mediated genetically and environmentally. This is done by comparing the cross-trait cross-twin correlations between MZ and DZ twins, with a higher MZ correlation implying shared additive genetic covariance between the traits. The bivariate model produces a statistic called genetic correlation (R_A_), which indexes pleiotropy by denoting how genetic effects on one phenotype correlate with genetic effects on the other, independent of the heritability of the traits. When this genetic correlation is weighted by the heritability of each phenotype, an estimate of bivariate heritability (the amount of covariance accounted for by genetic factors) is produced.

Cholesky decomposition is an alternate solution to the correlated ACE model that places a special distinction on the ordering of variables such that variance can be decomposed into shared and independent ACE effects. By extending the bivariate design to a trivariate design with *g* or verbal ability ordered as the first variable, a spatial ability factor second, and the outcome last, we can investigate the independent variance accounted for by the spatial ability factor on the outcome.

## Acknowledgements

We gratefully acknowledge the ongoing contribution of the participants in the TEDS and their families. TEDS has been supported by a programme grant to R.P. from the UK Medical Research Council (MR/M021475/1 and previously G0901245), with additional support from the US National Institutes of Health (AG046938). The research leading to these results has also received funding from the European Research Council under the European Union’s Seventh Framework Programme (FP7/2007-2013)/ grant agreement n° 602768. Q.Z. is funded by the QMUL-Chinese Scholarship Council joint PhD Scholarship. M.M. is supported by a starting grant from the School of Biological and Behavioural Sciences at Queen Mary University of London and a Jacobs Foundation Research Fellowship.

## Author Contributions

Conceptualization: Margherita Malanchini, Kaili Rimfeld, Ziye Wang, Quan Zhou Analysis: Ziye Wang, Quan Zhou

Writing: Margherita Malanchini, Ziye Wang, Quan Zhou

Editing, feedback, advice: Kaili Rimfeld, Andrea Allegrini, Robert Plomin

## References

1. OECD, Education at a Glance 2022: OECD Indicators (OECD, 2022).

2. A. Chevalier, Subject choice and earnings of UK graduates. Economics of Education Review 30, 1187–1201 (2011).

3. H. Rindermann, J. Thompson, Cognitive Capitalism: The Effect of Cognitive Ability on Wealth, as Mediated Through Scientific Achievement and Economic Freedom. Psychol Sci 22, 754–763 (2011).

4. H. B. Gonzalez, J. J. Kuenzi, Science, Technology, Engineering, and Mathematics (STEM) Education: A Primer.

5. T. J. Kennedy, M. R. L. Odell, Engaging Students in STEM Education. Science Education International 25, 246–258 (2014).

6. J. Wai, D. Lubinski, C. P. Benbow, Spatial ability for STEM domains: Aligning over 50 years of cumulative psychological knowledge solidifies its importance. Journal of Educational Psychology 101, 817–835 (2009).

7. R. Webb, D. Lubinski, C. Benbow, Spatial ability: A neglected dimension in talent searches for intellectually precocious youth. Journal of Educational Psychology 99, 397–420 (2007).

8. D. Lubinski, Spatial ability and STEM: A sleeping giant for talent identification and development. Personality and Individual Differences 49, 344–351 (2010).

9. D. F. Lohman, “Spatial Ability and g” in Human Abilities, (Psychology Press, 1996).

10. J. B. Carroll, Human cognitive abilities: A survey of factor-analytic studies (Cambridge University Press, 1993).

11. A. Binet, Th. Simon, Méthodes nouvelles pour le diagnostic du niveau intellectuel des anormaux. psy 11, 191–244 (1904).

12. R. E. Snow, D. F. Lohman, “Implications of cognitive psychology for educational measurement” in Educational Measurement*, 3rd Ed*, The American Council on Education/Macmillan series on higher education., (American Council on Education, 1989), pp. 263–331.

13. D. L. Shea, D. Lubinski, C. P. Benbow, Importance of assessing spatial ability in intellectually talented young adolescents: A 20-year longitudinal study. Journal of Educational Psychology 93, 604–614 (2001).

14. L. G. Humphreys, D. Lubinski, G. Yao, Utility of predicting group membership and the role of spatial visualization in becoming an engineer, physical scientist, or artist. Journal of Applied Psychology 78, 250–261 (1993).

15. K. Rimfeld, et al., Phenotypic and genetic evidence for a unifactorial structure of spatial abilities. Proceedings of the National Academy of Sciences 114, 2777–2782 (2017).

16. M. Malanchini, et al., Evidence for a unitary structure of spatial cognition beyond general intelligence. *npj Sci*. Learn. 5, 1–13 (2020).

17. M. J. King, D. P. Katz, L. A. Thompson, B. N. Macnamara, Genetic and environmental influences on spatial reasoning: A meta-analysis of twin studies. Intelligence 73, 65–77 (2019).

18. F. Procopio, et al., The genetics of specific cognitive abilities. Intelligence 95, 101689 (2022).

19. T. A. Brown, Confirmatory Factor Analysis for Applied Research*, Second Edition* (Guilford Publications, 2015).

20. K. P. Harden, et al., Genetic associations with mathematics tracking and persistence in secondary school. NPJ Sci Learn 5, 1 (2020).

21. M. Malanchini, et al., Evidence for a unitary structure of spatial cognition beyond general intelligence. *npj Sci*. Learn. 5, 1–13 (2020).

22. R. Cowan, J. Hurry, E. Midouhas, The relationship between learning mathematics and general cognitive ability in primary school. British Journal of Developmental Psychology 36, 277–284 (2018).

23. K. Cain, J. Oakhill, Matthew Effects in Young Readers: Reading Comprehension and Reading Experience Aid Vocabulary Development. J Learn Disabil 44, 431–443 (2011).

24. D. Lubinski, Neglected aspects and truncated appraisals in vocational counseling: Interpreting the interest–efficacy association from a broader perspective: Comment on Armstrong and Vogel (2009). Journal of Counseling Psychology 57, 226–238 (2010).

25. F. V. Rijsdijk, P. C. Sham, Analytic approaches to twin data using structural equation models. Brief Bioinform 3, 119–133 (2002).

26. D. Boomsma, A. Busjahn, L. Peltonen, Classical twin studies and beyond. Nat Rev Genet 3, 872–882 (2002).

27. J. Felson, What can we learn from twin studies? A comprehensive evaluation of the equal environments assumption. Soc Sci Res 43, 184–199 (2014).

28. D. H. Uttal, et al., The malleability of spatial skills: A meta-analysis of training studies. Psychological Bulletin 139, 352–402 (2013).

29. W. Yang, H. Liu, N. Chen, P. Xu, X. Lin, Is Early Spatial Skills Training Effective? A Meta-Analysis. Frontiers in Psychology 11 (2020).

30. E. G. Peterson, A. B. Weinberger, D. H. Uttal, B. Kolvoord, A. E. Green, Spatial activity participation in childhood and adolescence: consistency and relations to spatial thinking in adolescence. *Cogn*. Research 5, 43 (2020).

31. K. Rimfeld, et al., Twins Early Development Study: A Genetically Sensitive Investigation into Behavioral and Cognitive Development from Infancy to Emerging Adulthood. Twin Res Hum Genet 22, 508–513 (2019).

32. Y. Rosseel, lavaan: An R Package for Structural Equation Modeling. Journal of Statistical Software 48, 1–36 (2012).

33. S. M. Boker, et al., OpenMx: Extended Structural Equation Modelling. (2024). Deposited 17 August 2024.

34. W. Johnson, E. Turkheimer, I. I. Gottesman, T. J. Bouchard, Beyond Heritability: Twin Studies in Behavioral Research. Curr Dir Psychol Sci 18, 217–220 (2010).

