## Supplementary information for "Exploring the specific predictive ability of multiple domains of spatial ability on STEM educational outcomes"

**This PDF file includes:**

Figures S1 to S10 Tables S1 to S12

**Table S1.** Descriptive statistics for the 16 spatial ability tests, randomly selecting one twin from each pair. Outliers, defined by observations falling outside of 3 standard deviations of the mean, were removed from the present descriptive analyses and all subsequent analyses. The number of outliers for each measure ranged from 0 to 31, with an average of 10 per measure. All measures were standardized and residualized for age and sex by means of linear regression.

Descriptive statistics for spatial ability (King's Challenge & Spatial Spy) battery items

| vars | n | mean | sd | median | trimmed | mad | min | max | range | skew | kurtosis | se |
|---|---|---|---|---|---|---|---|---|---|---|---|---|
| King's Challenge: Cross sections | 1383 | 0.00 | 1.00 | 0.08 | 0.02 | 1.07 | −2.23 | 2.42 | 4.65 | −0.18 | −0.74 | 0.03 |
| King's Challenge: 2D drawing | 1378 | 0.00 | 1.00 | 0.15 | 0.08 | 0.96 | −3.04 | 1.63 | 4.66 | −0.68 | −0.02 | 0.03 |
| King's Challenge: Pattern assembly | 1325 | 0.00 | 1.00 | 0.20 | 0.05 | 1.00 | −2.31 | 2.47 | 4.78 | −0.41 | −0.64 | 0.03 |
| King's Challenge: Perspective taking | 1243 | 0.00 | 1.00 | −0.18 | −0.07 | 1.01 | −1.76 | 2.93 | 4.69 | 0.59 | −0.28 | 0.03 |
| King's Challenge: Mechanical reasoning | 1332 | 0.00 | 1.00 | 0.13 | 0.01 | 0.98 | −2.83 | 2.69 | 5.52 | −0.11 | −0.22 | 0.03 |
| King's Challenge: Paper folding | 1285 | 0.00 | 1.00 | 0.11 | 0.02 | 1.20 | −2.36 | 2.09 | 4.45 | −0.17 | −0.94 | 0.03 |
| King's Challenge: 3D drawing | 1222 | 0.00 | 1.00 | 0.00 | 0.00 | 1.22 | −2.04 | 2.39 | 4.43 | −0.01 | −0.99 | 0.03 |
| King's Challenge: Shape rotation | 1241 | 0.00 | 1.00 | 0.09 | 0.04 | 1.14 | −2.44 | 2.04 | 4.48 | −0.29 | −0.82 | 0.03 |
| King's Challenge: Elithorn mazes | 1143 | 0.00 | 1.00 | 0.19 | 0.11 | 0.85 | −3.63 | 1.93 | 5.56 | −1.09 | 1.30 | 0.03 |
| King's Challenge: Mazes | 1218 | 0.00 | 1.00 | 0.14 | 0.02 | 1.00 | −3.10 | 2.56 | 5.66 | −0.25 | −0.11 | 0.03 |
| Spatial Spy: Orientation direction | 1351 | 0.00 | 1.00 | 0.06 | 0.03 | 1.06 | −3.06 | 2.54 | 5.59 | −0.27 | −0.34 | 0.03 |
| Spatial Spy: Orientation landmarks | 1315 | 0.00 | 1.00 | 0.17 | 0.10 | 0.85 | −3.79 | 1.82 | 5.61 | −1.04 | 1.26 | 0.03 |
| Spatial Spy: Map reading (no memory) | 1277 | 0.00 | 1.00 | 0.24 | 0.09 | 0.71 | −3.61 | 1.67 | 5.28 | −0.96 | 1.05 | 0.03 |
| Spatial Spy: Map reading (memory) | 1257 | 0.00 | 1.00 | 0.17 | 0.12 | 0.80 | −3.71 | 1.65 | 5.36 | −1.21 | 1.55 | 0.03 |
| Spatial Spy: Large scale perspective taking | 1321 | 0.00 | 1.00 | 0.17 | 0.10 | 0.96 | −3.16 | 1.50 | 4.66 | −0.82 | 0.08 | 0.03 |
| Spatial Spy: Scanning | 1268 | 0.00 | 1.00 | 0.23 | 0.13 | 0.77 | −4.21 | 1.50 | 5.70 | −1.35 | 1.96 | 0.03 |

**Table S2.** Descriptive statistics for the 7 measures of *g* across development, randomly selecting one twin from each pair. All measures were standardized and residualized for age and sex by means of linear regression.

Descriptive statistics for g measures

| vars | n | mean | sd | median | trimmed | mad | min | max | range | skew | kurtosis | se |
|---|---|---|---|---|---|---|---|---|---|---|---|---|
| g (age 7) | 1186 | 0.00 | 1.00 | 0.05 | 0.02 | 0.95 | −4.77 | 4.67 | 9.44 | −0.25 | 0.83 | 0.03 |
| g (age 9) | 1131 | 0.00 | 1.00 | 0.16 | 0.07 | 1.00 | −3.34 | 2.21 | 5.55 | −0.63 | 0.04 | 0.03 |
| g (age 10) | 981 | 0.00 | 1.00 | 0.06 | 0.02 | 1.01 | −3.20 | 2.73 | 5.93 | −0.25 | 0.03 | 0.03 |
| g (age 12) | 984 | 0.00 | 1.00 | 0.08 | 0.04 | 0.98 | −3.79 | 2.71 | 6.50 | −0.43 | 0.20 | 0.03 |
| g (age 14) | 828 | 0.00 | 1.00 | 0.06 | 0.03 | 0.93 | −3.78 | 2.41 | 6.19 | −0.40 | 0.56 | 0.03 |
| g (age 16) | 1296 | 0.00 | 1.00 | −0.05 | −0.03 | 0.95 | −2.88 | 3.37 | 6.25 | 0.28 | 0.19 | 0.03 |
| g (age 21) | 728 | 0.00 | 1.00 | 0.15 | 0.06 | 1.02 | −3.47 | 2.08 | 5.54 | −0.52 | −0.14 | 0.04 |

**Table S3.** Descriptive statistics for the 7 measures of verbal ability across development, randomly selecting one twin from each pair. All measures were standardized and residualized for age and sex by means of linear regression.

Descriptive statistics for verbal ability measures

| vars | n | mean | sd | median | trimmed | mad | min | max | range | skew | kurtosis | se |
|---|---|---|---|---|---|---|---|---|---|---|---|---|
| Verbal ability (age 7) | 1189 | 0.00 | 1.00 | 0.03 | 0.01 | 0.96 | −3.36 | 5.70 | 9.06 | 0.06 | 0.94 | 0.03 |
| Verbal ability (age 9) | 1144 | 0.00 | 1.00 | 0.06 | 0.03 | 1.03 | −3.39 | 2.54 | 5.93 | −0.32 | −0.10 | 0.03 |
| Verbal ability (age 10) | 995 | 0.00 | 1.00 | 0.02 | 0.01 | 1.07 | −2.90 | 2.59 | 5.49 | −0.06 | −0.38 | 0.03 |
| Verbal ability (age 12) | 1019 | 0.00 | 1.00 | 0.11 | 0.05 | 1.02 | −3.15 | 2.27 | 5.42 | −0.42 | −0.23 | 0.03 |
| Verbal ability (age 14) | 906 | 0.00 | 1.00 | 0.11 | 0.08 | 0.86 | −4.08 | 2.09 | 6.17 | −0.82 | 0.99 | 0.03 |
| Verbal ability (age 16) | 1361 | 0.00 | 1.00 | −0.06 | −0.05 | 0.92 | −2.76 | 4.11 | 6.87 | 0.60 | 0.94 | 0.03 |
| Verbal ability (age 21) | 728 | 0.00 | 1.00 | 0.12 | 0.05 | 1.15 | −3.05 | 2.00 | 5.05 | −0.41 | −0.43 | 0.04 |

**Table S4.** Descriptive statistics for g, verbal ability, and the four spatial ability factors (uncorrected, corrected for g, and corrected for verbal ability). Variables were computed as factor scores extracted from the CFA models presented in Figs. S1-S3. Full information maximum likelihood modelling was used to account for missingness in the data. VA = verbal ability. Corrected predictors were derived by regressing the respective control variable (i.e., g or VA) on the spatial ability factor. All measures were standardized.

Descriptive statistics for cognitive predictors

| vars | n | mean | sd | median | trimmed | mad | min | max | range | skew | kurtosis | se |
|---|---|---|---|---|---|---|---|---|---|---|---|---|
| Object manipulation | 1968 | 0.00 | 1.00 | 0.08 | 0.04 | 1.02 | −3.34 | 2.93 | 6.27 | −0.31 | −0.33 | 0.02 |
| Object manipulation (g-corrected) | 1431 | 0.00 | 1.00 | 0.04 | 0.02 | 1.03 | −3.23 | 2.79 | 6.02 | −0.20 | −0.20 | 0.03 |
| Object manipulation (VA-corrected) | 1434 | 0.00 | 1.00 | 0.05 | 0.03 | 1.07 | −3.18 | 2.62 | 5.80 | −0.21 | −0.38 | 0.03 |
| Visualization | 1968 | 0.00 | 1.00 | 0.11 | 0.05 | 1.00 | −3.70 | 2.73 | 6.43 | −0.41 | −0.21 | 0.02 |
| Visualization (g-corrected) | 1431 | 0.00 | 1.00 | 0.08 | 0.03 | 1.01 | −3.40 | 2.64 | 6.04 | −0.30 | −0.16 | 0.03 |
| Visualization (VA-corrected) | 1434 | 0.00 | 1.00 | 0.08 | 0.04 | 1.04 | −3.37 | 2.63 | 5.99 | −0.31 | −0.26 | 0.03 |
| Navigation | 1968 | 0.00 | 1.00 | 0.09 | 0.05 | 1.00 | −3.93 | 2.50 | 6.43 | −0.44 | −0.05 | 0.02 |
| Navigation (g-corrected) | 1431 | 0.00 | 1.00 | 0.10 | 0.04 | 0.96 | −3.33 | 2.64 | 5.97 | −0.36 | 0.09 | 0.03 |
| Navigation (VA-corrected) | 1434 | 0.00 | 1.00 | 0.09 | 0.04 | 1.00 | −3.55 | 2.59 | 6.14 | −0.37 | −0.06 | 0.03 |
| Spatial ability | 1968 | 0.00 | 1.00 | 0.11 | 0.05 | 1.00 | −3.70 | 2.73 | 6.43 | −0.41 | −0.21 | 0.02 |
| Spatial ability (g-corrected) | 1431 | 0.00 | 1.00 | 0.08 | 0.03 | 1.01 | −3.40 | 2.64 | 6.04 | −0.30 | −0.16 | 0.03 |
| Spatial ability (VA-corrected) | 1434 | 0.00 | 1.00 | 0.08 | 0.04 | 1.04 | −3.37 | 2.63 | 5.99 | −0.31 | −0.26 | 0.03 |
| g | 1433 | 0.00 | 1.00 | 0.04 | 0.03 | 0.98 | −3.54 | 2.76 | 6.30 | −0.31 | 0.05 | 0.03 |
| Verbal ability | 1436 | 0.00 | 1.00 | 0.06 | 0.03 | 1.00 | −3.27 | 2.61 | 5.88 | −0.29 | −0.08 | 0.03 |

**Table S5.** Descriptive statistics for continuous GCSE outcome variables, randomly selecting one twin from each pair. All measures were standardized and residualized for age and sex by means of linear regression.

| vars | n | mean | sd | median | trimmed | mad | min | max | range | skew | kurtosis | se |
|---|---|---|---|---|---|---|---|---|---|---|---|---|
| GCSE STEM mean grade | 6471 | 0 | 1 | 0.02 | 0.04 | 1.05 | -3.89 | 1.74 | 5.63 | -0.45 | 0.17 | 0.01 |
| GCSE humanities mean grade | 6470 | 0 | 1 | 0.04 | 0.04 | 1.01 | -4.24 | 2.04 | 6.28 | -0.49 | 0.35 | 0.01 |
| GCSE STEM + humanities mean grade | 6486 | 0 | 1 | 0.02 | 0.04 | 1.04 | -4.27 | 1.97 | 6.25 | -0.45 | 0.22 | 0.01 |
| GCSE all subjects mean grade | 6486 | 0 | 1 | 0.03 | 0.04 | 1.05 | -4.11 | 1.96 | 6.07 | -0.44 | 0.16 | 0.01 |
| GCSE Core mean grade | 6480 | 0 | 1 | 0.00 | 0.04 | 1.07 | -4.10 | 1.82 | 5.92 | -0.45 | 0.22 | 0.01 |
| GCSE Maths mean grade | 6412 | 0 | 1 | 0.06 | 0.06 | 1.03 | -3.54 | 1.59 | 5.14 | -0.54 | 0.32 | 0.01 |
| GCSE English mean grade | 6450 | 0 | 1 | -0.01 | 0.04 | 1.04 | -4.30 | 2.00 | 6.30 | -0.43 | 0.29 | 0.01 |
| GCSE Science mean grade | 6002 | 0 | 1 | -0.01 | 0.04 | 1.14 | -3.92 | 1.57 | 5.50 | -0.38 | 0.01 | 0.01 |
| GCSE Technology mean grade | 3537 | 0 | 1 | 0.12 | 0.05 | 1.03 | -4.04 | 1.82 | 5.86 | -0.53 | 0.36 | 0.02 |
| GCSE Humanities mean grade | 6144 | 0 | 1 | 0.13 | 0.06 | 1.03 | -3.92 | 1.76 | 5.67 | -0.61 | 0.31 | 0.01 |
| GCSE Languages mean grade | 3870 | 0 | 1 | 0.06 | 0.03 | 1.06 | -3.52 | 1.71 | 5.24 | -0.33 | -0.20 | 0.02 |
| GCSE Vocational mean grade | 2579 | 0 | 1 | 0.08 | 0.03 | 1.12 | -3.86 | 1.78 | 5.65 | -0.42 | 0.03 | 0.02 |
| GCSE Maths grade | 6405 | 0 | 1 | 0.04 | 0.06 | 1.02 | -3.53 | 1.55 | 5.08 | -0.54 | 0.27 | 0.01 |
| GCSE Science core grade | 3376 | 0 | 1 | 0.28 | 0.02 | 1.14 | -3.77 | 2.00 | 5.77 | -0.28 | 0.53 | 0.02 |
| GCSE Stats grade | 719 | 0 | 1 | 0.04 | 0.02 | 1.22 | -4.10 | 1.73 | 5.84 | -0.33 | 0.14 | 0.04 |
| GCSE GCSE ICT grade | 1352 | 0 | 1 | -0.02 | 0.02 | 1.15 | -4.17 | 1.80 | 5.96 | -0.48 | 0.71 | 0.03 |
| GCSE Science Additional grade | 2712 | 0 | 1 | 0.13 | 0.00 | 1.12 | -3.88 | 1.97 | 5.85 | -0.13 | 0.05 | 0.02 |
| GCSE Physics grade | 2586 | 0 | 1 | 0.29 | 0.08 | 1.26 | -4.80 | 1.23 | 6.03 | -0.60 | -0.15 | 0.02 |
| GCSE Chemistry grade | 2592 | 0 | 1 | 0.32 | 0.08 | 1.21 | -4.64 | 1.22 | 5.86 | -0.61 | -0.09 | 0.02 |
| GCSE Biology grade | 2605 | 0 | 1 | 0.29 | 0.07 | 1.26 | -4.89 | 1.25 | 6.15 | -0.63 | 0.20 | 0.02 |
| GCSE English Language grade | 6399 | 0 | 1 | -0.02 | 0.02 | 1.14 | -4.17 | 1.93 | 6.11 | -0.32 | 0.12 | 0.01 |
| GCSE English Literature grade | 5702 | 0 | 1 | 0.03 | 0.03 | 1.06 | -4.48 | 1.88 | 6.36 | -0.38 | 0.10 | 0.01 |
| GCSE French grade | 2220 | 0 | 1 | -0.01 | 0.04 | 1.05 | -3.52 | 1.58 | 5.10 | -0.31 | -0.37 | 0.02 |
| GCSE History grade | 2847 | 0 | 1 | -0.05 | 0.09 | 1.01 | -3.62 | 1.34 | 4.96 | -0.74 | 0.20 | 0.02 |
| GCSE Spanish grade | 916 | 0 | 1 | -0.01 | 0.06 | 1.05 | -3.43 | 1.42 | 4.84 | -0.49 | -0.29 | 0.03 |
| GCSE German grade | 922 | 0 | 1 | 0.01 | 0.02 | 1.09 | -3.65 | 1.71 | 5.36 | -0.25 | -0.17 | 0.03 |
| Number of non-compulsory STEM GCSEs taken | 6514 | 0 | 1 | -0.04 | -0.02 | 1.17 | -1.63 | 2.50 | 4.13 | 0.19 | -1.02 | 0.01 |
| Total number of STEM GCSEs taken | 6514 | 0 | 1 | 0.27 | 0.00 | 1.00 | -2.94 | 2.87 | 5.81 | 0.03 | 0.56 | 0.01 |
| Number of non-compulsory humanities GCSEs taken | 6514 | 0 | 1 | 0.04 | 0.04 | 1.36 | -3.01 | 3.16 | 6.17 | -0.25 | -0.24 | 0.01 |
| Total number of humanities GCSEs taken | 6514 | 0 | 1 | -0.08 | -0.03 | 1.65 | -1.36 | 3.59 | 4.96 | 0.27 | -0.74 | 0.01 |

**Table S6.** Descriptive statistics for continuous A(S)-Level outcome variables and the STEM pipeline variable, randomly selecting one twin from each pair. All measures were standardized and residualized for age and sex by means of linear regression.

| vars | n | mean | sd | median | trimmed | mad | min | max | range | skew | kurtosis | se |
|---|---|---|---|---|---|---|---|---|---|---|---|---|
| A(S)-Level STEM mean grade | 2596 | 0 | 1 | 0.09 | 0.03 | 1.07 | -2.34 | 2.32 | 4.66 | -0.24 | -0.71 | 0.02 |
| A(S)-Level humanities mean grade | 3441 | 0 | 1 | 0.17 | 0.03 | 1.09 | -2.55 | 2.20 | 4.74 | -0.20 | -0.43 | 0.02 |
| A(S)-Level STEM + humanities mean grade | 3965 | 0 | 1 | 0.04 | 0.02 | 1.10 | -2.71 | 2.42 | 5.14 | -0.20 | -0.55 | 0.02 |
| A(S)-Level English mean grade | 1262 | 0 | 1 | 0.11 | -0.01 | 1.23 | -2.43 | 1.83 | 4.26 | -0.01 | -0.54 | 0.03 |
| A(S)-Level Maths mean grade | 1420 | 0 | 1 | -0.05 | 0.06 | 1.08 | -2.29 | 1.41 | 3.70 | -0.55 | -0.53 | 0.03 |
| A(S)-Level Science mean grade | 1829 | 0 | 1 | 0.21 | 0.03 | 1.07 | -1.95 | 1.75 | 3.70 | -0.22 | -0.92 | 0.02 |
| A(S)-Level Technology mean grade | 697 | 0 | 1 | 0.13 | 0.00 | 1.11 | -2.21 | 2.12 | 4.34 | -0.01 | -0.55 | 0.04 |
| A(S)-Level Humanities mean grade | 3215 | 0 | 1 | 0.15 | 0.03 | 1.11 | -2.48 | 2.06 | 4.54 | -0.23 | -0.43 | 0.02 |
| A(S)-Level Languages mean grade | 466 | 0 | 1 | 0.18 | 0.06 | 1.13 | -2.33 | 1.78 | 4.11 | -0.52 | -0.48 | 0.05 |
| A(S)-Level Vocational mean grade | 1031 | 0 | 1 | 0.21 | 0.01 | 1.15 | -2.24 | 2.16 | 4.40 | -0.11 | -0.55 | 0.03 |
| A(S)-Level overall mean grade | 4020 | 0 | 1 | 0.01 | 0.02 | 1.11 | -2.59 | 2.13 | 4.72 | -0.15 | -0.55 | 0.02 |
| A-Level overall mean grade | 3867 | 0 | 1 | 0.04 | 0.02 | 1.12 | -2.71 | 2.02 | 4.72 | -0.19 | -0.54 | 0.02 |
| Number of STEM A(S)-Levels taken | 6848 | 0 | 1 | -0.52 | -0.19 | 0.34 | -0.80 | 3.79 | 4.59 | 1.45 | 1.06 | 0.01 |
| Number of humanities A(S)-Levels taken | 6848 | 0 | 1 | -0.62 | -0.12 | 0.60 | -1.03 | 3.85 | 4.88 | 0.85 | -0.40 | 0.01 |
| STEM pipeline | 3504 | 0 | 1 | -0.48 | -0.01 | 1.39 | -2.13 | 1.98 | 4.11 | 0.14 | -0.58 | 0.02 |

**Table S7.** Counts for categorical educational outcome variables. For binary outcomes, 1 denotes “no” and 2 denotes “yes”. For the STEM pipeline, 1 denotes completing compulsory education, 2 denotes choice of a non-compulsory STEM GCSE, 3 denotes choice of a STEM A(S)-Level and 4 denotes choice of a STEM Bachelor’s degree (or higher).

| variable | n | 1 | 2 | 3 | 4 |
|---|---|---|---|---|---|
| Chose a STEM GCSE | 6514 | 903 | 5611 | | |
| Chose a humanities GCSE | 6514 | 1873 | 4641 | | |
| Chose a STEM A(S)-Level | 6887 | 4291 | 2596 | | |
| Chose a humanities A(S)-Level | 6887 | 3446 | 3441 | | |
| Chose a STEM degree | 4859 | 4201 | 658 | | |
| Chose a humanities degree | 4859 | 4160 | 699 | | |
| STEM pipeline | 3607 | 296 | 1526 | 1223 | 459 |

**Table S8.** Linear regression model estimates and fit indices. spatab = spatial ability; nav = nav; vis = visualization, objmanip = object manipulation, v = verbal ability. .g and .v suffixes denote *g*- and v-corrected predictors, respectively. p.adjusted denotes Benjamini-Hochberg corrected p-values.

| model | nobs | estimate | lower 95% | upper 95% | se | t | p.value | p.adjusted | sig | r.squared | adj.r.squared | sd | F | df | df.residual |
|---|---|---|---|---|---|---|---|---|---|---|---|---|---|---|---|
| GCSE STEM mean grade vs objmanip | 1667 | 0.47 | 0.43 | 0.50 | 0.02 | 23.18 | 3.15E-103 | 5.42E-101 | *** | 0.24 | 0.24 | 0.82 | 537.18 | 1 | 1665 |
| GCSE STEM mean grade vs vis | 1667 | 0.45 | 0.41 | 0.49 | 0.02 | 22.16 | 1.49E-95 | 8.56E-94 | *** | 0.23 | 0.23 | 0.82 | 491.01 | 1 | 1665 |
| GCSE STEM mean grade vs nav | 1667 | 0.44 | 0.40 | 0.48 | 0.02 | 21.05 | 1.98E-87 | 7.85E-86 | *** | 0.21 | 0.21 | 0.83 | 443.22 | 1 | 1665 |
| GCSE STEM mean grade vs spatab | 1667 | 0.45 | 0.41 | 0.49 | 0.02 | 22.16 | 1.49E-95 | 8.56E-94 | *** | 0.23 | 0.23 | 0.82 | 491.01 | 1 | 1665 |
| GCSE STEM mean grade vs objmanip.g | 1318 | 0.15 | 0.10 | 0.20 | 0.03 | 5.71 | 1.36E-08 | 4.06E-08 | *** | 0.02 | 0.02 | 0.93 | 32.65 | 1 | 1316 |
| GCSE STEM mean grade vs vis.g | 1318 | 0.13 | 0.08 | 0.18 | 0.03 | 5.17 | 2.75E-07 | 7.72E-07 | *** | 0.02 | 0.02 | 0.93 | 26.69 | 1 | 1316 |
| GCSE STEM mean grade vs nav.g | 1318 | 0.12 | 0.07 | 0.17 | 0.03 | 4.71 | 2.69E-06 | 7.08E-06 | *** | 0.02 | 0.02 | 0.93 | 22.22 | 1 | 1316 |
| GCSE STEM mean grade vs spatab.g | 1318 | 0.13 | 0.08 | 0.18 | 0.03 | 5.17 | 2.75E-07 | 7.72E-07 | *** | 0.02 | 0.02 | 0.93 | 26.69 | 1 | 1316 |
| GCSE STEM mean grade vs objmanip.v | 1320 | 0.26 | 0.21 | 0.30 | 0.02 | 10.32 | 4.53E-24 | 3.34E-23 | *** | 0.07 | 0.07 | 0.90 | 106.53 | 1 | 1318 |
| GCSE STEM mean grade vs vis.v | 1320 | 0.24 | 0.19 | 0.29 | 0.02 | 9.57 | 5.02E-21 | 3.20E-20 | *** | 0.06 | 0.06 | 0.91 | 91.60 | 1 | 1318 |
| GCSE STEM mean grade vs nav.v | 1320 | 0.22 | 0.17 | 0.27 | 0.02 | 8.74 | 6.82E-18 | 3.38E-17 | *** | 0.05 | 0.05 | 0.91 | 76.43 | 1 | 1318 |
| GCSE STEM mean grade vs spatab.v | 1320 | 0.24 | 0.19 | 0.29 | 0.02 | 9.57 | 5.02E-21 | 3.20E-20 | *** | 0.06 | 0.06 | 0.91 | 91.60 | 1 | 1318 |
| GCSE Maths mean grade vs objmanip | 1660 | 0.47 | 0.43 | 0.51 | 0.02 | 23.38 | 1.00E-104 | 2.58E-102 | *** | 0.25 | 0.25 | 0.81 | 546.66 | 1 | 1658 |
| GCSE Maths mean grade vs vis | 1660 | 0.46 | 0.42 | 0.50 | 0.02 | 22.50 | 4.67E-98 | 3.44E-96 | *** | 0.23 | 0.23 | 0.82 | 506.27 | 1 | 1658 |
| GCSE Maths mean grade vs nav | 1660 | 0.45 | 0.41 | 0.49 | 0.02 | 21.56 | 4.16E-91 | 1.95E-89 | *** | 0.22 | 0.22 | 0.83 | 464.98 | 1 | 1658 |
| GCSE Maths mean grade vs spatab | 1660 | 0.46 | 0.42 | 0.50 | 0.02 | 22.50 | 4.67E-98 | 3.44E-96 | *** | 0.23 | 0.23 | 0.82 | 506.27 | 1 | 1658 |
| GCSE Maths mean grade vs objmanip.g | 1312 | 0.16 | 0.11 | 0.21 | 0.03 | 6.21 | 6.92E-10 | 2.29E-09 | *** | 0.03 | 0.03 | 0.92 | 38.62 | 1 | 1310 |
| GCSE Maths mean grade vs vis.g | 1312 | 0.15 | 0.10 | 0.20 | 0.03 | 5.75 | 1.10E-08 | 3.29E-08 | *** | 0.02 | 0.02 | 0.92 | 33.08 | 1 | 1310 |
| GCSE Maths mean grade vs nav.g | 1312 | 0.14 | 0.09 | 0.19 | 0.03 | 5.31 | 1.29E-07 | 3.67E-07 | *** | 0.02 | 0.02 | 0.92 | 28.18 | 1 | 1310 |
| GCSE Maths mean grade vs spatab.g | 1312 | 0.15 | 0.10 | 0.20 | 0.03 | 5.75 | 1.10E-08 | 3.29E-08 | *** | 0.02 | 0.02 | 0.92 | 33.08 | 1 | 1310 |
| GCSE Maths mean grade vs objmanip.v | 1314 | 0.27 | 0.22 | 0.32 | 0.02 | 11.06 | 3.15E-27 | 2.76E-26 | *** | 0.09 | 0.08 | 0.89 | 122.22 | 1 | 1312 |
| GCSE Maths mean grade vs vis.v | 1314 | 0.26 | 0.21 | 0.31 | 0.02 | 10.39 | 2.47E-24 | 1.87E-23 | *** | 0.08 | 0.08 | 0.90 | 107.85 | 1 | 1312 |
| GCSE Maths mean grade vs nav.v | 1314 | 0.24 | 0.19 | 0.29 | 0.02 | 9.56 | 5.37E-21 | 3.38E-20 | *** | 0.07 | 0.06 | 0.90 | 91.47 | 1 | 1312 |
| GCSE Maths mean grade vs spatab.v | 1314 | 0.26 | 0.21 | 0.31 | 0.02 | 10.39 | 2.47E-24 | 1.87E-23 | *** | 0.08 | 0.08 | 0.90 | 107.85 | 1 | 1312 |
| GCSE Science mean grade vs objmanip | 1598 | 0.44 | 0.39 | 0.48 | 0.02 | 20.26 | 2.13E-81 | 6.45E-80 | *** | 0.20 | 0.20 | 0.84 | 410.44 | 1 | 1596 |
| GCSE Science mean grade vs vis | 1598 | 0.42 | 0.38 | 0.47 | 0.02 | 19.29 | 1.14E-74 | 2.55E-73 | *** | 0.19 | 0.19 | 0.85 | 371.97 | 1 | 1596 |
| GCSE Science mean grade vs nav | 1598 | 0.41 | 0.37 | 0.45 | 0.02 | 18.33 | 2.73E-68 | 5.23E-67 | *** | 0.17 | 0.17 | 0.86 | 336.16 | 1 | 1596 |
| GCSE Science mean grade vs spatab | 1598 | 0.42 | 0.38 | 0.47 | 0.02 | 19.29 | 1.14E-74 | 2.55E-73 | *** | 0.19 | 0.19 | 0.85 | 371.97 | 1 | 1596 |
| GCSE Science mean grade vs objmanip.g | 1274 | 0.13 | 0.07 | 0.18 | 0.03 | 4.74 | 2.38E-06 | 6.29E-06 | *** | 0.02 | 0.02 | 0.94 | 22.47 | 1 | 1272 |
| GCSE Science mean grade vs vis.g | 1274 | 0.11 | 0.06 | 0.16 | 0.03 | 4.19 | 2.98E-05 | 7.21E-05 | *** | 0.01 | 0.01 | 0.94 | 17.56 | 1 | 1272 |
| GCSE Science mean grade vs nav.g | 1274 | 0.10 | 0.05 | 0.16 | 0.03 | 3.89 | 1.06E-04 | 2.44E-04 | *** | 0.01 | 0.01 | 0.95 | 15.13 | 1 | 1272 |
| GCSE Science mean grade vs spatab.g | 1274 | 0.11 | 0.06 | 0.16 | 0.03 | 4.19 | 2.98E-05 | 7.21E-05 | *** | 0.01 | 0.01 | 0.94 | 17.56 | 1 | 1272 |
| GCSE Science mean grade vs objmanip.v | 1276 | 0.22 | 0.17 | 0.28 | 0.03 | 8.61 | 2.05E-17 | 1.00E-16 | *** | 0.06 | 0.05 | 0.92 | 74.20 | 1 | 1274 |
| GCSE Science mean grade vs vis.v | 1276 | 0.21 | 0.15 | 0.26 | 0.03 | 7.90 | 6.09E-15 | 2.60E-14 | *** | 0.05 | 0.05 | 0.93 | 62.38 | 1 | 1274 |
| GCSE Science mean grade vs nav.v | 1276 | 0.19 | 0.14 | 0.24 | 0.03 | 7.27 | 6.46E-13 | 2.53E-12 | *** | 0.04 | 0.04 | 0.93 | 52.79 | 1 | 1274 |
| GCSE Science mean grade vs spatab.v | 1276 | 0.21 | 0.15 | 0.26 | 0.03 | 7.90 | 6.09E-15 | 2.60E-14 | *** | 0.05 | 0.05 | 0.93 | 62.38 | 1 | 1274 |
| GCSE Physics grade vs objmanip | 708 | 0.39 | 0.31 | 0.46 | 0.04 | 10.69 | 8.22E-25 | 6.52E-24 | *** | 0.14 | 0.14 | 0.88 | 114.21 | 1 | 706 |

| | | | | | | | | | | | | | | | |
|---|---|---|---|---|---|---|---|---|---|---|---|---|---|---|---|
| GCSE Physics grade vs vis | 708 | 0.36 | 0.29 | 0.44 | 0.04 | 9.69 | 6.27E-21 | 3.76E-20 | *** | 0.12 | 0.12 | 0.89 | 93.89 | 1 | 706 |
| GCSE Physics grade vs nav | 708 | 0.34 | 0.26 | 0.41 | 0.04 | 8.88 | 5.54E-18 | 2.78E-17 | *** | 0.10 | 0.10 | 0.90 | 78.83 | 1 | 706 |
| GCSE Physics grade vs spatab | 708 | 0.36 | 0.29 | 0.44 | 0.04 | 9.69 | 6.27E-21 | 3.76E-20 | *** | 0.12 | 0.12 | 0.89 | 93.89 | 1 | 706 |
| GCSE Physics grade vs objmanip.g | 573 | 0.12 | 0.04 | 0.19 | 0.04 | 2.91 | 3.77E-03 | 7.24E-03 | ** | 0.01 | 0.01 | 0.92 | 8.46 | 1 | 571 |
| GCSE Physics grade vs vis.g | 573 | 0.08 | 0.00 | 0.15 | 0.04 | 1.90 | 5.81E-02 | 8.63E-02 | | 0.01 | 0.00 | 0.92 | 3.61 | 1 | 571 |
| GCSE Physics grade vs nav.g | 573 | 0.06 | -0.02 | 0.13 | 0.04 | 1.38 | 1.67E-01 | 2.18E-01 | | 0.00 | 0.00 | 0.93 | 1.91 | 1 | 571 |
| GCSE Physics grade vs spatab.g | 573 | 0.08 | 0.00 | 0.15 | 0.04 | 1.90 | 5.81E-02 | 8.63E-02 | | 0.01 | 0.00 | 0.92 | 3.61 | 1 | 571 |
| GCSE Physics grade vs objmanip.v | 575 | 0.20 | 0.12 | 0.28 | 0.04 | 5.16 | 3.41E-07 | 9.46E-07 | *** | 0.04 | 0.04 | 0.90 | 26.63 | 1 | 573 |
| GCSE Physics grade vs vis.v | 575 | 0.16 | 0.09 | 0.24 | 0.04 | 4.14 | 4.08E-05 | 9.65E-05 | *** | 0.03 | 0.03 | 0.91 | 17.10 | 1 | 573 |
| GCSE Physics grade vs nav.v | 575 | 0.14 | 0.06 | 0.21 | 0.04 | 3.45 | 5.92E-04 | 1.26E-03 | ** | 0.02 | 0.02 | 0.92 | 11.93 | 1 | 573 |
| GCSE Physics grade vs spatab.v | 575 | 0.16 | 0.09 | 0.24 | 0.04 | 4.14 | 4.08E-05 | 9.65E-05 | *** | 0.03 | 0.03 | 0.91 | 17.10 | 1 | 573 |
| GCSE Chemistry grade vs objmanip | 713 | 0.35 | 0.28 | 0.43 | 0.04 | 9.65 | 8.48E-21 | 5.03E-20 | *** | 0.12 | 0.11 | 0.90 | 93.17 | 1 | 711 |
| GCSE Chemistry grade vs vis | 713 | 0.33 | 0.25 | 0.40 | 0.04 | 8.59 | 5.68E-17 | 2.64E-16 | *** | 0.09 | 0.09 | 0.91 | 73.71 | 1 | 711 |
| GCSE Chemistry grade vs nav | 713 | 0.30 | 0.23 | 0.38 | 0.04 | 7.80 | 2.25E-14 | 9.42E-14 | *** | 0.08 | 0.08 | 0.92 | 60.81 | 1 | 711 |
| GCSE Chemistry grade vs spatab | 713 | 0.33 | 0.25 | 0.40 | 0.04 | 8.59 | 5.68E-17 | 2.64E-16 | *** | 0.09 | 0.09 | 0.91 | 73.71 | 1 | 711 |
| GCSE Chemistry grade vs objmanip.g | 577 | 0.11 | 0.03 | 0.19 | 0.04 | 2.65 | 8.33E-03 | 1.52E-02 | * | 0.01 | 0.01 | 0.96 | 7.01 | 1 | 575 |
| GCSE Chemistry grade vs vis.g | 577 | 0.07 | -0.01 | 0.15 | 0.04 | 1.61 | 1.08E-01 | 1.46E-01 | | 0.00 | 0.00 | 0.96 | 2.59 | 1 | 575 |
| GCSE Chemistry grade vs nav.g | 577 | 0.04 | -0.04 | 0.12 | 0.04 | 0.97 | 3.34E-01 | 4.00E-01 | | 0.00 | 0.00 | 0.96 | 0.93 | 1 | 575 |
| GCSE Chemistry grade vs spatab.g | 577 | 0.07 | -0.01 | 0.15 | 0.04 | 1.61 | 1.08E-01 | 1.46E-01 | | 0.00 | 0.00 | 0.96 | 2.59 | 1 | 575 |
| GCSE Chemistry grade vs objmanip.v | 579 | 0.19 | 0.11 | 0.27 | 0.04 | 4.66 | 3.92E-06 | 1.03E-05 | *** | 0.04 | 0.03 | 0.95 | 21.72 | 1 | 577 |
| GCSE Chemistry grade vs vis.v | 579 | 0.15 | 0.07 | 0.23 | 0.04 | 3.64 | 2.94E-04 | 6.55E-04 | *** | 0.02 | 0.02 | 0.95 | 13.27 | 1 | 577 |
| GCSE Chemistry grade vs nav.v | 579 | 0.12 | 0.04 | 0.20 | 0.04 | 2.86 | 4.39E-03 | 8.35E-03 | ** | 0.01 | 0.01 | 0.96 | 8.18 | 1 | 577 |
| GCSE Chemistry grade vs spatab.v | 579 | 0.15 | 0.07 | 0.23 | 0.04 | 3.64 | 2.94E-04 | 6.55E-04 | *** | 0.02 | 0.02 | 0.95 | 13.27 | 1 | 577 |
| GCSE Biology grade vs objmanip | 712 | 0.38 | 0.31 | 0.45 | 0.04 | 10.52 | 3.63E-24 | 2.72E-23 | *** | 0.13 | 0.13 | 0.90 | 110.75 | 1 | 710 |
| GCSE Biology grade vs vis | 712 | 0.36 | 0.29 | 0.44 | 0.04 | 9.70 | 5.78E-21 | 3.55E-20 | *** | 0.12 | 0.12 | 0.90 | 94.04 | 1 | 710 |
| GCSE Biology grade vs nav | 712 | 0.34 | 0.27 | 0.42 | 0.04 | 9.06 | 1.24E-18 | 6.52E-18 | *** | 0.10 | 0.10 | 0.91 | 82.10 | 1 | 710 |
| GCSE Biology grade vs spatab | 712 | 0.36 | 0.29 | 0.44 | 0.04 | 9.70 | 5.78E-21 | 3.55E-20 | *** | 0.12 | 0.12 | 0.90 | 94.04 | 1 | 710 |
| GCSE Biology grade vs objmanip.g | 577 | 0.12 | 0.04 | 0.20 | 0.04 | 2.90 | 3.84E-03 | 7.34E-03 | ** | 0.01 | 0.01 | 0.96 | 8.43 | 1 | 575 |
| GCSE Biology grade vs vis.g | 577 | 0.09 | 0.01 | 0.17 | 0.04 | 2.19 | 2.90E-02 | 4.63E-02 | * | 0.01 | 0.01 | 0.97 | 4.79 | 1 | 575 |
| GCSE Biology grade vs nav.g | 577 | 0.08 | -0.01 | 0.16 | 0.04 | 1.82 | 7.00E-02 | 1.01E-01 | | 0.01 | 0.00 | 0.97 | 3.30 | 1 | 575 |
| GCSE Biology grade vs spatab.g | 577 | 0.09 | 0.01 | 0.17 | 0.04 | 2.19 | 2.90E-02 | 4.63E-02 | * | 0.01 | 0.01 | 0.97 | 4.79 | 1 | 575 |
| GCSE Biology grade vs objmanip.v | 579 | 0.21 | 0.13 | 0.29 | 0.04 | 5.17 | 3.24E-07 | 9.02E-07 | *** | 0.04 | 0.04 | 0.95 | 26.73 | 1 | 577 |
| GCSE Biology grade vs vis.v | 579 | 0.18 | 0.10 | 0.26 | 0.04 | 4.44 | 1.10E-05 | 2.76E-05 | *** | 0.03 | 0.03 | 0.95 | 19.67 | 1 | 577 |
| GCSE Biology grade vs nav.v | 579 | 0.16 | 0.08 | 0.24 | 0.04 | 3.89 | 1.13E-04 | 2.60E-04 | *** | 0.03 | 0.02 | 0.96 | 15.11 | 1 | 577 |
| GCSE Biology grade vs spatab.v | 579 | 0.18 | 0.10 | 0.26 | 0.04 | 4.44 | 1.10E-05 | 2.76E-05 | *** | 0.03 | 0.03 | 0.95 | 19.67 | 1 | 577 |
| A(S)-Level STEM mean grade vs objmanip | 820 | 0.29 | 0.22 | 0.37 | 0.04 | 7.81 | 1.72E-14 | 7.27E-14 | *** | 0.07 | 0.07 | 0.97 | 61.04 | 1 | 818 |
| A(S)-Level STEM mean grade vs vis | 820 | 0.28 | 0.21 | 0.36 | 0.04 | 7.37 | 4.25E-13 | 1.67E-12 | *** | 0.06 | 0.06 | 0.98 | 54.28 | 1 | 818 |
| A(S)-Level STEM mean grade vs nav | 820 | 0.27 | 0.19 | 0.35 | 0.04 | 6.94 | 7.81E-12 | 2.96E-11 | *** | 0.06 | 0.05 | 0.98 | 48.21 | 1 | 818 |
| A(S)-Level STEM mean grade vs spatab | 820 | 0.28 | 0.21 | 0.36 | 0.04 | 7.37 | 4.25E-13 | 1.67E-12 | *** | 0.06 | 0.06 | 0.98 | 54.28 | 1 | 818 |
| A(S)-Level STEM mean grade vs objmanip.g | 656 | 0.09 | 0.01 | 0.17 | 0.04 | 2.17 | 3.06E-02 | 4.86E-02 | * | 0.01 | 0.01 | 1.01 | 4.69 | 1 | 654 |
| A(S)-Level STEM mean grade vs vis.g | 656 | 0.07 | -0.01 | 0.16 | 0.04 | 1.80 | 7.30E-02 | 1.04E-01 | | 0.00 | 0.00 | 1.01 | 3.22 | 1 | 654 |
| A(S)-Level STEM mean grade vs nav.g | 656 | 0.06 | -0.02 | 0.14 | 0.04 | 1.48 | 1.39E-01 | 1.83E-01 | | 0.00 | 0.00 | 1.01 | 2.20 | 1 | 654 |
| A(S)-Level STEM mean grade vs spatab.g | 656 | 0.07 | -0.01 | 0.16 | 0.04 | 1.80 | 7.30E-02 | 1.04E-01 | | 0.00 | 0.00 | 1.01 | 3.22 | 1 | 654 |

| | | | | | | | | | | | | | | | |
|---|---|---|---|---|---|---|---|---|---|---|---|---|---|---|---|
| A(S)-Level STEM mean grade vs objmanip.v | 656 | 0.14 | 0.06 | 0.22 | 0.04 | 3.44 | 6.16E-04 | 1.30E-03 | ** | 0.02 | 0.02 | 1.00 | 11.84 | 1 | 654 |
| A(S)-Level STEM mean grade vs vis.v | 656 | 0.13 | 0.04 | 0.21 | 0.04 | 3.04 | 2.43E-03 | 4.77E-03 | ** | 0.01 | 0.01 | 1.00 | 9.26 | 1 | 654 |
| A(S)-Level STEM mean grade vs nav.v | 656 | 0.11 | 0.03 | 0.19 | 0.04 | 2.64 | 8.46E-03 | 1.54E-02 | * | 0.01 | 0.01 | 1.00 | 6.98 | 1 | 654 |
| A(S)-Level STEM mean grade vs spatab.v | 656 | 0.13 | 0.04 | 0.21 | 0.04 | 3.04 | 2.43E-03 | 4.77E-03 | ** | 0.01 | 0.01 | 1.00 | 9.26 | 1 | 654 |
| A(S)-Level Maths mean grade vs objmanip | 484 | 0.24 | 0.14 | 0.34 | 0.05 | 4.84 | 1.77E-06 | 4.72E-06 | *** | 0.05 | 0.04 | 0.98 | 23.41 | 1 | 482 |
| A(S)-Level Maths mean grade vs vis | 484 | 0.23 | 0.13 | 0.33 | 0.05 | 4.55 | 6.75E-06 | 1.73E-05 | *** | 0.04 | 0.04 | 0.98 | 20.72 | 1 | 482 |
| A(S)-Level Maths mean grade vs nav | 484 | 0.22 | 0.12 | 0.32 | 0.05 | 4.25 | 2.52E-05 | 6.17E-05 | *** | 0.04 | 0.03 | 0.98 | 18.10 | 1 | 482 |
| A(S)-Level Maths mean grade vs spatab | 484 | 0.23 | 0.13 | 0.33 | 0.05 | 4.55 | 6.75E-06 | 1.73E-05 | *** | 0.04 | 0.04 | 0.98 | 20.72 | 1 | 482 |
| A(S)-Level Maths mean grade vs objmanip.g | 396 | 0.05 | -0.05 | 0.15 | 0.05 | 0.93 | 3.54E-01 | 4.20E-01 | | 0.00 | 0.00 | 0.98 | 0.86 | 1 | 394 |
| A(S)-Level Maths mean grade vs vis.g | 396 | 0.04 | -0.06 | 0.14 | 0.05 | 0.71 | 4.77E-01 | 5.41E-01 | | 0.00 | 0.00 | 0.98 | 0.51 | 1 | 394 |
| A(S)-Level Maths mean grade vs nav.g | 396 | 0.03 | -0.07 | 0.13 | 0.05 | 0.59 | 5.58E-01 | 6.23E-01 | | 0.00 | 0.00 | 0.98 | 0.34 | 1 | 394 |
| A(S)-Level Maths mean grade vs spatab.g | 396 | 0.04 | -0.06 | 0.14 | 0.05 | 0.71 | 4.77E-01 | 5.41E-01 | | 0.00 | 0.00 | 0.98 | 0.51 | 1 | 394 |
| A(S)-Level Maths mean grade vs objmanip.v | 396 | 0.10 | -0.01 | 0.20 | 0.05 | 1.85 | 6.56E-02 | 9.59E-02 | | 0.01 | 0.01 | 0.98 | 3.41 | 1 | 394 |
| A(S)-Level Maths mean grade vs vis.v | 396 | 0.08 | -0.02 | 0.19 | 0.05 | 1.60 | 1.10E-01 | 1.48E-01 | | 0.01 | 0.00 | 0.98 | 2.56 | 1 | 394 |
| A(S)-Level Maths mean grade vs nav.v | 396 | 0.07 | -0.03 | 0.17 | 0.05 | 1.39 | 1.66E-01 | 2.17E-01 | | 0.00 | 0.00 | 0.98 | 1.93 | 1 | 394 |
| A(S)-Level Maths mean grade vs spatab.v | 396 | 0.08 | -0.02 | 0.19 | 0.05 | 1.60 | 1.10E-01 | 1.48E-01 | | 0.01 | 0.00 | 0.98 | 2.56 | 1 | 394 |
| A(S)-Level Science mean grade vs objmanip | 592 | 0.29 | 0.20 | 0.38 | 0.05 | 6.41 | 2.91E-10 | 1.01E-09 | *** | 0.07 | 0.06 | 0.97 | 41.14 | 1 | 590 |
| A(S)-Level Science mean grade vs vis | 592 | 0.26 | 0.17 | 0.35 | 0.05 | 5.69 | 2.02E-08 | 5.93E-08 | *** | 0.05 | 0.05 | 0.98 | 32.35 | 1 | 590 |
| A(S)-Level Science mean grade vs nav | 592 | 0.25 | 0.16 | 0.34 | 0.05 | 5.36 | 1.21E-07 | 3.44E-07 | *** | 0.05 | 0.04 | 0.98 | 28.71 | 1 | 590 |
| A(S)-Level Science mean grade vs spatab | 592 | 0.26 | 0.17 | 0.35 | 0.05 | 5.69 | 2.02E-08 | 5.93E-08 | *** | 0.05 | 0.05 | 0.98 | 32.35 | 1 | 590 |
| A(S)-Level Science mean grade vs objmanip.g | 468 | 0.04 | -0.06 | 0.15 | 0.05 | 0.86 | 3.92E-01 | 4.54E-01 | | 0.00 | 0.00 | 1.01 | 0.73 | 1 | 466 |
| A(S)-Level Science mean grade vs vis.g | 468 | 0.01 | -0.08 | 0.11 | 0.05 | 0.29 | 7.69E-01 | 8.14E-01 | | 0.00 | 0.00 | 1.01 | 0.09 | 1 | 466 |
| A(S)-Level Science mean grade vs nav.g | 468 | 0.00 | -0.10 | 0.10 | 0.05 | 0.03 | 9.78E-01 | 9.81E-01 | | 0.00 | 0.00 | 1.01 | 0.00 | 1 | 466 |
| A(S)-Level Science mean grade vs spatab.g | 468 | 0.01 | -0.08 | 0.11 | 0.05 | 0.29 | 7.69E-01 | 8.14E-01 | | 0.00 | 0.00 | 1.01 | 0.09 | 1 | 466 |
| A(S)-Level Science mean grade vs objmanip.v | 468 | 0.12 | 0.02 | 0.23 | 0.05 | 2.40 | 1.69E-02 | 2.86E-02 | * | 0.01 | 0.01 | 1.01 | 5.75 | 1 | 466 |
| A(S)-Level Science mean grade vs vis.v | 468 | 0.09 | -0.01 | 0.19 | 0.05 | 1.79 | 7.48E-02 | 1.06E-01 | | 0.01 | 0.00 | 1.01 | 3.19 | 1 | 466 |
| A(S)-Level Science mean grade vs nav.v | 468 | 0.07 | -0.03 | 0.17 | 0.05 | 1.39 | 1.65E-01 | 2.17E-01 | | 0.00 | 0.00 | 1.01 | 1.93 | 1 | 466 |
| A(S)-Level Science mean grade vs spatab.v | 468 | 0.09 | -0.01 | 0.19 | 0.05 | 1.79 | 7.48E-02 | 1.06E-01 | | 0.01 | 0.00 | 1.01 | 3.19 | 1 | 466 |
| STEM pipeline vs objmanip | 1254 | 0.42 | 0.37 | 0.48 | 0.03 | 15.65 | 1.57E-50 | 2.46E-49 | *** | 0.16 | 0.16 | 0.95 | 244.77 | 1 | 1252 |
| STEM pipeline vs vis | 1254 | 0.40 | 0.35 | 0.45 | 0.03 | 14.36 | 2.05E-43 | 2.78E-42 | *** | 0.14 | 0.14 | 0.97 | 206.28 | 1 | 1252 |
| STEM pipeline vs nav | 1254 | 0.38 | 0.32 | 0.43 | 0.03 | 13.34 | 4.53E-38 | 4.98E-37 | *** | 0.12 | 0.12 | 0.98 | 178.03 | 1 | 1252 |
| STEM pipeline vs spatab | 1254 | 0.40 | 0.35 | 0.45 | 0.03 | 14.36 | 2.05E-43 | 2.78E-42 | *** | 0.14 | 0.14 | 0.97 | 206.28 | 1 | 1252 |
| STEM pipeline vs objmanip.g | 1042 | 0.26 | 0.19 | 0.32 | 0.03 | 7.95 | 4.81E-15 | 2.09E-14 | *** | 0.06 | 0.06 | 1.01 | 63.21 | 1 | 1040 |
| STEM pipeline vs vis.g | 1042 | 0.22 | 0.15 | 0.28 | 0.03 | 6.73 | 2.85E-11 | 1.04E-10 | *** | 0.04 | 0.04 | 1.02 | 45.26 | 1 | 1040 |
| STEM pipeline vs nav.g | 1042 | 0.19 | 0.13 | 0.25 | 0.03 | 5.86 | 6.08E-09 | 1.87E-08 | *** | 0.03 | 0.03 | 1.03 | 34.38 | 1 | 1040 |
| STEM pipeline vs spatab.g | 1042 | 0.22 | 0.15 | 0.28 | 0.03 | 6.73 | 2.85E-11 | 1.04E-10 | *** | 0.04 | 0.04 | 1.02 | 45.26 | 1 | 1040 |
| STEM pipeline vs objmanip.v | 1043 | 0.33 | 0.27 | 0.39 | 0.03 | 10.54 | 9.43E-25 | 7.38E-24 | *** | 0.10 | 0.10 | 0.99 | 111.10 | 1 | 1041 |
| STEM pipeline vs vis.v | 1043 | 0.29 | 0.23 | 0.35 | 0.03 | 9.32 | 6.53E-20 | 3.70E-19 | *** | 0.08 | 0.08 | 1.00 | 86.93 | 1 | 1041 |
| STEM pipeline vs nav.v | 1043 | 0.26 | 0.20 | 0.32 | 0.03 | 8.33 | 2.47E-16 | 1.11E-15 | *** | 0.06 | 0.06 | 1.01 | 69.44 | 1 | 1041 |
| STEM pipeline vs spatab.v | 1043 | 0.29 | 0.23 | 0.35 | 0.03 | 9.32 | 6.53E-20 | 3.70E-19 | *** | 0.08 | 0.08 | 1.00 | 86.93 | 1 | 1041 |
| Number of STEM GCSEs takenvs objmanip | 1671 | 0.25 | 0.20 | 0.29 | 0.02 | 10.94 | 5.74E-27 | 4.94E-26 | *** | 0.07 | 0.07 | 0.91 | 119.76 | 1 | 1669 |
| Number of STEM GCSEs takenvs vis | 1671 | 0.24 | 0.20 | 0.28 | 0.02 | 10.63 | 1.44E-25 | 1.16E-24 | *** | 0.06 | 0.06 | 0.91 | 112.92 | 1 | 1669 |
| Number of STEM GCSEs takenvs nav | 1671 | 0.23 | 0.19 | 0.28 | 0.02 | 10.24 | 6.48E-24 | 4.71E-23 | *** | 0.06 | 0.06 | 0.92 | 104.89 | 1 | 1669 |

| | | | | | | | | | | | | | | | |
|---|---|---|---|---|---|---|---|---|---|---|---|---|---|---|---|
| Number of STEM GCSEs takenvs spatab | 1671 | 0.24 | 0.20 | 0.28 | 0.02 | 10.63 | 1.44E-25 | 1.16E-24 | *** | 0.06 | 0.06 | 0.91 | 112.92 | 1 | 1669 |
| Number of STEM GCSEs takenvs objmanip.g | 1322 | 0.11 | 0.06 | 0.16 | 0.03 | 4.37 | 1.34E-05 | 3.34E-05 | *** | 0.01 | 0.01 | 0.93 | 19.10 | 1 | 1320 |
| Number of STEM GCSEs takenvs vis.g | 1322 | 0.11 | 0.06 | 0.16 | 0.03 | 4.23 | 2.47E-05 | 6.07E-05 | *** | 0.01 | 0.01 | 0.93 | 17.91 | 1 | 1320 |
| Number of STEM GCSEs takenvs nav.g | 1322 | 0.10 | 0.05 | 0.15 | 0.03 | 3.95 | 8.24E-05 | 1.92E-04 | *** | 0.01 | 0.01 | 0.93 | 15.60 | 1 | 1320 |
| Number of STEM GCSEs takenvs spatab.g | 1322 | 0.11 | 0.06 | 0.16 | 0.03 | 4.23 | 2.47E-05 | 6.07E-05 | *** | 0.01 | 0.01 | 0.93 | 17.91 | 1 | 1320 |
| Number of STEM GCSEs takenvs objmanip.v | 1324 | 0.16 | 0.11 | 0.21 | 0.03 | 6.46 | 1.43E-10 | 4.97E-10 | *** | 0.03 | 0.03 | 0.92 | 41.79 | 1 | 1322 |
| Number of STEM GCSEs takenvs vis.v | 1324 | 0.16 | 0.11 | 0.21 | 0.03 | 6.25 | 5.56E-10 | 1.85E-09 | *** | 0.03 | 0.03 | 0.92 | 39.05 | 1 | 1322 |
| Number of STEM GCSEs takenvs nav.v | 1324 | 0.15 | 0.10 | 0.20 | 0.03 | 5.83 | 6.92E-09 | 2.10E-08 | *** | 0.03 | 0.02 | 0.93 | 34.00 | 1 | 1322 |
| Number of STEM GCSEs takenvs spatab.v | 1324 | 0.16 | 0.11 | 0.21 | 0.03 | 6.25 | 5.56E-10 | 1.85E-09 | *** | 0.03 | 0.03 | 0.92 | 39.05 | 1 | 1322 |
| Number of STEM A(S)-Levels takenvs objmanip | 1703 | 0.43 | 0.38 | 0.47 | 0.02 | 17.78 | 5.02E-65 | 9.25E-64 | *** | 0.16 | 0.16 | 1.00 | 316.27 | 1 | 1701 |
| Number of STEM A(S)-Levels takenvs vis | 1703 | 0.40 | 0.35 | 0.45 | 0.02 | 16.42 | 2.29E-56 | 3.82E-55 | *** | 0.14 | 0.14 | 1.01 | 269.68 | 1 | 1701 |
| Number of STEM A(S)-Levels takenvs nav | 1703 | 0.38 | 0.33 | 0.43 | 0.02 | 15.40 | 3.37E-50 | 5.12E-49 | *** | 0.12 | 0.12 | 1.02 | 237.17 | 1 | 1701 |
| Number of STEM A(S)-Levels takenvs spatab | 1703 | 0.40 | 0.35 | 0.45 | 0.02 | 16.42 | 2.29E-56 | 3.82E-55 | *** | 0.14 | 0.14 | 1.01 | 269.68 | 1 | 1701 |
| Number of STEM A(S)-Levels takenvs objmanip.g | 1290 | 0.26 | 0.20 | 0.32 | 0.03 | 8.79 | 4.58E-18 | 2.34E-17 | *** | 0.06 | 0.06 | 1.07 | 77.32 | 1 | 1288 |
| Number of STEM A(S)-Levels takenvs vis.g | 1290 | 0.23 | 0.17 | 0.29 | 0.03 | 7.53 | 9.37E-14 | 3.87E-13 | *** | 0.04 | 0.04 | 1.08 | 56.73 | 1 | 1288 |
| Number of STEM A(S)-Levels takenvs nav.g | 1290 | 0.20 | 0.14 | 0.26 | 0.03 | 6.64 | 4.60E-11 | 1.64E-10 | *** | 0.03 | 0.03 | 1.09 | 44.10 | 1 | 1288 |
| Number of STEM A(S)-Levels takenvs spatab.g | 1290 | 0.23 | 0.17 | 0.29 | 0.03 | 7.53 | 9.37E-14 | 3.87E-13 | *** | 0.04 | 0.04 | 1.08 | 56.73 | 1 | 1288 |
| Number of STEM A(S)-Levels takenvs objmanip.v | 1291 | 0.33 | 0.28 | 0.39 | 0.03 | 11.46 | 5.09E-29 | 4.53E-28 | *** | 0.09 | 0.09 | 1.05 | 131.32 | 1 | 1289 |
| Number of STEM A(S)-Levels takenvs vis.v | 1291 | 0.30 | 0.24 | 0.36 | 0.03 | 10.18 | 1.91E-23 | 1.35E-22 | *** | 0.07 | 0.07 | 1.06 | 103.53 | 1 | 1289 |
| Number of STEM A(S)-Levels takenvs nav.v | 1291 | 0.27 | 0.21 | 0.33 | 0.03 | 9.15 | 2.09E-19 | 1.14E-18 | *** | 0.06 | 0.06 | 1.07 | 83.79 | 1 | 1289 |
| Number of STEM A(S)-Levels takenvs spatab.v | 1291 | 0.30 | 0.24 | 0.36 | 0.03 | 10.18 | 1.91E-23 | 1.35E-22 | *** | 0.07 | 0.07 | 1.06 | 103.53 | 1 | 1289 |
| GCSE humanities mean gradevs objmanip | 1666 | 0.33 | 0.29 | 0.37 | 0.02 | 15.54 | 5.70E-51 | 9.19E-50 | *** | 0.13 | 0.13 | 0.86 | 241.57 | 1 | 1664 |
| GCSE humanities mean gradevs vis | 1666 | 0.32 | 0.28 | 0.36 | 0.02 | 15.00 | 8.38E-48 | 1.20E-46 | *** | 0.12 | 0.12 | 0.86 | 225.01 | 1 | 1664 |
| GCSE humanities mean gradevs nav | 1666 | 0.31 | 0.27 | 0.35 | 0.02 | 14.15 | 5.16E-43 | 6.83E-42 | *** | 0.11 | 0.11 | 0.87 | 200.26 | 1 | 1664 |
| GCSE humanities mean gradevs spatab | 1666 | 0.32 | 0.28 | 0.36 | 0.02 | 15.00 | 8.38E-48 | 1.20E-46 | *** | 0.12 | 0.12 | 0.86 | 225.01 | 1 | 1664 |
| GCSE humanities mean gradevs objmanip.g | 1317 | 0.01 | -0.04 | 0.06 | 0.03 | 0.26 | 7.94E-01 | 8.29E-01 | | 0.00 | 0.00 | 0.92 | 0.07 | 1 | 1315 |
| GCSE humanities mean gradevs vis.g | 1317 | 0.00 | -0.05 | 0.05 | 0.03 | 0.06 | 9.48E-01 | 9.54E-01 | | 0.00 | 0.00 | 0.92 | 0.00 | 1 | 1315 |
| GCSE humanities mean gradevs nav.g | 1317 | 0.00 | -0.05 | 0.05 | 0.03 | -0.16 | 8.73E-01 | 8.97E-01 | | 0.00 | 0.00 | 0.92 | 0.03 | 1 | 1315 |
| GCSE humanities mean gradevs spatab.g | 1317 | 0.00 | -0.05 | 0.05 | 0.03 | 0.06 | 9.48E-01 | 9.54E-01 | | 0.00 | 0.00 | 0.92 | 0.00 | 1 | 1315 |
| GCSE humanities mean gradevs objmanip.v | 1319 | 0.10 | 0.05 | 0.15 | 0.03 | 3.93 | 8.77E-05 | 2.03E-04 | *** | 0.01 | 0.01 | 0.91 | 15.48 | 1 | 1317 |
| GCSE humanities mean gradevs vis.v | 1319 | 0.09 | 0.04 | 0.14 | 0.03 | 3.54 | 4.16E-04 | 9.05E-04 | *** | 0.01 | 0.01 | 0.91 | 12.52 | 1 | 1317 |
| GCSE humanities mean gradevs nav.v | 1319 | 0.07 | 0.02 | 0.12 | 0.03 | 2.95 | 3.20E-03 | 6.22E-03 | ** | 0.01 | 0.01 | 0.92 | 8.72 | 1 | 1317 |
| GCSE humanities mean gradevs spatab.v | 1319 | 0.09 | 0.04 | 0.14 | 0.03 | 3.54 | 4.16E-04 | 9.05E-04 | *** | 0.01 | 0.01 | 0.91 | 12.52 | 1 | 1317 |
| A(S)-Level humanities mean gradevs objmanip | 1054 | 0.10 | 0.03 | 0.16 | 0.03 | 3.02 | 2.62E-03 | 5.12E-03 | ** | 0.01 | 0.01 | 1.00 | 9.10 | 1 | 1052 |
| A(S)-Level humanities mean gradevs vis | 1054 | 0.09 | 0.02 | 0.15 | 0.03 | 2.62 | 8.91E-03 | 1.61E-02 | * | 0.01 | 0.01 | 1.00 | 6.87 | 1 | 1052 |
| A(S)-Level humanities mean gradevs nav | 1054 | 0.08 | 0.02 | 0.15 | 0.03 | 2.44 | 1.50E-02 | 2.58E-02 | * | 0.01 | 0.00 | 1.00 | 5.93 | 1 | 1052 |
| A(S)-Level humanities mean gradevs spatab | 1054 | 0.09 | 0.02 | 0.15 | 0.03 | 2.62 | 8.91E-03 | 1.61E-02 | * | 0.01 | 0.01 | 1.00 | 6.87 | 1 | 1052 |
| A(S)-Level humanities mean gradevs objmanip.g | 823 | -0.11 | -0.18 | -0.04 | 0.04 | -3.20 | 1.45E-03 | 2.93E-03 | ** | 0.01 | 0.01 | 0.99 | 10.21 | 1 | 821 |
| A(S)-Level humanities mean gradevs vis.g | 823 | -0.12 | -0.19 | -0.06 | 0.04 | -3.54 | 4.22E-04 | 9.10E-04 | *** | 0.02 | 0.01 | 0.99 | 12.54 | 1 | 821 |
| A(S)-Level humanities mean gradevs nav.g | 823 | -0.12 | -0.19 | -0.05 | 0.03 | -3.44 | 6.03E-04 | 1.28E-03 | ** | 0.01 | 0.01 | 0.99 | 11.86 | 1 | 821 |
| A(S)-Level humanities mean gradevs spatab.g | 823 | -0.12 | -0.19 | -0.06 | 0.04 | -3.54 | 4.22E-04 | 9.10E-04 | *** | 0.02 | 0.01 | 0.99 | 12.54 | 1 | 821 |
| A(S)-Level humanities mean gradevs objmanip.v | 824 | -0.07 | -0.14 | 0.00 | 0.04 | -1.96 | 4.99E-02 | 7.53E-02 | | 0.00 | 0.00 | 0.99 | 3.85 | 1 | 822 |
| A(S)-Level humanities mean gradevs vis.v | 824 | -0.08 | -0.15 | -0.01 | 0.04 | -2.37 | 1.81E-02 | 3.02E-02 | * | 0.01 | 0.01 | 0.99 | 5.61 | 1 | 822 |

| | | | | | | | | | | | | | | | |
|---|---|---|---|---|---|---|---|---|---|---|---|---|---|---|---|
| A(S)-Level humanities mean gradevs nav.v | 824 | -0.09 | -0.16 | -0.02 | 0.04 | -2.47 | 1.38E-02 | 2.41E-02 | * | 0.01 | 0.01 | 0.99 | 6.10 | 1 | 822 |
| A(S)-Level humanities mean gradevs spatab.v | 824 | -0.08 | -0.15 | -0.01 | 0.04 | -2.37 | 1.81E-02 | 3.02E-02 | * | 0.01 | 0.01 | 0.99 | 5.61 | 1 | 822 |
| Number of humanities GCSEs takenvs objmanip | 1671 | 0.19 | 0.15 | 0.24 | 0.02 | 8.49 | 4.33E-17 | 2.09E-16 | *** | 0.04 | 0.04 | 0.92 | 72.16 | 1 | 1669 |
| Number of humanities GCSEs takenvs vis | 1671 | 0.20 | 0.15 | 0.24 | 0.02 | 8.78 | 4.11E-18 | 2.12E-17 | *** | 0.04 | 0.04 | 0.92 | 77.01 | 1 | 1669 |
| Number of humanities GCSEs takenvs nav | 1671 | 0.19 | 0.15 | 0.24 | 0.02 | 8.44 | 6.64E-17 | 3.06E-16 | *** | 0.04 | 0.04 | 0.92 | 71.28 | 1 | 1669 |
| Number of humanities GCSEs takenvs spatab | 1671 | 0.20 | 0.15 | 0.24 | 0.02 | 8.78 | 4.11E-18 | 2.12E-17 | *** | 0.04 | 0.04 | 0.92 | 77.01 | 1 | 1669 |
| Number of humanities GCSEs takenvs objmanip.g | 1322 | -0.03 | -0.08 | 0.02 | 0.03 | -1.23 | 2.18E-01 | 2.74E-01 | | 0.00 | 0.00 | 0.95 | 1.52 | 1 | 1320 |
| Number of humanities GCSEs takenvs vis.g | 1322 | -0.02 | -0.07 | 0.04 | 0.03 | -0.59 | 5.54E-01 | 6.20E-01 | | 0.00 | 0.00 | 0.95 | 0.35 | 1 | 1320 |
| Number of humanities GCSEs takenvs nav.g | 1322 | -0.01 | -0.07 | 0.04 | 0.03 | -0.53 | 5.95E-01 | 6.61E-01 | | 0.00 | 0.00 | 0.95 | 0.28 | 1 | 1320 |
| Number of humanities GCSEs takenvs spatab.g | 1322 | -0.02 | -0.07 | 0.04 | 0.03 | -0.59 | 5.54E-01 | 6.20E-01 | | 0.00 | 0.00 | 0.95 | 0.35 | 1 | 1320 |
| Number of humanities GCSEs takenvs objmanip.v | 1324 | 0.02 | -0.03 | 0.07 | 0.03 | 0.85 | 3.98E-01 | 4.59E-01 | | 0.00 | 0.00 | 0.95 | 0.72 | 1 | 1322 |
| Number of humanities GCSEs takenvs vis.v | 1324 | 0.03 | -0.02 | 0.08 | 0.03 | 1.26 | 2.08E-01 | 2.65E-01 | | 0.00 | 0.00 | 0.95 | 1.58 | 1 | 1322 |
| Number of humanities GCSEs takenvs nav.v | 1324 | 0.03 | -0.02 | 0.08 | 0.03 | 1.08 | 2.82E-01 | 3.46E-01 | | 0.00 | 0.00 | 0.95 | 1.16 | 1 | 1322 |
| Number of humanities GCSEs takenvs spatab.v | 1324 | 0.03 | -0.02 | 0.08 | 0.03 | 1.26 | 2.08E-01 | 2.65E-01 | | 0.00 | 0.00 | 0.95 | 1.58 | 1 | 1322 |
| Number of humanities A(S)-Levels takenvs objmanip | 1703 | 0.07 | 0.02 | 0.12 | 0.03 | 2.75 | 6.05E-03 | 1.13E-02 | * | 0.00 | 0.00 | 1.07 | 7.55 | 1 | 1701 |
| Number of humanities A(S)-Levels takenvs vis | 1703 | 0.09 | 0.03 | 0.14 | 0.03 | 3.30 | 9.71E-04 | 1.99E-03 | ** | 0.01 | 0.01 | 1.07 | 10.92 | 1 | 1701 |
| Number of humanities A(S)-Levels takenvs nav | 1703 | 0.09 | 0.04 | 0.14 | 0.03 | 3.54 | 4.04E-04 | 8.87E-04 | *** | 0.01 | 0.01 | 1.07 | 12.56 | 1 | 1701 |
| Number of humanities A(S)-Levels takenvs spatab | 1703 | 0.09 | 0.03 | 0.14 | 0.03 | 3.30 | 9.71E-04 | 1.99E-03 | ** | 0.01 | 0.01 | 1.07 | 10.92 | 1 | 1701 |
| Number of humanities A(S)-Levels takenvs objmanip.g | 1290 | -0.12 | -0.18 | -0.06 | 0.03 | -4.01 | 6.38E-05 | 1.50E-04 | *** | 0.01 | 0.01 | 1.08 | 16.09 | 1 | 1288 |
| Number of humanities A(S)-Levels takenvs vis.g | 1290 | -0.09 | -0.15 | -0.04 | 0.03 | -3.14 | 1.72E-03 | 3.42E-03 | ** | 0.01 | 0.01 | 1.08 | 9.87 | 1 | 1288 |
| Number of humanities A(S)-Levels takenvs nav.g | 1290 | -0.08 | -0.14 | -0.02 | 0.03 | -2.57 | 1.03E-02 | 1.84E-02 | * | 0.01 | 0.00 | 1.09 | 6.60 | 1 | 1288 |
| Number of humanities A(S)-Levels takenvs spatab.g | 1290 | -0.09 | -0.15 | -0.04 | 0.03 | -3.14 | 1.72E-03 | 3.42E-03 | ** | 0.01 | 0.01 | 1.08 | 9.87 | 1 | 1288 |
| Number of humanities A(S)-Levels takenvs objmanip.v | 1291 | -0.09 | -0.15 | -0.03 | 0.03 | -2.90 | 3.74E-03 | 7.20E-03 | ** | 0.01 | 0.01 | 1.09 | 8.44 | 1 | 1289 |
| Number of humanities A(S)-Levels takenvs vis.v | 1291 | -0.07 | -0.13 | -0.01 | 0.03 | -2.24 | 2.55E-02 | 4.13E-02 | * | 0.00 | 0.00 | 1.09 | 5.00 | 1 | 1289 |
| Number of humanities A(S)-Levels takenvs nav.v | 1291 | -0.06 | -0.12 | 0.00 | 0.03 | -1.88 | 5.99E-02 | 8.86E-02 | | 0.00 | 0.00 | 1.09 | 3.55 | 1 | 1289 |
| Number of humanities A(S)-Levels takenvs spatab.v | 1291 | -0.07 | -0.13 | -0.01 | 0.03 | -2.24 | 2.55E-02 | 4.13E-02 | * | 0.00 | 0.00 | 1.09 | 5.00 | 1 | 1289 |
| GCSE STEM + humanities mean gradevs objmanip | 1669 | 0.42 | 0.38 | 0.46 | 0.02 | 20.46 | 3.67E-83 | 1.18E-81 | *** | 0.20 | 0.20 | 0.83 | 418.47 | 1 | 1667 |
| GCSE STEM + humanities mean gradevs vis | 1669 | 0.41 | 0.37 | 0.45 | 0.02 | 19.64 | 2.03E-77 | 5.52E-76 | *** | 0.19 | 0.19 | 0.84 | 385.73 | 1 | 1667 |
| GCSE STEM + humanities mean gradevs nav | 1669 | 0.39 | 0.35 | 0.43 | 0.02 | 18.59 | 2.79E-70 | 5.54E-69 | *** | 0.17 | 0.17 | 0.84 | 345.75 | 1 | 1667 |
| GCSE STEM + humanities mean gradevs spatab | 1669 | 0.41 | 0.37 | 0.45 | 0.02 | 19.64 | 2.03E-77 | 5.52E-76 | *** | 0.19 | 0.19 | 0.84 | 385.73 | 1 | 1667 |
| GCSE STEM + humanities mean gradevs objmanip.g | 1320 | 0.08 | 0.03 | 0.13 | 0.03 | 3.15 | 1.67E-03 | 3.36E-03 | ** | 0.01 | 0.01 | 0.92 | 9.92 | 1 | 1318 |
| GCSE STEM + humanities mean gradevs vis.g | 1320 | 0.07 | 0.02 | 0.12 | 0.03 | 2.77 | 5.69E-03 | 1.07E-02 | * | 0.01 | 0.01 | 0.92 | 7.67 | 1 | 1318 |
| GCSE STEM + humanities mean gradevs nav.g | 1320 | 0.06 | 0.01 | 0.11 | 0.03 | 2.41 | 1.59E-02 | 2.71E-02 | * | 0.00 | 0.00 | 0.92 | 5.83 | 1 | 1318 |
| GCSE STEM + humanities mean gradevs spatab.g | 1320 | 0.07 | 0.02 | 0.12 | 0.03 | 2.77 | 5.69E-03 | 1.07E-02 | * | 0.01 | 0.01 | 0.92 | 7.67 | 1 | 1318 |
| GCSE STEM + humanities mean gradevs objmanip.v | 1322 | 0.19 | 0.14 | 0.24 | 0.02 | 7.48 | 1.35E-13 | 5.54E-13 | *** | 0.04 | 0.04 | 0.91 | 55.95 | 1 | 1320 |
| GCSE STEM + humanities mean gradevs vis.v | 1322 | 0.17 | 0.12 | 0.22 | 0.02 | 6.89 | 8.51E-12 | 3.18E-11 | *** | 0.03 | 0.03 | 0.91 | 47.50 | 1 | 1320 |

| | | | | | | | | | | | | | | | |
|---|---|---|---|---|---|---|---|---|---|---|---|---|---|---|---|
| GCSE STEM + humanities mean gradevs nav.v | 1322 | 0.15 | 0.10 | 0.20 | 0.03 | 6.15 | 1.00E-09 | 3.29E-09 | *** | 0.03 | 0.03 | 0.91 | 37.87 | 1 | 1320 |
| GCSE STEM + humanities mean gradevs spatab.v | 1322 | 0.17 | 0.12 | 0.22 | 0.02 | 6.89 | 8.51E-12 | 3.18E-11 | *** | 0.03 | 0.03 | 0.91 | 47.50 | 1 | 1320 |
| A(S)-Level STEM + humanities mean gradevs objmanip | 1199 | 0.17 | 0.11 | 0.23 | 0.03 | 5.64 | 2.13E-08 | 6.17E-08 | *** | 0.03 | 0.03 | 0.98 | 31.80 | 1 | 1197 |
| A(S)-Level STEM + humanities mean gradevs vis | 1199 | 0.16 | 0.10 | 0.22 | 0.03 | 5.10 | 3.89E-07 | 1.07E-06 | *** | 0.02 | 0.02 | 0.99 | 26.04 | 1 | 1197 |
| A(S)-Level STEM + humanities mean gradevs nav | 1199 | 0.15 | 0.09 | 0.21 | 0.03 | 4.83 | 1.51E-06 | 4.06E-06 | *** | 0.02 | 0.02 | 0.99 | 23.37 | 1 | 1197 |
| A(S)-Level STEM + humanities mean gradevs spatab | 1199 | 0.16 | 0.10 | 0.22 | 0.03 | 5.10 | 3.89E-07 | 1.07E-06 | *** | 0.02 | 0.02 | 0.99 | 26.04 | 1 | 1197 |
| A(S)-Level STEM + humanities mean gradevs objmanip.g | 944 | -0.05 | -0.12 | 0.01 | 0.03 | -1.56 | 1.18E-01 | 1.57E-01 | | 0.00 | 0.00 | 0.99 | 2.45 | 1 | 942 |
| A(S)-Level STEM + humanities mean gradevs vis.g | 944 | -0.07 | -0.13 | 0.00 | 0.03 | -2.03 | 4.26E-02 | 6.49E-02 | | 0.00 | 0.00 | 0.99 | 4.12 | 1 | 942 |
| A(S)-Level STEM + humanities mean gradevs nav.g | 944 | -0.07 | -0.13 | 0.00 | 0.03 | -2.07 | 3.84E-02 | 5.93E-02 | | 0.00 | 0.00 | 0.99 | 4.30 | 1 | 942 |
| A(S)-Level STEM + humanities mean gradevs spatab.g | 944 | -0.07 | -0.13 | 0.00 | 0.03 | -2.03 | 4.26E-02 | 6.49E-02 | | 0.00 | 0.00 | 0.99 | 4.12 | 1 | 942 |
| A(S)-Level STEM + humanities mean gradevs objmanip.v | 945 | 0.00 | -0.06 | 0.06 | 0.03 | 0.01 | 9.95E-01 | 9.95E-01 | | 0.00 | 0.00 | 0.99 | 0.00 | 1 | 943 |
| A(S)-Level STEM + humanities mean gradevs vis.v | 945 | -0.02 | -0.08 | 0.05 | 0.03 | -0.51 | 6.11E-01 | 6.69E-01 | | 0.00 | 0.00 | 0.99 | 0.26 | 1 | 943 |
| A(S)-Level STEM + humanities mean gradevs nav.v | 945 | -0.02 | -0.09 | 0.04 | 0.03 | -0.74 | 4.59E-01 | 5.25E-01 | | 0.00 | 0.00 | 0.99 | 0.55 | 1 | 943 |
| A(S)-Level STEM + humanities mean gradevs spatab.v | 945 | -0.02 | -0.08 | 0.05 | 0.03 | -0.51 | 6.11E-01 | 6.69E-01 | | 0.00 | 0.00 | 0.99 | 0.26 | 1 | 943 |
| GCSE Core mean grade vs objmanip | 1667 | 0.43 | 0.39 | 0.47 | 0.02 | 21.38 | 7.86E-90 | 3.38E-88 | *** | 0.22 | 0.21 | 0.82 | 457.24 | 1 | 1665 |
| GCSE Core mean grade vs vis | 1667 | 0.42 | 0.38 | 0.46 | 0.02 | 20.59 | 3.92E-84 | 1.35E-82 | *** | 0.20 | 0.20 | 0.82 | 424.13 | 1 | 1665 |
| GCSE Core mean grade vs nav | 1667 | 0.41 | 0.37 | 0.45 | 0.02 | 19.58 | 5.61E-77 | 1.38E-75 | *** | 0.19 | 0.19 | 0.83 | 383.28 | 1 | 1665 |
| GCSE Core mean grade vs spatab | 1667 | 0.42 | 0.38 | 0.46 | 0.02 | 20.59 | 3.92E-84 | 1.35E-82 | *** | 0.20 | 0.20 | 0.82 | 424.13 | 1 | 1665 |
| GCSE Core mean grade vs objmanip.g | 1318 | 0.09 | 0.04 | 0.14 | 0.03 | 3.63 | 2.91E-04 | 6.53E-04 | *** | 0.01 | 0.01 | 0.92 | 13.20 | 1 | 1316 |
| GCSE Core mean grade vs vis.g | 1318 | 0.08 | 0.03 | 0.13 | 0.03 | 3.28 | 1.07E-03 | 2.18E-03 | ** | 0.01 | 0.01 | 0.92 | 10.74 | 1 | 1316 |
| GCSE Core mean grade vs nav.g | 1318 | 0.08 | 0.03 | 0.12 | 0.03 | 2.97 | 3.06E-03 | 5.96E-03 | ** | 0.01 | 0.01 | 0.92 | 8.80 | 1 | 1316 |
| GCSE Core mean grade vs spatab.g | 1318 | 0.08 | 0.03 | 0.13 | 0.03 | 3.28 | 1.07E-03 | 2.18E-03 | ** | 0.01 | 0.01 | 0.92 | 10.74 | 1 | 1316 |
| GCSE Core mean grade vs objmanip.v | 1320 | 0.20 | 0.15 | 0.25 | 0.02 | 7.97 | 3.52E-15 | 1.55E-14 | *** | 0.05 | 0.05 | 0.90 | 63.46 | 1 | 1318 |
| GCSE Core mean grade vs vis.v | 1320 | 0.18 | 0.14 | 0.23 | 0.02 | 7.40 | 2.47E-13 | 9.86E-13 | *** | 0.04 | 0.04 | 0.91 | 54.72 | 1 | 1318 |
| GCSE Core mean grade vs nav.v | 1320 | 0.17 | 0.12 | 0.22 | 0.02 | 6.70 | 3.07E-11 | 1.10E-10 | *** | 0.03 | 0.03 | 0.91 | 44.90 | 1 | 1318 |
| GCSE Core mean grade vs spatab.v | 1320 | 0.18 | 0.14 | 0.23 | 0.02 | 7.40 | 2.47E-13 | 9.86E-13 | *** | 0.04 | 0.04 | 0.91 | 54.72 | 1 | 1318 |
| GCSE overall mean grade vs objmanip | 1669 | 0.41 | 0.36 | 0.45 | 0.02 | 19.61 | 3.55E-77 | 9.15E-76 | *** | 0.19 | 0.19 | 0.84 | 384.36 | 1 | 1667 |
| GCSE overall mean grade vs vis | 1669 | 0.39 | 0.35 | 0.44 | 0.02 | 18.80 | 1.23E-71 | 2.54E-70 | *** | 0.17 | 0.17 | 0.85 | 353.28 | 1 | 1667 |
| GCSE overall mean grade vs nav | 1669 | 0.38 | 0.34 | 0.42 | 0.02 | 17.76 | 9.26E-65 | 1.65E-63 | *** | 0.16 | 0.16 | 0.86 | 315.38 | 1 | 1667 |
| GCSE overall mean grade vs spatab | 1669 | 0.39 | 0.35 | 0.44 | 0.02 | 18.80 | 1.23E-71 | 2.54E-70 | *** | 0.17 | 0.17 | 0.85 | 353.28 | 1 | 1667 |
| GCSE overall mean grade vs objmanip.g | 1320 | 0.07 | 0.02 | 0.12 | 0.03 | 2.74 | 6.16E-03 | 1.15E-02 | * | 0.01 | 0.00 | 0.93 | 7.53 | 1 | 1318 |
| GCSE overall mean grade vs vis.g | 1320 | 0.06 | 0.01 | 0.11 | 0.03 | 2.34 | 1.95E-02 | 3.22E-02 | * | 0.00 | 0.00 | 0.93 | 5.47 | 1 | 1318 |
| GCSE overall mean grade vs nav.g | 1320 | 0.05 | 0.00 | 0.10 | 0.03 | 1.95 | 5.13E-02 | 7.70E-02 | | 0.00 | 0.00 | 0.93 | 3.80 | 1 | 1318 |
| GCSE overall mean grade vs spatab.g | 1320 | 0.06 | 0.01 | 0.11 | 0.03 | 2.34 | 1.95E-02 | 3.22E-02 | * | 0.00 | 0.00 | 0.93 | 5.47 | 1 | 1318 |
| GCSE overall mean grade vs objmanip.v | 1322 | 0.17 | 0.13 | 0.22 | 0.03 | 6.93 | 6.78E-12 | 2.59E-11 | *** | 0.04 | 0.03 | 0.92 | 47.96 | 1 | 1320 |
| GCSE overall mean grade vs vis.v | 1322 | 0.16 | 0.11 | 0.21 | 0.03 | 6.32 | 3.54E-10 | 1.21E-09 | *** | 0.03 | 0.03 | 0.92 | 39.96 | 1 | 1320 |
| GCSE overall mean grade vs nav.v | 1322 | 0.14 | 0.09 | 0.19 | 0.03 | 5.56 | 3.26E-08 | 9.41E-08 | *** | 0.02 | 0.02 | 0.92 | 30.91 | 1 | 1320 |

| | | | | | | | | | | | | | | | |
|---|---|---|---|---|---|---|---|---|---|---|---|---|---|---|---|
| GCSE overall mean grade vs spatab.v | 1322 | 0.16 | 0.11 | 0.21 | 0.03 | 6.32 | 3.54E-10 | 1.21E-09 | *** | 0.03 | 0.03 | 0.92 | 39.96 | 1 | 1320 |
| GCSE English mean grade vs objmanip | 1662 | 0.30 | 0.26 | 0.34 | 0.02 | 14.01 | 3.10E-42 | 3.90E-41 | *** | 0.11 | 0.11 | 0.87 | 196.29 | 1 | 1660 |
| GCSE English mean grade vs vis | 1662 | 0.30 | 0.26 | 0.34 | 0.02 | 13.72 | 1.18E-40 | 1.35E-39 | *** | 0.10 | 0.10 | 0.87 | 188.21 | 1 | 1660 |
| GCSE English mean grade vs nav | 1662 | 0.29 | 0.24 | 0.33 | 0.02 | 13.02 | 5.75E-37 | 5.70E-36 | *** | 0.09 | 0.09 | 0.88 | 169.50 | 1 | 1660 |
| GCSE English mean grade vs spatab | 1662 | 0.30 | 0.26 | 0.34 | 0.02 | 13.72 | 1.18E-40 | 1.35E-39 | *** | 0.10 | 0.10 | 0.87 | 188.21 | 1 | 1660 |
| GCSE English mean grade vs objmanip.g | 1313 | -0.02 | -0.07 | 0.03 | 0.03 | -0.89 | 3.75E-01 | 4.34E-01 | | 0.00 | 0.00 | 0.91 | 0.79 | 1 | 1311 |
| GCSE English mean grade vs vis.g | 1313 | -0.02 | -0.07 | 0.03 | 0.03 | -0.89 | 3.74E-01 | 4.34E-01 | | 0.00 | 0.00 | 0.91 | 0.79 | 1 | 1311 |
| GCSE English mean grade vs nav.g | 1313 | -0.03 | -0.07 | 0.02 | 0.03 | -1.00 | 3.16E-01 | 3.80E-01 | | 0.00 | 0.00 | 0.91 | 1.01 | 1 | 1311 |
| GCSE English mean grade vs spatab.g | 1313 | -0.02 | -0.07 | 0.03 | 0.03 | -0.89 | 3.74E-01 | 4.34E-01 | | 0.00 | 0.00 | 0.91 | 0.79 | 1 | 1311 |
| GCSE English mean grade vs objmanip.v | 1315 | 0.06 | 0.01 | 0.11 | 0.03 | 2.43 | 1.50E-02 | 2.58E-02 | * | 0.00 | 0.00 | 0.91 | 5.93 | 1 | 1313 |
| GCSE English mean grade vs vis.v | 1315 | 0.06 | 0.01 | 0.11 | 0.03 | 2.22 | 2.66E-02 | 4.29E-02 | * | 0.00 | 0.00 | 0.91 | 4.93 | 1 | 1313 |
| GCSE English mean grade vs nav.v | 1315 | 0.04 | -0.01 | 0.09 | 0.03 | 1.75 | 8.00E-02 | 1.13E-01 | | 0.00 | 0.00 | 0.91 | 3.07 | 1 | 1313 |
| GCSE English mean grade vs spatab.v | 1315 | 0.06 | 0.01 | 0.11 | 0.03 | 2.22 | 2.66E-02 | 4.29E-02 | * | 0.00 | 0.00 | 0.91 | 4.93 | 1 | 1313 |
| GCSE Technology mean grade vs objmanip | 963 | 0.31 | 0.25 | 0.37 | 0.03 | 10.07 | 9.56E-23 | 6.58E-22 | *** | 0.10 | 0.09 | 0.92 | 101.42 | 1 | 961 |
| GCSE Technology mean grade vs vis | 963 | 0.30 | 0.24 | 0.36 | 0.03 | 9.52 | 1.32E-20 | 7.67E-20 | *** | 0.09 | 0.09 | 0.93 | 90.68 | 1 | 961 |
| GCSE Technology mean grade vs nav | 963 | 0.28 | 0.21 | 0.34 | 0.03 | 8.79 | 6.90E-18 | 3.39E-17 | *** | 0.07 | 0.07 | 0.93 | 77.23 | 1 | 961 |
| GCSE Technology mean grade vs spatab | 963 | 0.30 | 0.24 | 0.36 | 0.03 | 9.52 | 1.32E-20 | 7.67E-20 | *** | 0.09 | 0.09 | 0.93 | 90.68 | 1 | 961 |
| GCSE Technology mean grade vs objmanip.g | 765 | 0.07 | 0.00 | 0.14 | 0.03 | 2.06 | 3.96E-02 | 6.09E-02 | | 0.01 | 0.00 | 0.95 | 4.25 | 1 | 763 |
| GCSE Technology mean grade vs vis.g | 765 | 0.06 | -0.01 | 0.13 | 0.04 | 1.70 | 9.01E-02 | 1.25E-01 | | 0.00 | 0.00 | 0.95 | 2.88 | 1 | 763 |
| GCSE Technology mean grade vs nav.g | 765 | 0.05 | -0.02 | 0.11 | 0.03 | 1.29 | 1.96E-01 | 2.54E-01 | | 0.00 | 0.00 | 0.95 | 1.67 | 1 | 763 |
| GCSE Technology mean grade vs spatab.g | 765 | 0.06 | -0.01 | 0.13 | 0.04 | 1.70 | 9.01E-02 | 1.25E-01 | | 0.00 | 0.00 | 0.95 | 2.88 | 1 | 763 |
| GCSE Technology mean grade vs objmanip.v | 766 | 0.15 | 0.09 | 0.22 | 0.03 | 4.43 | 1.08E-05 | 2.74E-05 | *** | 0.03 | 0.02 | 0.94 | 19.62 | 1 | 764 |
| GCSE Technology mean grade vs vis.v | 766 | 0.14 | 0.07 | 0.21 | 0.03 | 3.99 | 7.37E-05 | 1.72E-04 | *** | 0.02 | 0.02 | 0.94 | 15.89 | 1 | 764 |
| GCSE Technology mean grade vs nav.v | 766 | 0.12 | 0.05 | 0.19 | 0.03 | 3.39 | 7.25E-04 | 1.52E-03 | ** | 0.01 | 0.01 | 0.94 | 11.52 | 1 | 764 |
| GCSE Technology mean grade vs spatab.v | 766 | 0.14 | 0.07 | 0.21 | 0.03 | 3.99 | 7.37E-05 | 1.72E-04 | *** | 0.02 | 0.02 | 0.94 | 15.89 | 1 | 764 |
| GCSE Humanities mean grade vs objmanip | 1616 | 0.30 | 0.26 | 0.35 | 0.02 | 13.72 | 1.26E-40 | 1.41E-39 | *** | 0.10 | 0.10 | 0.88 | 188.36 | 1 | 1614 |
| GCSE Humanities mean grade vs vis | 1616 | 0.29 | 0.25 | 0.34 | 0.02 | 13.18 | 8.96E-38 | 9.43E-37 | *** | 0.10 | 0.10 | 0.88 | 173.83 | 1 | 1614 |
| GCSE Humanities mean grade vs nav | 1616 | 0.28 | 0.24 | 0.33 | 0.02 | 12.41 | 7.82E-34 | 7.20E-33 | *** | 0.09 | 0.09 | 0.89 | 153.96 | 1 | 1614 |
| GCSE Humanities mean grade vs spatab | 1616 | 0.29 | 0.25 | 0.34 | 0.02 | 13.18 | 8.96E-38 | 9.43E-37 | *** | 0.10 | 0.10 | 0.88 | 173.83 | 1 | 1614 |
| GCSE Humanities mean grade vs objmanip.g | 1279 | 0.03 | -0.02 | 0.08 | 0.03 | 1.19 | 2.33E-01 | 2.92E-01 | | 0.00 | 0.00 | 0.92 | 1.42 | 1 | 1277 |
| GCSE Humanities mean grade vs vis.g | 1279 | 0.02 | -0.03 | 0.07 | 0.03 | 0.90 | 3.67E-01 | 4.30E-01 | | 0.00 | 0.00 | 0.92 | 0.81 | 1 | 1277 |
| GCSE Humanities mean grade vs nav.g | 1279 | 0.01 | -0.04 | 0.06 | 0.03 | 0.49 | 6.24E-01 | 6.82E-01 | | 0.00 | 0.00 | 0.92 | 0.24 | 1 | 1277 |
| GCSE Humanities mean grade vs spatab.g | 1279 | 0.02 | -0.03 | 0.07 | 0.03 | 0.90 | 3.67E-01 | 4.30E-01 | | 0.00 | 0.00 | 0.92 | 0.81 | 1 | 1277 |
| GCSE Humanities mean grade vs objmanip.v | 1281 | 0.11 | 0.06 | 0.16 | 0.03 | 4.18 | 3.06E-05 | 7.37E-05 | *** | 0.01 | 0.01 | 0.92 | 17.51 | 1 | 1279 |
| GCSE Humanities mean grade vs vis.v | 1281 | 0.10 | 0.05 | 0.15 | 0.03 | 3.75 | 1.86E-04 | 4.19E-04 | *** | 0.01 | 0.01 | 0.92 | 14.05 | 1 | 1279 |
| GCSE Humanities mean grade vs nav.v | 1281 | 0.08 | 0.03 | 0.13 | 0.03 | 3.07 | 2.19E-03 | 4.33E-03 | ** | 0.01 | 0.01 | 0.92 | 9.42 | 1 | 1279 |
| GCSE Humanities mean grade vs spatab.v | 1281 | 0.10 | 0.05 | 0.15 | 0.03 | 3.75 | 1.86E-04 | 4.19E-04 | *** | 0.01 | 0.01 | 0.92 | 14.05 | 1 | 1279 |
| GCSE Languages mean grade vs objmanip | 1138 | 0.26 | 0.20 | 0.32 | 0.03 | 9.06 | 5.75E-19 | 3.09E-18 | *** | 0.07 | 0.07 | 0.93 | 82.01 | 1 | 1136 |
| GCSE Languages mean grade vs vis | 1138 | 0.25 | 0.19 | 0.31 | 0.03 | 8.52 | 5.11E-17 | 2.42E-16 | *** | 0.06 | 0.06 | 0.93 | 72.55 | 1 | 1136 |
| GCSE Languages mean grade vs nav | 1138 | 0.24 | 0.18 | 0.30 | 0.03 | 8.11 | 1.30E-15 | 5.80E-15 | *** | 0.05 | 0.05 | 0.93 | 65.77 | 1 | 1136 |
| GCSE Languages mean grade vs spatab | 1138 | 0.25 | 0.19 | 0.31 | 0.03 | 8.52 | 5.11E-17 | 2.42E-16 | *** | 0.06 | 0.06 | 0.93 | 72.55 | 1 | 1136 |
| GCSE Languages mean grade vs objmanip.g | 910 | -0.01 | -0.08 | 0.05 | 0.03 | -0.47 | 6.41E-01 | 6.98E-01 | | 0.00 | 0.00 | 0.95 | 0.22 | 1 | 908 |
| GCSE Languages mean grade vs vis.g | 910 | -0.03 | -0.09 | 0.03 | 0.03 | -0.91 | 3.65E-01 | 4.29E-01 | | 0.00 | 0.00 | 0.95 | 0.82 | 1 | 908 |

| | | | | | | | | | | | | | | | |
|---|---|---|---|---|---|---|---|---|---|---|---|---|---|---|---|
| GCSE Languages mean grade vs nav.g | 910 | -0.03 | -0.10 | 0.03 | 0.03 | -1.03 | 3.03E-01 | 3.67E-01 | | 0.00 | 0.00 | 0.95 | 1.06 | 1 | 908 |
| GCSE Languages mean grade vs spatab.g | 910 | -0.03 | -0.09 | 0.03 | 0.03 | -0.91 | 3.65E-01 | 4.29E-01 | | 0.00 | 0.00 | 0.95 | 0.82 | 1 | 908 |
| GCSE Languages mean grade vs objmanip.v | 911 | 0.07 | 0.01 | 0.13 | 0.03 | 2.19 | 2.89E-02 | 4.63E-02 | * | 0.01 | 0.00 | 0.94 | 4.79 | 1 | 909 |
| GCSE Languages mean grade vs vis.v | 911 | 0.05 | -0.01 | 0.11 | 0.03 | 1.66 | 9.69E-02 | 1.34E-01 | | 0.00 | 0.00 | 0.94 | 2.76 | 1 | 909 |
| GCSE Languages mean grade vs nav.v | 911 | 0.04 | -0.02 | 0.10 | 0.03 | 1.31 | 1.89E-01 | 2.46E-01 | | 0.00 | 0.00 | 0.95 | 1.73 | 1 | 909 |
| GCSE Languages mean grade vs spatab.v | 911 | 0.05 | -0.01 | 0.11 | 0.03 | 1.66 | 9.69E-02 | 1.34E-01 | | 0.00 | 0.00 | 0.94 | 2.76 | 1 | 909 |
| GCSE Vocational mean grade vs objmanip | 658 | 0.27 | 0.19 | 0.35 | 0.04 | 6.78 | 2.66E-11 | 9.86E-11 | *** | 0.07 | 0.06 | 0.98 | 45.99 | 1 | 656 |
| GCSE Vocational mean grade vs vis | 658 | 0.26 | 0.18 | 0.34 | 0.04 | 6.30 | 5.35E-10 | 1.80E-09 | *** | 0.06 | 0.06 | 0.98 | 39.73 | 1 | 656 |
| GCSE Vocational mean grade vs nav | 658 | 0.23 | 0.15 | 0.31 | 0.04 | 5.67 | 2.10E-08 | 6.13E-08 | *** | 0.05 | 0.05 | 0.99 | 32.19 | 1 | 656 |
| GCSE Vocational mean grade vs spatab | 658 | 0.26 | 0.18 | 0.34 | 0.04 | 6.30 | 5.35E-10 | 1.80E-09 | *** | 0.06 | 0.06 | 0.98 | 39.73 | 1 | 656 |
| GCSE Vocational mean grade vs objmanip.g | 516 | 0.00 | -0.09 | 0.09 | 0.05 | 0.03 | 9.80E-01 | 9.81E-01 | | 0.00 | 0.00 | 1.03 | 0.00 | 1 | 514 |
| GCSE Vocational mean grade vs vis.g | 516 | -0.02 | -0.11 | 0.07 | 0.05 | -0.35 | 7.24E-01 | 7.74E-01 | | 0.00 | 0.00 | 1.03 | 0.12 | 1 | 514 |
| GCSE Vocational mean grade vs nav.g | 516 | -0.02 | -0.11 | 0.06 | 0.04 | -0.54 | 5.90E-01 | 6.56E-01 | | 0.00 | 0.00 | 1.03 | 0.29 | 1 | 514 |
| GCSE Vocational mean grade vs spatab.g | 516 | -0.02 | -0.11 | 0.07 | 0.05 | -0.35 | 7.24E-01 | 7.74E-01 | | 0.00 | 0.00 | 1.03 | 0.12 | 1 | 514 |
| GCSE Vocational mean grade vs objmanip.v | 517 | 0.08 | -0.01 | 0.17 | 0.05 | 1.71 | 8.78E-02 | 1.23E-01 | | 0.01 | 0.00 | 1.03 | 2.93 | 1 | 515 |
| GCSE Vocational mean grade vs vis.v | 517 | 0.06 | -0.03 | 0.15 | 0.05 | 1.28 | 2.01E-01 | 2.58E-01 | | 0.00 | 0.00 | 1.03 | 1.64 | 1 | 515 |
| GCSE Vocational mean grade vs nav.v | 517 | 0.04 | -0.05 | 0.13 | 0.04 | 0.91 | 3.64E-01 | 4.29E-01 | | 0.00 | 0.00 | 1.03 | 0.83 | 1 | 515 |
| GCSE Vocational mean grade vs spatab.v | 517 | 0.06 | -0.03 | 0.15 | 0.05 | 1.28 | 2.01E-01 | 2.58E-01 | | 0.00 | 0.00 | 1.03 | 1.64 | 1 | 515 |
| GCSE Maths grade vs objmanip | 1658 | 0.48 | 0.44 | 0.52 | 0.02 | 23.91 | 8.72E-109 | 4.50E-106 | *** | 0.26 | 0.26 | 0.80 | 571.72 | 1 | 1656 |
| GCSE Maths grade vs vis | 1658 | 0.47 | 0.43 | 0.51 | 0.02 | 22.88 | 6.52E-101 | 6.73E-99 | *** | 0.24 | 0.24 | 0.81 | 523.56 | 1 | 1656 |
| GCSE Maths grade vs nav | 1658 | 0.45 | 0.41 | 0.49 | 0.02 | 21.84 | 3.93E-93 | 2.03E-91 | *** | 0.22 | 0.22 | 0.82 | 477.00 | 1 | 1656 |
| GCSE Maths grade vs spatab | 1658 | 0.47 | 0.43 | 0.51 | 0.02 | 22.88 | 6.52E-101 | 6.73E-99 | *** | 0.24 | 0.24 | 0.81 | 523.56 | 1 | 1656 |
| GCSE Maths grade vs objmanip.g | 1310 | 0.17 | 0.12 | 0.22 | 0.03 | 6.55 | 8.40E-11 | 2.95E-10 | *** | 0.03 | 0.03 | 0.91 | 42.87 | 1 | 1308 |
| GCSE Maths grade vs vis.g | 1310 | 0.15 | 0.10 | 0.20 | 0.03 | 5.98 | 2.81E-09 | 8.90E-09 | *** | 0.03 | 0.03 | 0.92 | 35.80 | 1 | 1308 |
| GCSE Maths grade vs nav.g | 1310 | 0.14 | 0.09 | 0.19 | 0.03 | 5.45 | 5.89E-08 | 1.69E-07 | *** | 0.02 | 0.02 | 0.92 | 29.74 | 1 | 1308 |
| GCSE Maths grade vs spatab.g | 1310 | 0.15 | 0.10 | 0.20 | 0.03 | 5.98 | 2.81E-09 | 8.90E-09 | *** | 0.03 | 0.03 | 0.92 | 35.80 | 1 | 1308 |
| GCSE Maths grade vs objmanip.v | 1312 | 0.28 | 0.24 | 0.33 | 0.02 | 11.55 | 1.82E-29 | 1.65E-28 | *** | 0.09 | 0.09 | 0.88 | 133.45 | 1 | 1310 |
| GCSE Maths grade vs vis.v | 1312 | 0.27 | 0.22 | 0.31 | 0.02 | 10.78 | 5.34E-26 | 4.44E-25 | *** | 0.08 | 0.08 | 0.89 | 116.11 | 1 | 1310 |
| GCSE Maths grade vs nav.v | 1312 | 0.24 | 0.20 | 0.29 | 0.02 | 9.86 | 3.65E-22 | 2.48E-21 | *** | 0.07 | 0.07 | 0.90 | 97.18 | 1 | 1310 |
| GCSE Maths grade vs spatab.v | 1312 | 0.27 | 0.22 | 0.31 | 0.02 | 10.78 | 5.34E-26 | 4.44E-25 | *** | 0.08 | 0.08 | 0.89 | 116.11 | 1 | 1310 |
| GCSE Core Science grade vs objmanip | 878 | 0.41 | 0.35 | 0.47 | 0.03 | 13.37 | 3.06E-37 | 3.10E-36 | *** | 0.17 | 0.17 | 0.89 | 178.78 | 1 | 876 |
| GCSE Core Science grade vs vis | 878 | 0.40 | 0.34 | 0.47 | 0.03 | 13.10 | 6.05E-36 | 5.78E-35 | *** | 0.16 | 0.16 | 0.89 | 171.66 | 1 | 876 |
| GCSE Core Science grade vs nav | 878 | 0.40 | 0.33 | 0.46 | 0.03 | 12.72 | 3.96E-34 | 3.71E-33 | *** | 0.16 | 0.15 | 0.89 | 161.76 | 1 | 876 |
| GCSE Core Science grade vs spatab | 878 | 0.40 | 0.34 | 0.47 | 0.03 | 13.10 | 6.05E-36 | 5.78E-35 | *** | 0.16 | 0.16 | 0.89 | 171.66 | 1 | 876 |
| GCSE Core Science grade vs objmanip.g | 692 | 0.09 | 0.02 | 0.17 | 0.04 | 2.46 | 1.40E-02 | 2.44E-02 | * | 0.01 | 0.01 | 0.99 | 6.08 | 1 | 690 |
| GCSE Core Science grade vs vis.g | 692 | 0.10 | 0.02 | 0.17 | 0.04 | 2.58 | 1.00E-02 | 1.79E-02 | * | 0.01 | 0.01 | 0.99 | 6.67 | 1 | 690 |
| GCSE Core Science grade vs nav.g | 692 | 0.10 | 0.03 | 0.18 | 0.04 | 2.71 | 6.93E-03 | 1.28E-02 | * | 0.01 | 0.01 | 0.99 | 7.33 | 1 | 690 |
| GCSE Core Science grade vs spatab.g | 692 | 0.10 | 0.02 | 0.17 | 0.04 | 2.58 | 1.00E-02 | 1.79E-02 | * | 0.01 | 0.01 | 0.99 | 6.67 | 1 | 690 |
| GCSE Core Science grade vs objmanip.v | 692 | 0.18 | 0.11 | 0.26 | 0.04 | 4.94 | 9.87E-07 | 2.69E-06 | *** | 0.03 | 0.03 | 0.98 | 24.39 | 1 | 690 |
| GCSE Core Science grade vs vis.v | 692 | 0.18 | 0.11 | 0.25 | 0.04 | 4.89 | 1.28E-06 | 3.46E-06 | *** | 0.03 | 0.03 | 0.98 | 23.87 | 1 | 690 |
| GCSE Core Science grade vs nav.v | 692 | 0.18 | 0.10 | 0.25 | 0.04 | 4.77 | 2.24E-06 | 5.96E-06 | *** | 0.03 | 0.03 | 0.98 | 22.76 | 1 | 690 |
| GCSE Core Science grade vs spatab.v | 692 | 0.18 | 0.11 | 0.25 | 0.04 | 4.89 | 1.28E-06 | 3.46E-06 | *** | 0.03 | 0.03 | 0.98 | 23.87 | 1 | 690 |
| GCSE Statistics grade vs objmanip | 226 | 0.25 | 0.11 | 0.38 | 0.07 | 3.55 | 4.78E-04 | 1.03E-03 | ** | 0.05 | 0.05 | 0.99 | 12.57 | 1 | 224 |

| | | | | | | | | | | | | | | | |
|---|---|---|---|---|---|---|---|---|---|---|---|---|---|---|---|
| GCSE Statistics grade vs vis | 226 | 0.24 | 0.10 | 0.37 | 0.07 | 3.50 | 5.69E-04 | 1.21E-03 | ** | 0.05 | 0.05 | 0.99 | 12.22 | 1 | 224 |
| GCSE Statistics grade vs nav | 226 | 0.23 | 0.09 | 0.36 | 0.07 | 3.36 | 9.04E-04 | 1.87E-03 | ** | 0.05 | 0.04 | 0.99 | 11.32 | 1 | 224 |
| GCSE Statistics grade vs spatab | 226 | 0.24 | 0.10 | 0.37 | 0.07 | 3.50 | 5.69E-04 | 1.21E-03 | ** | 0.05 | 0.05 | 0.99 | 12.22 | 1 | 224 |
| GCSE Statistics grade vs objmanip.g | 188 | 0.02 | -0.13 | 0.16 | 0.07 | 0.23 | 8.15E-01 | 8.45E-01 | | 0.00 | -0.01 | 1.00 | 0.05 | 1 | 186 |
| GCSE Statistics grade vs vis.g | 188 | 0.02 | -0.12 | 0.16 | 0.07 | 0.33 | 7.43E-01 | 7.89E-01 | | 0.00 | 0.00 | 1.00 | 0.11 | 1 | 186 |
| GCSE Statistics grade vs nav.g | 188 | 0.03 | -0.10 | 0.16 | 0.07 | 0.44 | 6.59E-01 | 7.13E-01 | | 0.00 | 0.00 | 1.00 | 0.20 | 1 | 186 |
| GCSE Statistics grade vs spatab.g | 188 | 0.02 | -0.12 | 0.16 | 0.07 | 0.33 | 7.43E-01 | 7.89E-01 | | 0.00 | 0.00 | 1.00 | 0.11 | 1 | 186 |
| GCSE Statistics grade vs objmanip.v | 188 | 0.07 | -0.07 | 0.22 | 0.07 | 1.00 | 3.19E-01 | 3.83E-01 | | 0.01 | 0.00 | 1.00 | 1.00 | 1 | 186 |
| GCSE Statistics grade vs vis.v | 188 | 0.07 | -0.07 | 0.21 | 0.07 | 1.02 | 3.08E-01 | 3.72E-01 | | 0.01 | 0.00 | 1.00 | 1.04 | 1 | 186 |
| GCSE Statistics grade vs nav.v | 188 | 0.07 | -0.06 | 0.20 | 0.07 | 1.04 | 3.01E-01 | 3.65E-01 | | 0.01 | 0.00 | 1.00 | 1.08 | 1 | 186 |
| GCSE Statistics grade vs spatab.v | 188 | 0.07 | -0.07 | 0.21 | 0.07 | 1.02 | 3.08E-01 | 3.72E-01 | | 0.01 | 0.00 | 1.00 | 1.04 | 1 | 186 |
| GCSE ICT grade vs objmanip | 355 | 0.31 | 0.21 | 0.41 | 0.05 | 6.09 | 3.02E-09 | 9.51E-09 | *** | 0.09 | 0.09 | 0.90 | 37.04 | 1 | 353 |
| GCSE ICT grade vs vis | 355 | 0.30 | 0.20 | 0.40 | 0.05 | 5.98 | 5.57E-09 | 1.73E-08 | *** | 0.09 | 0.09 | 0.90 | 35.72 | 1 | 353 |
| GCSE ICT grade vs nav | 355 | 0.30 | 0.19 | 0.40 | 0.05 | 5.75 | 1.92E-08 | 5.69E-08 | *** | 0.09 | 0.08 | 0.91 | 33.08 | 1 | 353 |
| GCSE ICT grade vs spatab | 355 | 0.30 | 0.20 | 0.40 | 0.05 | 5.98 | 5.57E-09 | 1.73E-08 | *** | 0.09 | 0.09 | 0.90 | 35.72 | 1 | 353 |
| GCSE ICT grade vs objmanip.g | 283 | 0.07 | -0.04 | 0.18 | 0.06 | 1.25 | 2.11E-01 | 2.68E-01 | | 0.01 | 0.00 | 0.92 | 1.57 | 1 | 281 |
| GCSE ICT grade vs vis.g | 283 | 0.07 | -0.04 | 0.18 | 0.06 | 1.25 | 2.13E-01 | 2.68E-01 | | 0.01 | 0.00 | 0.92 | 1.56 | 1 | 281 |
| GCSE ICT grade vs nav.g | 283 | 0.07 | -0.04 | 0.18 | 0.06 | 1.23 | 2.21E-01 | 2.77E-01 | | 0.01 | 0.00 | 0.92 | 1.51 | 1 | 281 |
| GCSE ICT grade vs spatab.g | 283 | 0.07 | -0.04 | 0.18 | 0.06 | 1.25 | 2.13E-01 | 2.68E-01 | | 0.01 | 0.00 | 0.92 | 1.56 | 1 | 281 |
| GCSE ICT grade vs objmanip.v | 283 | 0.16 | 0.05 | 0.27 | 0.06 | 2.80 | 5.39E-03 | 1.02E-02 | * | 0.03 | 0.02 | 0.91 | 7.87 | 1 | 281 |
| GCSE ICT grade vs vis.v | 283 | 0.15 | 0.04 | 0.26 | 0.06 | 2.72 | 6.94E-03 | 1.28E-02 | * | 0.03 | 0.02 | 0.91 | 7.40 | 1 | 281 |
| GCSE ICT grade vs nav.v | 283 | 0.15 | 0.03 | 0.26 | 0.06 | 2.58 | 1.04E-02 | 1.85E-02 | * | 0.02 | 0.02 | 0.91 | 6.66 | 1 | 281 |
| GCSE ICT grade vs spatab.v | 283 | 0.15 | 0.04 | 0.26 | 0.06 | 2.72 | 6.94E-03 | 1.28E-02 | * | 0.03 | 0.02 | 0.91 | 7.40 | 1 | 281 |
| GCSE Additional Science grade vs objmanip | 752 | 0.36 | 0.29 | 0.43 | 0.04 | 10.23 | 4.36E-23 | 3.04E-22 | *** | 0.12 | 0.12 | 0.92 | 104.67 | 1 | 750 |
| GCSE Additional Science grade vs vis | 752 | 0.35 | 0.28 | 0.42 | 0.04 | 9.85 | 1.31E-21 | 8.69E-21 | *** | 0.11 | 0.11 | 0.93 | 97.02 | 1 | 750 |
| GCSE Additional Science grade vs nav | 752 | 0.33 | 0.26 | 0.40 | 0.04 | 9.39 | 7.04E-20 | 3.95E-19 | *** | 0.11 | 0.10 | 0.93 | 88.17 | 1 | 750 |
| GCSE Additional Science grade vs spatab | 752 | 0.35 | 0.28 | 0.42 | 0.04 | 9.85 | 1.31E-21 | 8.69E-21 | *** | 0.11 | 0.11 | 0.93 | 97.02 | 1 | 750 |
| GCSE Additional Science grade vs objmanip.g | 587 | 0.07 | -0.01 | 0.15 | 0.04 | 1.67 | 9.48E-02 | 1.31E-01 | | 0.00 | 0.00 | 0.99 | 2.80 | 1 | 585 |
| GCSE Additional Science grade vs vis.g | 587 | 0.07 | -0.01 | 0.14 | 0.04 | 1.61 | 1.09E-01 | 1.47E-01 | | 0.00 | 0.00 | 0.99 | 2.58 | 1 | 585 |
| GCSE Additional Science grade vs nav.g | 587 | 0.06 | -0.02 | 0.14 | 0.04 | 1.52 | 1.28E-01 | 1.69E-01 | | 0.00 | 0.00 | 0.99 | 2.32 | 1 | 585 |
| GCSE Additional Science grade vs spatab.g | 587 | 0.07 | -0.01 | 0.14 | 0.04 | 1.61 | 1.09E-01 | 1.47E-01 | | 0.00 | 0.00 | 0.99 | 2.58 | 1 | 585 |
| GCSE Additional Science grade vs objmanip.v | 587 | 0.16 | 0.08 | 0.23 | 0.04 | 3.84 | 1.39E-04 | 3.16E-04 | *** | 0.02 | 0.02 | 0.98 | 14.72 | 1 | 585 |
| GCSE Additional Science grade vs vis.v | 587 | 0.15 | 0.07 | 0.22 | 0.04 | 3.63 | 3.10E-04 | 6.83E-04 | *** | 0.02 | 0.02 | 0.98 | 13.17 | 1 | 585 |
| GCSE Additional Science grade vs nav.v | 587 | 0.13 | 0.06 | 0.21 | 0.04 | 3.35 | 8.69E-04 | 1.80E-03 | ** | 0.02 | 0.02 | 0.98 | 11.20 | 1 | 585 |
| GCSE Additional Science grade vs spatab.v | 587 | 0.15 | 0.07 | 0.22 | 0.04 | 3.63 | 3.10E-04 | 6.83E-04 | *** | 0.02 | 0.02 | 0.98 | 13.17 | 1 | 585 |
| GCSE English Language grade vs objmanip | 1652 | 0.31 | 0.26 | 0.35 | 0.02 | 14.06 | 1.63E-42 | 2.11E-41 | *** | 0.11 | 0.11 | 0.88 | 197.79 | 1 | 1650 |
| GCSE English Language grade vs vis | 1652 | 0.31 | 0.26 | 0.35 | 0.02 | 13.86 | 2.09E-41 | 2.51E-40 | *** | 0.10 | 0.10 | 0.88 | 192.11 | 1 | 1650 |
| GCSE English Language grade vs nav | 1652 | 0.29 | 0.25 | 0.34 | 0.02 | 13.16 | 1.12E-37 | 1.16E-36 | *** | 0.09 | 0.09 | 0.88 | 173.14 | 1 | 1650 |
| GCSE English Language grade vs spatab | 1652 | 0.31 | 0.26 | 0.35 | 0.02 | 13.86 | 2.09E-41 | 2.51E-40 | *** | 0.10 | 0.10 | 0.88 | 192.11 | 1 | 1650 |
| GCSE English Language grade vs objmanip.g | 1308 | -0.02 | -0.07 | 0.03 | 0.03 | -0.75 | 4.51E-01 | 5.19E-01 | | 0.00 | 0.00 | 0.93 | 0.57 | 1 | 1306 |
| GCSE English Language grade vs vis.g | 1308 | -0.02 | -0.07 | 0.03 | 0.03 | -0.64 | 5.21E-01 | 5.87E-01 | | 0.00 | 0.00 | 0.93 | 0.41 | 1 | 1306 |
| GCSE English Language grade vs nav.g | 1308 | -0.02 | -0.07 | 0.03 | 0.03 | -0.75 | 4.55E-01 | 5.22E-01 | | 0.00 | 0.00 | 0.93 | 0.56 | 1 | 1306 |
| GCSE English Language grade vs spatab.g | 1308 | -0.02 | -0.07 | 0.03 | 0.03 | -0.64 | 5.21E-01 | 5.87E-01 | | 0.00 | 0.00 | 0.93 | 0.41 | 1 | 1306 |

| | | | | | | | | | | | | | | | |
|---|---|---|---|---|---|---|---|---|---|---|---|---|---|---|---|
| GCSE English Language grade vs objmanip.v | 1310 | 0.07 | 0.02 | 0.12 | 0.03 | 2.55 | 1.10E-02 | 1.95E-02 | * | 0.00 | 0.00 | 0.93 | 6.48 | 1 | 1308 |
| GCSE English Language grade vs vis.v | 1310 | 0.06 | 0.01 | 0.11 | 0.03 | 2.43 | 1.51E-02 | 2.58E-02 | * | 0.00 | 0.00 | 0.93 | 5.92 | 1 | 1308 |
| GCSE English Language grade vs nav.v | 1310 | 0.05 | 0.00 | 0.10 | 0.03 | 1.98 | 4.79E-02 | 7.25E-02 | | 0.00 | 0.00 | 0.93 | 3.92 | 1 | 1308 |
| GCSE English Language grade vs spatab.v | 1310 | 0.06 | 0.01 | 0.11 | 0.03 | 2.43 | 1.51E-02 | 2.58E-02 | * | 0.00 | 0.00 | 0.93 | 5.92 | 1 | 1308 |
| GCSE English Literature grade vs objmanip | 1531 | 0.23 | 0.18 | 0.28 | 0.02 | 9.57 | 4.03E-21 | 2.63E-20 | *** | 0.06 | 0.06 | 0.91 | 91.63 | 1 | 1529 |
| GCSE English Literature grade vs vis | 1531 | 0.22 | 0.18 | 0.27 | 0.02 | 9.24 | 8.02E-20 | 4.40E-19 | *** | 0.05 | 0.05 | 0.91 | 85.37 | 1 | 1529 |
| GCSE English Literature grade vs nav | 1531 | 0.21 | 0.17 | 0.26 | 0.02 | 8.76 | 5.21E-18 | 2.63E-17 | *** | 0.05 | 0.05 | 0.91 | 76.69 | 1 | 1529 |
| GCSE English Literature grade vs spatab | 1531 | 0.22 | 0.18 | 0.27 | 0.02 | 9.24 | 8.02E-20 | 4.40E-19 | *** | 0.05 | 0.05 | 0.91 | 85.37 | 1 | 1529 |
| GCSE English Literature grade vs objmanip.g | 1209 | -0.05 | -0.11 | 0.00 | 0.03 | -2.00 | 4.55E-02 | 6.90E-02 | | 0.00 | 0.00 | 0.92 | 4.01 | 1 | 1207 |
| GCSE English Literature grade vs vis.g | 1209 | -0.06 | -0.11 | 0.00 | 0.03 | -2.14 | 3.25E-02 | 5.13E-02 | | 0.00 | 0.00 | 0.92 | 4.58 | 1 | 1207 |
| GCSE English Literature grade vs nav.g | 1209 | -0.06 | -0.11 | -0.01 | 0.03 | -2.28 | 2.26E-02 | 3.72E-02 | * | 0.00 | 0.00 | 0.92 | 5.21 | 1 | 1207 |
| GCSE English Literature grade vs spatab.g | 1209 | -0.06 | -0.11 | 0.00 | 0.03 | -2.14 | 3.25E-02 | 5.13E-02 | | 0.00 | 0.00 | 0.92 | 4.58 | 1 | 1207 |
| GCSE English Literature grade vs objmanip.v | 1211 | 0.02 | -0.03 | 0.07 | 0.03 | 0.66 | 5.09E-01 | 5.77E-01 | | 0.00 | 0.00 | 0.92 | 0.44 | 1 | 1209 |
| GCSE English Literature grade vs vis.v | 1211 | 0.01 | -0.04 | 0.06 | 0.03 | 0.36 | 7.19E-01 | 7.71E-01 | | 0.00 | 0.00 | 0.92 | 0.13 | 1 | 1209 |
| GCSE English Literature grade vs nav.v | 1211 | 0.00 | -0.05 | 0.05 | 0.03 | -0.07 | 9.43E-01 | 9.53E-01 | | 0.00 | 0.00 | 0.92 | 0.01 | 1 | 1209 |
| GCSE English Literature grade vs spatab.v | 1211 | 0.01 | -0.04 | 0.06 | 0.03 | 0.36 | 7.19E-01 | 7.71E-01 | | 0.00 | 0.00 | 0.92 | 0.13 | 1 | 1209 |
| GCSE French grade vs objmanip | 653 | 0.34 | 0.27 | 0.41 | 0.04 | 9.09 | 1.14E-18 | 6.09E-18 | *** | 0.11 | 0.11 | 0.92 | 82.70 | 1 | 651 |
| GCSE French grade vs vis | 653 | 0.32 | 0.25 | 0.40 | 0.04 | 8.45 | 1.89E-16 | 8.57E-16 | *** | 0.10 | 0.10 | 0.92 | 71.41 | 1 | 651 |
| GCSE French grade vs nav | 653 | 0.31 | 0.24 | 0.39 | 0.04 | 8.02 | 4.78E-15 | 2.09E-14 | *** | 0.09 | 0.09 | 0.93 | 64.38 | 1 | 651 |
| GCSE French grade vs spatab | 653 | 0.32 | 0.25 | 0.40 | 0.04 | 8.45 | 1.89E-16 | 8.57E-16 | *** | 0.10 | 0.10 | 0.92 | 71.41 | 1 | 651 |
| GCSE French grade vs objmanip.g | 524 | 0.05 | -0.04 | 0.13 | 0.04 | 1.09 | 2.76E-01 | 3.39E-01 | | 0.00 | 0.00 | 0.94 | 1.19 | 1 | 522 |
| GCSE French grade vs vis.g | 524 | 0.02 | -0.06 | 0.10 | 0.04 | 0.46 | 6.43E-01 | 6.98E-01 | | 0.00 | 0.00 | 0.94 | 0.22 | 1 | 522 |
| GCSE French grade vs nav.g | 524 | 0.01 | -0.07 | 0.09 | 0.04 | 0.23 | 8.21E-01 | 8.49E-01 | | 0.00 | 0.00 | 0.94 | 0.05 | 1 | 522 |
| GCSE French grade vs spatab.g | 524 | 0.02 | -0.06 | 0.10 | 0.04 | 0.46 | 6.43E-01 | 6.98E-01 | | 0.00 | 0.00 | 0.94 | 0.22 | 1 | 522 |
| GCSE French grade vs objmanip.v | 525 | 0.12 | 0.04 | 0.21 | 0.04 | 2.95 | 3.35E-03 | 6.48E-03 | ** | 0.02 | 0.01 | 0.94 | 8.68 | 1 | 523 |
| GCSE French grade vs vis.v | 525 | 0.09 | 0.01 | 0.18 | 0.04 | 2.27 | 2.37E-02 | 3.88E-02 | * | 0.01 | 0.01 | 0.94 | 5.14 | 1 | 523 |
| GCSE French grade vs nav.v | 525 | 0.08 | 0.00 | 0.16 | 0.04 | 1.89 | 5.93E-02 | 8.79E-02 | | 0.01 | 0.00 | 0.94 | 3.57 | 1 | 523 |
| GCSE French grade vs spatab.v | 525 | 0.09 | 0.01 | 0.18 | 0.04 | 2.27 | 2.37E-02 | 3.88E-02 | * | 0.01 | 0.01 | 0.94 | 5.14 | 1 | 523 |
| GCSE History grade vs objmanip | 801 | 0.25 | 0.18 | 0.31 | 0.03 | 7.50 | 1.66E-13 | 6.75E-13 | *** | 0.07 | 0.06 | 0.91 | 56.30 | 1 | 799 |
| GCSE History grade vs vis | 801 | 0.23 | 0.17 | 0.30 | 0.03 | 7.09 | 2.95E-12 | 1.13E-11 | *** | 0.06 | 0.06 | 0.91 | 50.27 | 1 | 799 |
| GCSE History grade vs nav | 801 | 0.22 | 0.16 | 0.29 | 0.03 | 6.75 | 2.87E-11 | 1.04E-10 | *** | 0.05 | 0.05 | 0.92 | 45.54 | 1 | 799 |
| GCSE History grade vs spatab | 801 | 0.23 | 0.17 | 0.30 | 0.03 | 7.09 | 2.95E-12 | 1.13E-11 | *** | 0.06 | 0.06 | 0.91 | 50.27 | 1 | 799 |
| GCSE History grade vs objmanip.g | 634 | 0.00 | -0.07 | 0.08 | 0.04 | 0.07 | 9.41E-01 | 9.52E-01 | | 0.00 | 0.00 | 0.94 | 0.01 | 1 | 632 |
| GCSE History grade vs vis.g | 634 | -0.01 | -0.08 | 0.07 | 0.04 | -0.14 | 8.88E-01 | 9.02E-01 | | 0.00 | 0.00 | 0.94 | 0.02 | 1 | 632 |
| GCSE History grade vs nav.g | 634 | -0.01 | -0.08 | 0.06 | 0.04 | -0.25 | 8.01E-01 | 8.35E-01 | | 0.00 | 0.00 | 0.94 | 0.06 | 1 | 632 |
| GCSE History grade vs spatab.g | 634 | -0.01 | -0.08 | 0.07 | 0.04 | -0.14 | 8.88E-01 | 9.02E-01 | | 0.00 | 0.00 | 0.94 | 0.02 | 1 | 632 |
| GCSE History grade vs objmanip.v | 635 | 0.08 | 0.00 | 0.15 | 0.04 | 2.09 | 3.66E-02 | 5.68E-02 | | 0.01 | 0.01 | 0.94 | 4.39 | 1 | 633 |
| GCSE History grade vs vis.v | 635 | 0.07 | -0.01 | 0.14 | 0.04 | 1.77 | 7.72E-02 | 1.09E-01 | | 0.00 | 0.00 | 0.94 | 3.13 | 1 | 633 |
| GCSE History grade vs nav.v | 635 | 0.05 | -0.02 | 0.13 | 0.04 | 1.46 | 1.44E-01 | 1.89E-01 | | 0.00 | 0.00 | 0.94 | 2.14 | 1 | 633 |
| GCSE History grade vs spatab.v | 635 | 0.07 | -0.01 | 0.14 | 0.04 | 1.77 | 7.72E-02 | 1.09E-01 | | 0.00 | 0.00 | 0.94 | 3.13 | 1 | 633 |
| GCSE Spanish grade vs objmanip | 234 | 0.29 | 0.17 | 0.41 | 0.06 | 4.72 | 4.05E-06 | 1.06E-05 | *** | 0.09 | 0.08 | 0.92 | 22.29 | 1 | 232 |
| GCSE Spanish grade vs vis | 234 | 0.30 | 0.17 | 0.43 | 0.07 | 4.61 | 6.79E-06 | 1.73E-05 | *** | 0.08 | 0.08 | 0.92 | 21.21 | 1 | 232 |
| GCSE Spanish grade vs nav | 234 | 0.31 | 0.17 | 0.44 | 0.07 | 4.56 | 8.12E-06 | 2.06E-05 | *** | 0.08 | 0.08 | 0.92 | 20.84 | 1 | 232 |

| | | | | | | | | | | | | | | | |
|---|---|---|---|---|---|---|---|---|---|---|---|---|---|---|---|
| GCSE Spanish grade vs spatab | 234 | 0.30 | 0.17 | 0.43 | 0.07 | 4.61 | 6.79E-06 | 1.73E-05 | *** | 0.08 | 0.08 | 0.92 | 21.21 | 1 | 232 |
| GCSE Spanish grade vs objmanip.g | 198 | 0.03 | -0.10 | 0.16 | 0.07 | 0.44 | 6.61E-01 | 7.13E-01 | | 0.00 | 0.00 | 0.96 | 0.19 | 1 | 196 |
| GCSE Spanish grade vs vis.g | 198 | 0.02 | -0.12 | 0.16 | 0.07 | 0.27 | 7.85E-01 | 8.22E-01 | | 0.00 | 0.00 | 0.96 | 0.07 | 1 | 196 |
| GCSE Spanish grade vs nav.g | 198 | 0.02 | -0.12 | 0.16 | 0.07 | 0.33 | 7.38E-01 | 7.87E-01 | | 0.00 | 0.00 | 0.96 | 0.11 | 1 | 196 |
| GCSE Spanish grade vs spatab.g | 198 | 0.02 | -0.12 | 0.16 | 0.07 | 0.27 | 7.85E-01 | 8.22E-01 | | 0.00 | 0.00 | 0.96 | 0.07 | 1 | 196 |
| GCSE Spanish grade vs objmanip.v | 198 | 0.13 | 0.00 | 0.26 | 0.07 | 1.97 | 5.07E-02 | 7.62E-02 | | 0.02 | 0.01 | 0.95 | 3.87 | 1 | 196 |
| GCSE Spanish grade vs vis.v | 198 | 0.12 | -0.01 | 0.26 | 0.07 | 1.81 | 7.25E-02 | 1.04E-01 | | 0.02 | 0.01 | 0.95 | 3.26 | 1 | 196 |
| GCSE Spanish grade vs nav.v | 198 | 0.12 | -0.02 | 0.26 | 0.07 | 1.73 | 8.49E-02 | 1.19E-01 | | 0.02 | 0.01 | 0.95 | 3.00 | 1 | 196 |
| GCSE Spanish grade vs spatab.v | 198 | 0.12 | -0.01 | 0.26 | 0.07 | 1.81 | 7.25E-02 | 1.04E-01 | | 0.02 | 0.01 | 0.95 | 3.26 | 1 | 196 |
| GCSE German grade vs objmanip | 314 | 0.17 | 0.05 | 0.28 | 0.06 | 2.83 | 4.92E-03 | 9.33E-03 | ** | 0.03 | 0.02 | 0.98 | 8.02 | 1 | 312 |
| GCSE German grade vs vis | 314 | 0.16 | 0.04 | 0.27 | 0.06 | 2.71 | 7.15E-03 | 1.31E-02 | * | 0.02 | 0.02 | 0.98 | 7.33 | 1 | 312 |
| GCSE German grade vs nav | 314 | 0.15 | 0.03 | 0.27 | 0.06 | 2.51 | 1.26E-02 | 2.22E-02 | * | 0.02 | 0.02 | 0.98 | 6.29 | 1 | 312 |
| GCSE German grade vs spatab | 314 | 0.16 | 0.04 | 0.27 | 0.06 | 2.71 | 7.15E-03 | 1.31E-02 | * | 0.02 | 0.02 | 0.98 | 7.33 | 1 | 312 |
| GCSE German grade vs objmanip.g | 244 | -0.12 | -0.25 | 0.01 | 0.07 | -1.84 | 6.66E-02 | 9.70E-02 | | 0.01 | 0.01 | 1.00 | 3.40 | 1 | 242 |
| GCSE German grade vs vis.g | 244 | -0.11 | -0.23 | 0.02 | 0.06 | -1.66 | 9.83E-02 | 1.35E-01 | | 0.01 | 0.01 | 1.00 | 2.75 | 1 | 242 |
| GCSE German grade vs nav.g | 244 | -0.11 | -0.24 | 0.02 | 0.06 | -1.71 | 8.92E-02 | 1.25E-01 | | 0.01 | 0.01 | 1.00 | 2.91 | 1 | 242 |
| GCSE German grade vs spatab.g | 244 | -0.11 | -0.23 | 0.02 | 0.06 | -1.66 | 9.83E-02 | 1.35E-01 | | 0.01 | 0.01 | 1.00 | 2.75 | 1 | 242 |
| GCSE German grade vs objmanip.v | 244 | -0.02 | -0.15 | 0.11 | 0.06 | -0.29 | 7.72E-01 | 8.15E-01 | | 0.00 | 0.00 | 1.01 | 0.08 | 1 | 242 |
| GCSE German grade vs vis.v | 244 | -0.02 | -0.14 | 0.11 | 0.06 | -0.25 | 8.04E-01 | 8.35E-01 | | 0.00 | 0.00 | 1.01 | 0.06 | 1 | 242 |
| GCSE German grade vs nav.v | 244 | -0.03 | -0.15 | 0.10 | 0.06 | -0.44 | 6.59E-01 | 7.13E-01 | | 0.00 | 0.00 | 1.01 | 0.19 | 1 | 242 |
| GCSE German grade vs spatab.v | 244 | -0.02 | -0.14 | 0.11 | 0.06 | -0.25 | 8.04E-01 | 8.35E-01 | | 0.00 | 0.00 | 1.01 | 0.06 | 1 | 242 |
| A(S)-Level English mean grade vs objmanip | 403 | 0.13 | 0.02 | 0.23 | 0.05 | 2.41 | 1.62E-02 | 2.76E-02 | * | 0.01 | 0.01 | 0.98 | 5.83 | 1 | 401 |
| A(S)-Level English mean grade vs vis | 403 | 0.13 | 0.03 | 0.23 | 0.05 | 2.45 | 1.45E-02 | 2.53E-02 | * | 0.01 | 0.01 | 0.98 | 6.02 | 1 | 401 |
| A(S)-Level English mean grade vs nav | 403 | 0.14 | 0.03 | 0.24 | 0.05 | 2.55 | 1.11E-02 | 1.96E-02 | * | 0.02 | 0.01 | 0.98 | 6.51 | 1 | 401 |
| A(S)-Level English mean grade vs spatab | 403 | 0.13 | 0.03 | 0.23 | 0.05 | 2.45 | 1.45E-02 | 2.53E-02 | * | 0.01 | 0.01 | 0.98 | 6.02 | 1 | 401 |
| A(S)-Level English mean grade vs objmanip.g | 313 | -0.14 | -0.25 | -0.02 | 0.06 | -2.33 | 2.04E-02 | 3.37E-02 | * | 0.02 | 0.01 | 1.00 | 5.43 | 1 | 311 |
| A(S)-Level English mean grade vs vis.g | 313 | -0.12 | -0.23 | -0.01 | 0.06 | -2.11 | 3.59E-02 | 5.59E-02 | | 0.01 | 0.01 | 1.01 | 4.44 | 1 | 311 |
| A(S)-Level English mean grade vs nav.g | 313 | -0.11 | -0.22 | 0.00 | 0.06 | -1.94 | 5.37E-02 | 8.02E-02 | | 0.01 | 0.01 | 1.01 | 3.75 | 1 | 311 |
| A(S)-Level English mean grade vs spatab.g | 313 | -0.12 | -0.23 | -0.01 | 0.06 | -2.11 | 3.59E-02 | 5.59E-02 | | 0.01 | 0.01 | 1.01 | 4.44 | 1 | 311 |
| A(S)-Level English mean grade vs objmanip.v | 314 | -0.07 | -0.19 | 0.04 | 0.06 | -1.24 | 2.16E-01 | 2.72E-01 | | 0.00 | 0.00 | 1.01 | 1.53 | 1 | 312 |
| A(S)-Level English mean grade vs vis.v | 314 | -0.06 | -0.18 | 0.05 | 0.06 | -1.12 | 2.66E-01 | 3.28E-01 | | 0.00 | 0.00 | 1.01 | 1.24 | 1 | 312 |
| A(S)-Level English mean grade vs nav.v | 314 | -0.06 | -0.18 | 0.05 | 0.06 | -1.10 | 2.72E-01 | 3.35E-01 | | 0.00 | 0.00 | 1.01 | 1.21 | 1 | 312 |
| A(S)-Level English mean grade vs spatab.v | 314 | -0.06 | -0.18 | 0.05 | 0.06 | -1.12 | 2.66E-01 | 3.28E-01 | | 0.00 | 0.00 | 1.01 | 1.24 | 1 | 312 |
| A(S)-Level Technology mean grade vs objmanip | 205 | 0.31 | 0.17 | 0.45 | 0.07 | 4.33 | 2.30E-05 | 5.72E-05 | *** | 0.08 | 0.08 | 0.96 | 18.78 | 1 | 203 |
| A(S)-Level Technology mean grade vs vis | 205 | 0.32 | 0.17 | 0.46 | 0.07 | 4.21 | 3.88E-05 | 9.28E-05 | *** | 0.08 | 0.08 | 0.96 | 17.70 | 1 | 203 |
| A(S)-Level Technology mean grade vs nav | 205 | 0.30 | 0.15 | 0.46 | 0.08 | 3.86 | 1.55E-04 | 3.52E-04 | *** | 0.07 | 0.06 | 0.96 | 14.87 | 1 | 203 |
| A(S)-Level Technology mean grade vs spatab | 205 | 0.32 | 0.17 | 0.46 | 0.07 | 4.21 | 3.88E-05 | 9.28E-05 | *** | 0.08 | 0.08 | 0.96 | 17.70 | 1 | 203 |
| A(S)-Level Technology mean grade vs objmanip.g | 167 | 0.14 | -0.01 | 0.29 | 0.08 | 1.89 | 6.10E-02 | 8.94E-02 | | 0.02 | 0.02 | 0.97 | 3.56 | 1 | 165 |
| A(S)-Level Technology mean grade vs vis.g | 167 | 0.13 | -0.03 | 0.29 | 0.08 | 1.65 | 1.01E-01 | 1.38E-01 | | 0.02 | 0.01 | 0.97 | 2.72 | 1 | 165 |
| A(S)-Level Technology mean grade vs nav.g | 167 | 0.11 | -0.05 | 0.28 | 0.08 | 1.36 | 1.75E-01 | 2.28E-01 | | 0.01 | 0.01 | 0.97 | 1.86 | 1 | 165 |
| A(S)-Level Technology mean grade vs spatab.g | 167 | 0.13 | -0.03 | 0.29 | 0.08 | 1.65 | 1.01E-01 | 1.38E-01 | | 0.02 | 0.01 | 0.97 | 2.72 | 1 | 165 |
| A(S)-Level Technology mean grade vs objmanip.v | 167 | 0.16 | 0.01 | 0.30 | 0.07 | 2.07 | 3.96E-02 | 6.09E-02 | | 0.03 | 0.02 | 0.97 | 4.30 | 1 | 165 |
| A(S)-Level Technology mean grade vs vis.v | 167 | 0.14 | -0.01 | 0.29 | 0.08 | 1.84 | 6.72E-02 | 9.73E-02 | | 0.02 | 0.01 | 0.97 | 3.40 | 1 | 165 |

| | | | | | | | | | | | | | | | |
|---|---|---|---|---|---|---|---|---|---|---|---|---|---|---|---|
| A(S)-Level Technology mean grade vs nav.v | 167 | 0.13 | -0.03 | 0.28 | 0.08 | 1.56 | 1.20E-01 | 1.59E-01 | | 0.01 | 0.01 | 0.97 | 2.44 | 1 | 165 |
| A(S)-Level Technology mean grade vs spatab.v | 167 | 0.14 | -0.01 | 0.29 | 0.08 | 1.84 | 6.72E-02 | 9.73E-02 | | 0.02 | 0.01 | 0.97 | 3.40 | 1 | 165 |
| A(S)-Level Humanities mean grade vs objmanip | 990 | 0.08 | 0.01 | 0.15 | 0.03 | 2.38 | 1.73E-02 | 2.90E-02 | * | 0.01 | 0.00 | 1.00 | 5.69 | 1 | 988 |
| A(S)-Level Humanities mean grade vs vis | 990 | 0.07 | 0.01 | 0.14 | 0.03 | 2.12 | 3.45E-02 | 5.41E-02 | | 0.00 | 0.00 | 1.00 | 4.48 | 1 | 988 |
| A(S)-Level Humanities mean grade vs nav | 990 | 0.07 | 0.00 | 0.14 | 0.03 | 2.05 | 4.11E-02 | 6.29E-02 | | 0.00 | 0.00 | 1.00 | 4.18 | 1 | 988 |
| A(S)-Level Humanities mean grade vs spatab | 990 | 0.07 | 0.01 | 0.14 | 0.03 | 2.12 | 3.45E-02 | 5.41E-02 | | 0.00 | 0.00 | 1.00 | 4.48 | 1 | 988 |
| A(S)-Level Humanities mean grade vs objmanip.g | 772 | -0.11 | -0.18 | -0.04 | 0.04 | -3.12 | 1.87E-03 | 3.72E-03 | ** | 0.01 | 0.01 | 0.99 | 9.74 | 1 | 770 |
| A(S)-Level Humanities mean grade vs vis.g | 772 | -0.12 | -0.19 | -0.05 | 0.04 | -3.36 | 8.20E-04 | 1.71E-03 | ** | 0.01 | 0.01 | 0.98 | 11.29 | 1 | 770 |
| A(S)-Level Humanities mean grade vs nav.g | 772 | -0.11 | -0.18 | -0.04 | 0.04 | -3.15 | 1.72E-03 | 3.42E-03 | ** | 0.01 | 0.01 | 0.99 | 9.90 | 1 | 770 |
| A(S)-Level Humanities mean grade vs spatab.g | 772 | -0.12 | -0.19 | -0.05 | 0.04 | -3.36 | 8.20E-04 | 1.71E-03 | ** | 0.01 | 0.01 | 0.98 | 11.29 | 1 | 770 |
| A(S)-Level Humanities mean grade vs objmanip.v | 773 | -0.08 | -0.15 | 0.00 | 0.04 | -2.09 | 3.66E-02 | 5.68E-02 | | 0.01 | 0.00 | 0.99 | 4.38 | 1 | 771 |
| A(S)-Level Humanities mean grade vs vis.v | 773 | -0.09 | -0.16 | -0.02 | 0.04 | -2.39 | 1.71E-02 | 2.88E-02 | * | 0.01 | 0.01 | 0.99 | 5.72 | 1 | 771 |
| A(S)-Level Humanities mean grade vs nav.v | 773 | -0.08 | -0.16 | -0.01 | 0.04 | -2.36 | 1.86E-02 | 3.09E-02 | * | 0.01 | 0.01 | 0.99 | 5.57 | 1 | 771 |
| A(S)-Level Humanities mean grade vs spatab.v | 773 | -0.09 | -0.16 | -0.02 | 0.04 | -2.39 | 1.71E-02 | 2.88E-02 | * | 0.01 | 0.01 | 0.99 | 5.72 | 1 | 771 |
| A(S)-Level Languages mean grade vs objmanip | 160 | 0.07 | -0.10 | 0.25 | 0.09 | 0.85 | 3.99E-01 | 4.60E-01 | | 0.00 | 0.00 | 1.02 | 0.72 | 1 | 158 |
| A(S)-Level Languages mean grade vs vis | 160 | 0.06 | -0.11 | 0.24 | 0.09 | 0.72 | 4.73E-01 | 5.39E-01 | | 0.00 | 0.00 | 1.02 | 0.52 | 1 | 158 |
| A(S)-Level Languages mean grade vs nav | 160 | 0.05 | -0.13 | 0.23 | 0.09 | 0.52 | 6.07E-01 | 6.67E-01 | | 0.00 | 0.00 | 1.02 | 0.27 | 1 | 158 |
| A(S)-Level Languages mean grade vs spatab | 160 | 0.06 | -0.11 | 0.24 | 0.09 | 0.72 | 4.73E-01 | 5.39E-01 | | 0.00 | 0.00 | 1.02 | 0.52 | 1 | 158 |
| A(S)-Level Languages mean grade vs objmanip.g | 126 | -0.11 | -0.29 | 0.06 | 0.09 | -1.27 | 2.06E-01 | 2.63E-01 | | 0.01 | 0.00 | 0.94 | 1.62 | 1 | 124 |
| A(S)-Level Languages mean grade vs vis.g | 126 | -0.11 | -0.29 | 0.06 | 0.09 | -1.29 | 2.01E-01 | 2.58E-01 | | 0.01 | 0.01 | 0.94 | 1.65 | 1 | 124 |
| A(S)-Level Languages mean grade vs nav.g | 126 | -0.11 | -0.29 | 0.07 | 0.09 | -1.19 | 2.36E-01 | 2.94E-01 | | 0.01 | 0.00 | 0.94 | 1.42 | 1 | 124 |
| A(S)-Level Languages mean grade vs spatab.g | 126 | -0.11 | -0.29 | 0.06 | 0.09 | -1.29 | 2.01E-01 | 2.58E-01 | | 0.01 | 0.01 | 0.94 | 1.65 | 1 | 124 |
| A(S)-Level Languages mean grade vs objmanip.v | 126 | -0.08 | -0.25 | 0.09 | 0.09 | -0.90 | 3.70E-01 | 4.33E-01 | | 0.01 | 0.00 | 0.94 | 0.81 | 1 | 124 |
| A(S)-Level Languages mean grade vs vis.v | 126 | -0.08 | -0.26 | 0.09 | 0.09 | -0.96 | 3.39E-01 | 4.04E-01 | | 0.01 | 0.00 | 0.94 | 0.92 | 1 | 124 |
| A(S)-Level Languages mean grade vs nav.v | 126 | -0.09 | -0.27 | 0.09 | 0.09 | -0.94 | 3.50E-01 | 4.16E-01 | | 0.01 | 0.00 | 0.94 | 0.88 | 1 | 124 |
| A(S)-Level Languages mean grade vs spatab.v | 126 | -0.08 | -0.26 | 0.09 | 0.09 | -0.96 | 3.39E-01 | 4.04E-01 | | 0.01 | 0.00 | 0.94 | 0.92 | 1 | 124 |
| A(S)-Level Vocational mean grade vs objmanip | 268 | 0.09 | -0.05 | 0.22 | 0.07 | 1.28 | 2.02E-01 | 2.59E-01 | | 0.01 | 0.00 | 1.02 | 1.63 | 1 | 266 |
| A(S)-Level Vocational mean grade vs vis | 268 | 0.13 | -0.01 | 0.26 | 0.07 | 1.88 | 6.07E-02 | 8.92E-02 | | 0.01 | 0.01 | 1.02 | 3.55 | 1 | 266 |
| A(S)-Level Vocational mean grade vs nav | 268 | 0.15 | 0.02 | 0.29 | 0.07 | 2.19 | 2.95E-02 | 4.70E-02 | * | 0.02 | 0.01 | 1.01 | 4.79 | 1 | 266 |
| A(S)-Level Vocational mean grade vs spatab | 268 | 0.13 | -0.01 | 0.26 | 0.07 | 1.88 | 6.07E-02 | 8.92E-02 | | 0.01 | 0.01 | 1.02 | 3.55 | 1 | 266 |
| A(S)-Level Vocational mean grade vs objmanip.g | 204 | -0.06 | -0.20 | 0.08 | 0.07 | -0.89 | 3.74E-01 | 4.34E-01 | | 0.00 | 0.00 | 1.02 | 0.79 | 1 | 202 |
| A(S)-Level Vocational mean grade vs vis.g | 204 | -0.01 | -0.15 | 0.13 | 0.07 | -0.15 | 8.81E-01 | 9.02E-01 | | 0.00 | 0.00 | 1.02 | 0.02 | 1 | 202 |
| A(S)-Level Vocational mean grade vs nav.g | 204 | 0.01 | -0.13 | 0.16 | 0.07 | 0.19 | 8.53E-01 | 8.79E-01 | | 0.00 | 0.00 | 1.02 | 0.03 | 1 | 202 |
| A(S)-Level Vocational mean grade vs spatab.g | 204 | -0.01 | -0.15 | 0.13 | 0.07 | -0.15 | 8.81E-01 | 9.02E-01 | | 0.00 | 0.00 | 1.02 | 0.02 | 1 | 202 |
| A(S)-Level Vocational mean grade vs objmanip.v | 205 | -0.03 | -0.17 | 0.11 | 0.07 | -0.38 | 7.02E-01 | 7.56E-01 | | 0.00 | 0.00 | 1.03 | 0.15 | 1 | 203 |
| A(S)-Level Vocational mean grade vs vis.v | 205 | 0.02 | -0.12 | 0.16 | 0.07 | 0.27 | 7.84E-01 | 8.22E-01 | | 0.00 | 0.00 | 1.03 | 0.08 | 1 | 203 |
| A(S)-Level Vocational mean grade vs nav.v | 205 | 0.04 | -0.10 | 0.18 | 0.07 | 0.53 | 5.97E-01 | 6.61E-01 | | 0.00 | 0.00 | 1.03 | 0.28 | 1 | 203 |
| A(S)-Level Vocational mean grade vs spatab.v | 205 | 0.02 | -0.12 | 0.16 | 0.07 | 0.27 | 7.84E-01 | 8.22E-01 | | 0.00 | 0.00 | 1.03 | 0.08 | 1 | 203 |
| A(S)-Level overall mean grade vs objmanip | 1207 | 0.20 | 0.14 | 0.26 | 0.03 | 6.60 | 6.19E-11 | 2.19E-10 | *** | 0.03 | 0.03 | 0.98 | 43.55 | 1 | 1205 |
| A(S)-Level overall mean grade vs vis | 1207 | 0.19 | 0.13 | 0.25 | 0.03 | 6.09 | 1.47E-09 | 4.72E-09 | *** | 0.03 | 0.03 | 0.99 | 37.14 | 1 | 1205 |
| A(S)-Level overall mean grade vs nav | 1207 | 0.18 | 0.12 | 0.24 | 0.03 | 5.84 | 6.58E-09 | 2.01E-08 | *** | 0.03 | 0.03 | 0.99 | 34.14 | 1 | 1205 |
| A(S)-Level overall mean grade vs spatab | 1207 | 0.19 | 0.13 | 0.25 | 0.03 | 6.09 | 1.47E-09 | 4.72E-09 | *** | 0.03 | 0.03 | 0.99 | 37.14 | 1 | 1205 |

| | | | | | | | | | | | | | | | |
|---|---|---|---|---|---|---|---|---|---|---|---|---|---|---|---|
| A(S)-Level overall mean grade vs objmanip.g | 951 | -0.04 | -0.10 | 0.03 | 0.03 | -1.17 | 2.42E-01 | 3.01E-01 | | 0.00 | 0.00 | 1.00 | 1.37 | 1 | 949 |
| A(S)-Level overall mean grade vs vis.g | 951 | -0.05 | -0.12 | 0.01 | 0.03 | -1.57 | 1.17E-01 | 1.57E-01 | | 0.00 | 0.00 | 1.00 | 2.46 | 1 | 949 |
| A(S)-Level overall mean grade vs nav.g | 951 | -0.05 | -0.12 | 0.01 | 0.03 | -1.64 | 1.02E-01 | 1.39E-01 | | 0.00 | 0.00 | 1.00 | 2.68 | 1 | 949 |
| A(S)-Level overall mean grade vs spatab.g | 951 | -0.05 | -0.12 | 0.01 | 0.03 | -1.57 | 1.17E-01 | 1.57E-01 | | 0.00 | 0.00 | 1.00 | 2.46 | 1 | 949 |
| A(S)-Level overall mean grade vs objmanip.v | 952 | 0.02 | -0.04 | 0.08 | 0.03 | 0.61 | 5.44E-01 | 6.12E-01 | | 0.00 | 0.00 | 1.00 | 0.37 | 1 | 950 |
| A(S)-Level overall mean grade vs vis.v | 952 | 0.00 | -0.06 | 0.07 | 0.03 | 0.15 | 8.84E-01 | 9.02E-01 | | 0.00 | 0.00 | 1.00 | 0.02 | 1 | 950 |
| A(S)-Level overall mean grade vs nav.v | 952 | 0.00 | -0.07 | 0.06 | 0.03 | -0.12 | 9.04E-01 | 9.17E-01 | | 0.00 | 0.00 | 1.00 | 0.01 | 1 | 950 |
| A(S)-Level overall mean grade vs spatab.v | 952 | 0.00 | -0.06 | 0.07 | 0.03 | 0.15 | 8.84E-01 | 9.02E-01 | | 0.00 | 0.00 | 1.00 | 0.02 | 1 | 950 |
| A-Level overall mean grade vs objmanip | 1171 | 0.21 | 0.15 | 0.27 | 0.03 | 6.72 | 2.88E-11 | 1.04E-10 | *** | 0.04 | 0.04 | 0.99 | 45.12 | 1 | 1169 |
| A-Level overall mean grade vs vis | 1171 | 0.19 | 0.13 | 0.25 | 0.03 | 6.14 | 1.13E-09 | 3.65E-09 | *** | 0.03 | 0.03 | 1.00 | 37.70 | 1 | 1169 |
| A-Level overall mean grade vs nav | 1171 | 0.18 | 0.12 | 0.25 | 0.03 | 5.86 | 6.09E-09 | 1.87E-08 | *** | 0.03 | 0.03 | 1.00 | 34.31 | 1 | 1169 |
| A-Level overall mean grade vs spatab | 1171 | 0.19 | 0.13 | 0.25 | 0.03 | 6.14 | 1.13E-09 | 3.65E-09 | *** | 0.03 | 0.03 | 1.00 | 37.70 | 1 | 1169 |
| A-Level overall mean grade vs objmanip.g | 919 | -0.02 | -0.09 | 0.05 | 0.03 | -0.56 | 5.77E-01 | 6.43E-01 | | 0.00 | 0.00 | 1.00 | 0.31 | 1 | 917 |
| A-Level overall mean grade vs vis.g | 919 | -0.04 | -0.10 | 0.03 | 0.03 | -1.05 | 2.92E-01 | 3.55E-01 | | 0.00 | 0.00 | 1.00 | 1.11 | 1 | 917 |
| A-Level overall mean grade vs nav.g | 919 | -0.04 | -0.11 | 0.03 | 0.03 | -1.19 | 2.36E-01 | 2.94E-01 | | 0.00 | 0.00 | 1.00 | 1.41 | 1 | 917 |
| A-Level overall mean grade vs spatab.g | 919 | -0.04 | -0.10 | 0.03 | 0.03 | -1.05 | 2.92E-01 | 3.55E-01 | | 0.00 | 0.00 | 1.00 | 1.11 | 1 | 917 |
| A-Level overall mean grade vs objmanip.v | 920 | 0.04 | -0.03 | 0.10 | 0.03 | 1.07 | 2.85E-01 | 3.49E-01 | | 0.00 | 0.00 | 1.00 | 1.14 | 1 | 918 |
| A-Level overall mean grade vs vis.v | 920 | 0.02 | -0.05 | 0.08 | 0.03 | 0.52 | 6.00E-01 | 6.61E-01 | | 0.00 | 0.00 | 1.00 | 0.28 | 1 | 918 |
| A-Level overall mean grade vs nav.v | 920 | 0.01 | -0.06 | 0.07 | 0.03 | 0.22 | 8.25E-01 | 8.52E-01 | | 0.00 | 0.00 | 1.00 | 0.05 | 1 | 918 |
| A-Level overall mean grade vs spatab.v | 920 | 0.02 | -0.05 | 0.08 | 0.03 | 0.52 | 6.00E-01 | 6.61E-01 | | 0.00 | 0.00 | 1.00 | 0.28 | 1 | 918 |

**Table S9.** Linear regression model estimates and fit indices for individual spatial ability items and two key outcomes (GCSE STEM mean grade and STEM pipeline). p.adjusted denotes Benjamini-Hochberg corrected p-values.

| mod | nobs | estimate | lower 95% | upper 95% | se | t | p.value | p.adjusted | sig | r.squared | adj.r.squared | sd | F | df | df.residual |
|---|---|---|---|---|---|---|---|---|---|---|---|---|---|---|---|
| GCSE STEM mean grade vs cross sections | 1275 | 0.33 | 0.28 | 0.38 | 0.02 | 13.63 | 1.41E-39 | 6.46E-39 | *** | 0.13 | 0.13 | 0.86 | 185.72 | 1 | 1273 |
| GCSE STEM mean grade vs 2D drawing | 1271 | 0.39 | 0.35 | 0.44 | 0.02 | 17.03 | 1.02E-58 | 3.28E-57 | *** | 0.19 | 0.19 | 0.82 | 289.94 | 1 | 1269 |
| GCSE STEM mean grade vs pattern assembly | 1217 | 0.37 | 0.32 | 0.42 | 0.02 | 15.19 | 7.19E-48 | 7.67E-47 | *** | 0.16 | 0.16 | 0.84 | 230.87 | 1 | 1215 |
| GCSE STEM mean grade vs perspective taking | 1149 | 0.19 | 0.14 | 0.24 | 0.03 | 7.56 | 8.23E-14 | 1.32E-13 | *** | 0.05 | 0.05 | 0.87 | 57.15 | 1 | 1147 |
| GCSE STEM mean grade vs mechanical reasoning | 1222 | 0.33 | 0.28 | 0.38 | 0.02 | 13.46 | 1.24E-38 | 4.98E-38 | *** | 0.13 | 0.13 | 0.85 | 181.31 | 1 | 1220 |
| GCSE STEM mean grade vs paper folding | 1181 | 0.36 | 0.31 | 0.41 | 0.02 | 14.82 | 1.03E-45 | 6.60E-45 | *** | 0.16 | 0.16 | 0.83 | 219.73 | 1 | 1179 |
| GCSE STEM mean grade vs 3D drawing | 1125 | 0.36 | 0.31 | 0.41 | 0.02 | 14.87 | 8.79E-46 | 6.60E-45 | *** | 0.16 | 0.16 | 0.81 | 221.08 | 1 | 1123 |
| GCSE STEM mean grade vs shape rotation | 1148 | 0.38 | 0.33 | 0.43 | 0.02 | 15.40 | 9.28E-49 | 1.48E-47 | *** | 0.17 | 0.17 | 0.83 | 237.04 | 1 | 1146 |
| GCSE STEM mean grade vs Elithorn mazes | 1056 | 0.22 | 0.17 | 0.27 | 0.03 | 8.48 | 7.63E-17 | 1.44E-16 | *** | 0.06 | 0.06 | 0.84 | 71.88 | 1 | 1054 |
| GCSE STEM mean grade vs mazes | 1121 | 0.21 | 0.16 | 0.26 | 0.03 | 7.98 | 3.51E-15 | 6.24E-15 | *** | 0.05 | 0.05 | 0.87 | 63.73 | 1 | 1119 |
| GCSE STEM mean grade vs orientation directions | 1100 | 0.36 | 0.32 | 0.41 | 0.02 | 14.80 | 2.38E-45 | 1.27E-44 | *** | 0.17 | 0.17 | 0.81 | 219.17 | 1 | 1098 |
| GCSE STEM mean grade vs orientation landmarks | 1074 | 0.20 | 0.15 | 0.25 | 0.03 | 7.44 | 2.09E-13 | 3.19E-13 | *** | 0.05 | 0.05 | 0.86 | 55.32 | 1 | 1072 |
| GCSE STEM mean grade vs map reading (no memory) | 1044 | 0.21 | 0.15 | 0.26 | 0.03 | 7.73 | 2.62E-14 | 4.40E-14 | *** | 0.05 | 0.05 | 0.85 | 59.68 | 1 | 1042 |
| GCSE STEM mean grade vs map reading (memory) | 1030 | 0.12 | 0.06 | 0.17 | 0.03 | 4.24 | 2.44E-05 | 2.79E-05 | *** | 0.02 | 0.02 | 0.86 | 17.97 | 1 | 1028 |
| GCSE STEM mean grade vs large-scale perspective taking | 1077 | 0.18 | 0.13 | 0.24 | 0.03 | 6.78 | 1.93E-11 | 2.81E-11 | *** | 0.04 | 0.04 | 0.86 | 46.02 | 1 | 1075 |
| GCSE STEM mean grade vs scanning | 1033 | 0.08 | 0.03 | 0.14 | 0.03 | 3.11 | 1.95E-03 | 2.08E-03 | ** | 0.01 | 0.01 | 0.87 | 9.64 | 1 | 1031 |
| STEM pipeline vs cross sections | 1005 | 0.30 | 0.24 | 0.36 | 0.03 | 9.45 | 2.32E-20 | 4.94E-20 | *** | 0.08 | 0.08 | 0.99 | 89.30 | 1 | 1003 |
| STEM pipeline vs 2D drawing | 1011 | 0.37 | 0.31 | 0.43 | 0.03 | 11.86 | 1.74E-30 | 5.57E-30 | *** | 0.12 | 0.12 | 0.97 | 140.76 | 1 | 1009 |
| STEM pipeline vs pattern assembly | 972 | 0.34 | 0.28 | 0.41 | 0.03 | 10.59 | 7.32E-25 | 1.80E-24 | *** | 0.10 | 0.10 | 0.99 | 112.09 | 1 | 970 |
| STEM pipeline vs | 934 | 0.19 | 0.12 | 0.25 | 0.03 | 5.67 | 1.92E-08 | 2.46E-08 | *** | 0.03 | 0.03 | 1.02 | 32.13 | 1 | 932 |

| | | | | | | | | | | | | | | | |
|---|---|---|---|---|---|---|---|---|---|---|---|---|---|---|---|
| perspective taking | | | | | | | | | | | | | | | |
| STEM pipeline vs mechanical reasoning | 981 | 0.35 | 0.29 | 0.41 | 0.03 | 11.01 | 1.22E-26 | 3.55E-26 | *** | 0.11 | 0.11 | 0.98 | 121.11 | 1 | 979 |
| STEM pipeline vs paper folding | 943 | 0.39 | 0.33 | 0.45 | 0.03 | 12.38 | 1.01E-32 | 3.58E-32 | *** | 0.14 | 0.14 | 0.97 | 153.28 | 1 | 941 |
| STEM pipeline vs 3D drawing | 909 | 0.36 | 0.29 | 0.42 | 0.03 | 10.71 | 2.85E-25 | 7.59E-25 | *** | 0.11 | 0.11 | 0.99 | 114.63 | 1 | 907 |
| STEM pipeline vs shape rotation | 921 | 0.34 | 0.28 | 0.41 | 0.03 | 10.38 | 5.87E-24 | 1.34E-23 | *** | 0.10 | 0.10 | 0.99 | 107.81 | 1 | 919 |
| STEM pipeline vs Elithorn mazes | 865 | 0.21 | 0.14 | 0.27 | 0.03 | 6.05 | 2.10E-09 | 2.93E-09 | *** | 0.04 | 0.04 | 1.00 | 36.65 | 1 | 863 |
| STEM pipeline vs mazes | 903 | 0.20 | 0.14 | 0.27 | 0.04 | 5.77 | 1.11E-08 | 1.48E-08 | *** | 0.04 | 0.03 | 1.02 | 33.26 | 1 | 901 |
| STEM pipeline vs orientation directions | 844 | 0.32 | 0.25 | 0.39 | 0.03 | 9.28 | 1.39E-19 | 2.78E-19 | *** | 0.09 | 0.09 | 0.99 | 86.13 | 1 | 842 |
| STEM pipeline vs orientation landmarks | 826 | 0.20 | 0.12 | 0.27 | 0.04 | 5.31 | 1.41E-07 | 1.73E-07 | *** | 0.03 | 0.03 | 1.02 | 28.20 | 1 | 824 |
| STEM pipeline vs map reading (no memory) | 805 | 0.16 | 0.09 | 0.23 | 0.04 | 4.35 | 1.56E-05 | 1.85E-05 | *** | 0.02 | 0.02 | 1.03 | 18.89 | 1 | 803 |
| STEM pipeline vs map reading (memory) | 796 | 0.08 | 0.00 | 0.16 | 0.04 | 2.09 | 3.68E-02 | 3.80E-02 | * | 0.01 | 0.00 | 1.03 | 4.38 | 1 | 794 |
| STEM pipeline vs large-scale perspective taking | 830 | 0.13 | 0.06 | 0.20 | 0.04 | 3.48 | 5.25E-04 | 5.79E-04 | *** | 0.01 | 0.01 | 1.03 | 12.12 | 1 | 828 |
| STEM pipeline vs scanning | 796 | 0.06 | -0.01 | 0.13 | 0.04 | 1.63 | 1.04E-01 | 1.04E-01 | | 0.00 | 0.00 | 1.04 | 2.65 | 1 | 794 |

**Table S10.** Logistic regression model estimates and fit indices. spatab = spatial ability; nav = nav; vis = visualization, objmanip = object manipulation, v = verbal ability. .g and .v suffixes denote *g*- and v-corrected predictors, respectively. p.adjusted denote Benjamini-Hochberg corrected p-values.

| model | nobs | odds.ratio | 95% lower | 95% upper | se | z | p.value | p.adjusted | sig | nagelkerke r^2 | null.deviance | df.null | deviance | df.residual | logLik | AIC |
|---|---|---|---|---|---|---|---|---|---|---|---|---|---|---|---|---|
| Chose a STEM GCSE vs objmanip | 1671 | 2.33 | 1.95 | 2.81 | 0.09 | 9.06 | 1.32E-19 | 1.39E-18 | *** | 0.12 | 962.20 | 1670 | 872.72 | 1669 | -436.36 | 876.72 |
| Chose a STEM GCSE vs vis | 1671 | 2.27 | 1.91 | 2.73 | 0.09 | 8.98 | 2.78E-19 | 2.33E-18 | *** | 0.11 | 962.20 | 1670 | 876.00 | 1669 | -438.00 | 880.00 |
| Chose a STEM GCSE vs nav | 1671 | 2.20 | 1.84 | 2.62 | 0.09 | 8.74 | 2.35E-18 | 1.65E-17 | *** | 0.11 | 962.20 | 1670 | 882.10 | 1669 | -441.05 | 886.10 |
| Chose a STEM GCSE vs spatab | 1671 | 2.27 | 1.91 | 2.73 | 0.09 | 8.98 | 2.78E-19 | 2.33E-18 | *** | 0.11 | 962.20 | 1670 | 876.00 | 1669 | -438.00 | 880.00 |
| Chose a STEM GCSE vs objmanip.g | 1322 | 1.54 | 1.26 | 1.89 | 0.10 | 4.20 | 2.72E-05 | 5.58E-05 | *** | 0.03 | 708.58 | 1321 | 690.94 | 1320 | -345.47 | 694.94 |
| Chose a STEM GCSE vs vis.g | 1322 | 1.53 | 1.25 | 1.86 | 0.10 | 4.18 | 2.95E-05 | 5.77E-05 | *** | 0.03 | 708.58 | 1321 | 691.25 | 1320 | -345.62 | 695.25 |
| Chose a STEM GCSE vs nav.g | 1322 | 1.48 | 1.22 | 1.80 | 0.10 | 3.94 | 8.31E-05 | 1.59E-04 | *** | 0.03 | 708.58 | 1321 | 693.37 | 1320 | -346.68 | 697.37 |
| Chose a STEM GCSE vs spatab.g | 1322 | 1.53 | 1.25 | 1.86 | 0.10 | 4.18 | 2.95E-05 | 5.77E-05 | *** | 0.03 | 708.58 | 1321 | 691.25 | 1320 | -345.62 | 695.25 |
| Chose a STEM GCSE vs objmanip.v | 1324 | 1.73 | 1.41 | 2.12 | 0.10 | 5.23 | 1.67E-07 | 4.26E-07 | *** | 0.05 | 708.90 | 1323 | 680.91 | 1322 | -340.46 | 684.91 |
| Chose a STEM GCSE vs vis.v | 1324 | 1.69 | 1.39 | 2.07 | 0.10 | 5.16 | 2.50E-07 | 5.99E-07 | *** | 0.05 | 708.90 | 1323 | 682.09 | 1322 | -341.05 | 686.09 |
| Chose a STEM GCSE vs nav.v | 1324 | 1.62 | 1.34 | 1.98 | 0.10 | 4.86 | 1.20E-06 | 2.59E-06 | *** | 0.04 | 708.90 | 1323 | 685.51 | 1322 | -342.76 | 689.51 |
| Chose a STEM GCSE vs spatab.v | 1324 | 1.69 | 1.39 | 2.07 | 0.10 | 5.16 | 2.50E-07 | 5.99E-07 | *** | 0.05 | 708.90 | 1323 | 682.09 | 1322 | -341.05 | 686.09 |
| Chose a STEM A(S)-Level vs objmanip | 1710 | 2.04 | 1.83 | 2.28 | 0.06 | 12.79 | 1.90E-37 | 1.60E-35 | *** | 0.14 | 2367.70 | 1709 | 2178.33 | 1708 | -1089.17 | 2182.33 |
| Chose a STEM A(S)-Level vs vis | 1710 | 1.95 | 1.75 | 2.17 | 0.06 | 12.06 | 1.75E-33 | 4.91E-32 | *** | 0.12 | 2367.70 | 1709 | 2201.47 | 1708 | -1100.74 | 2205.47 |
| Chose a STEM A(S)-Level vs nav | 1710 | 1.89 | 1.70 | 2.11 | 0.06 | 11.55 | 7.51E-31 | 1.58E-29 | *** | 0.11 | 2367.70 | 1709 | 2216.19 | 1708 | -1108.09 | 2220.19 |
| Chose a STEM A(S)-Level vs spatab | 1710 | 1.95 | 1.75 | 2.17 | 0.06 | 12.06 | 1.75E-33 | 4.91E-32 | *** | 0.12 | 2367.70 | 1709 | 2201.47 | 1708 | -1100.74 | 2205.47 |
| Chose a STEM A(S)-Level vs objmanip.g | 1296 | 1.45 | 1.30 | 1.63 | 0.06 | 6.42 | 1.39E-10 | 4.16E-10 | *** | 0.04 | 1796.44 | 1295 | 1753.29 | 1294 | -876.64 | 1757.29 |
| Chose a STEM A(S)-Level vs vis.g | 1296 | 1.38 | 1.23 | 1.54 | 0.06 | 5.54 | 3.05E-08 | 8.01E-08 | *** | 0.03 | 1796.44 | 1295 | 1764.68 | 1294 | -882.34 | 1768.68 |
| Chose a STEM A(S)-Level vs nav.g | 1296 | 1.34 | 1.19 | 1.50 | 0.06 | 5.03 | 4.79E-07 | 1.12E-06 | *** | 0.03 | 1796.44 | 1295 | 1770.30 | 1294 | -885.15 | 1774.30 |
| Chose a STEM A(S)-Level vs spatab.g | 1296 | 1.38 | 1.23 | 1.54 | 0.06 | 5.54 | 3.05E-08 | 8.01E-08 | *** | 0.03 | 1796.44 | 1295 | 1764.68 | 1294 | -882.34 | 1768.68 |
| Chose a STEM A(S)-Level vs objmanip.v | 1297 | 1.64 | 1.46 | 1.84 | 0.06 | 8.28 | 1.23E-16 | 7.98E-16 | *** | 0.07 | 1797.85 | 1296 | 1724.02 | 1295 | -862.01 | 1728.02 |
| Chose a STEM A(S)-Level vs vis.v | 1297 | 1.55 | 1.38 | 1.74 | 0.06 | 7.44 | 1.03E-13 | 4.57E-13 | *** | 0.06 | 1797.85 | 1296 | 1739.05 | 1295 | -869.53 | 1743.05 |
| Chose a STEM A(S)-Level vs nav.v | 1297 | 1.49 | 1.33 | 1.68 | 0.06 | 6.86 | 7.13E-12 | 2.30E-11 | *** | 0.05 | 1797.85 | 1296 | 1748.20 | 1295 | -874.10 | 1752.20 |
| Chose a STEM A(S)-Level vs spatab.v | 1297 | 1.55 | 1.38 | 1.74 | 0.06 | 7.44 | 1.03E-13 | 4.57E-13 | *** | 0.06 | 1797.85 | 1296 | 1739.05 | 1295 | -869.53 | 1743.05 |
| Chose a STEM degree vs objmanip | 1659 | 2.12 | 1.84 | 2.46 | 0.07 | 10.27 | 9.59E-25 | 1.61E-23 | *** | 0.11 | 1682.82 | 1658 | 1559.83 | 1657 | -779.91 | 1563.83 |
| Chose a STEM degree vs vis | 1659 | 2.00 | 1.74 | 2.32 | 0.07 | 9.46 | 2.94E-21 | 3.53E-20 | *** | 0.09 | 1682.82 | 1658 | 1579.21 | 1657 | -789.61 | 1583.21 |
| Chose a STEM degree vs nav | 1659 | 1.88 | 1.64 | 2.17 | 0.07 | 8.78 | 1.68E-18 | 1.28E-17 | *** | 0.08 | 1682.82 | 1658 | 1595.09 | 1657 | -797.54 | 1599.09 |
| Chose a STEM degree vs spatab | 1659 | 2.00 | 1.74 | 2.32 | 0.07 | 9.46 | 2.94E-21 | 3.53E-20 | *** | 0.09 | 1682.82 | 1658 | 1579.21 | 1657 | -789.61 | 1583.21 |
| Chose a STEM degree vs objmanip.g | 1246 | 1.57 | 1.36 | 1.83 | 0.08 | 6.02 | 1.71E-09 | 4.78E-09 | *** | 0.05 | 1307.67 | 1245 | 1268.93 | 1244 | -634.46 | 1272.93 |
| Chose a STEM degree vs vis.g | 1246 | 1.45 | 1.26 | 1.68 | 0.07 | 5.02 | 5.12E-07 | 1.13E-06 | *** | 0.03 | 1307.67 | 1245 | 1281.04 | 1244 | -640.52 | 1285.04 |
| Chose a STEM degree vs nav.g | 1246 | 1.37 | 1.19 | 1.58 | 0.07 | 4.28 | 1.90E-05 | 4.00E-05 | *** | 0.02 | 1307.67 | 1245 | 1288.53 | 1244 | -644.27 | 1292.53 |
| Chose a STEM degree vs spatab.g | 1246 | 1.45 | 1.26 | 1.68 | 0.07 | 5.02 | 5.12E-07 | 1.13E-06 | *** | 0.03 | 1307.67 | 1245 | 1281.04 | 1244 | -640.52 | 1285.04 |
| Chose a STEM degree vs objmanip.v | 1248 | 1.85 | 1.59 | 2.17 | 0.08 | 7.88 | 3.15E-15 | 1.89E-14 | *** | 0.08 | 1311.20 | 1247 | 1241.95 | 1246 | -620.98 | 1245.95 |
| Chose a STEM degree vs vis.v | 1248 | 1.72 | 1.48 | 2.01 | 0.08 | 7.00 | 2.57E-12 | 9.80E-12 | *** | 0.07 | 1311.20 | 1247 | 1257.16 | 1246 | -628.58 | 1261.16 |
| Chose a STEM degree vs nav.v | 1248 | 1.61 | 1.39 | 1.87 | 0.08 | 6.25 | 4.18E-10 | 1.21E-09 | *** | 0.05 | 1311.20 | 1247 | 1268.71 | 1246 | -634.36 | 1272.71 |
| Chose a STEM degree vs spatab.v | 1248 | 1.72 | 1.48 | 2.01 | 0.08 | 7.00 | 2.57E-12 | 9.80E-12 | *** | 0.07 | 1311.20 | 1247 | 1257.16 | 1246 | -628.58 | 1261.16 |

| | | | | | | | | | | | | | | | |
|---|---|---|---|---|---|---|---|---|---|---|---|---|---|---|---|
| Chose a humanities GCSE vs objmanip | 1671 | 1.61 | 1.43 | 1.82 | 0.06 | 7.74 | 9.63E-15 | 4.76E-14 | *** | 0.06 | 1712.57 | 1670 | 1650.41 | 1669 | -825.20 | 1654.41 |
| Chose a humanities GCSE vs vis | 1671 | 1.63 | 1.44 | 1.84 | 0.06 | 7.87 | 3.59E-15 | 1.89E-14 | *** | 0.06 | 1712.57 | 1670 | 1648.70 | 1669 | -824.35 | 1652.70 |
| Chose a humanities GCSE vs nav | 1671 | 1.58 | 1.40 | 1.79 | 0.06 | 7.42 | 1.13E-13 | 4.75E-13 | *** | 0.05 | 1712.57 | 1670 | 1656.12 | 1669 | -828.06 | 1660.12 |
| Chose a humanities GCSE vs spatab | 1671 | 1.63 | 1.44 | 1.84 | 0.06 | 7.87 | 3.59E-15 | 1.89E-14 | *** | 0.06 | 1712.57 | 1670 | 1648.70 | 1669 | -824.35 | 1652.70 |
| Chose a humanities GCSE vs objmanip.g | 1322 | 1.00 | 0.88 | 1.14 | 0.07 | 0.01 | 9.89E-01 | 9.89E-01 | | 0.00 | 1365.18 | 1321 | 1365.18 | 1320 | -682.59 | 1369.18 |
| Chose a humanities GCSE vs vis.g | 1322 | 1.03 | 0.90 | 1.17 | 0.07 | 0.43 | 6.71E-01 | 7.13E-01 | | 0.00 | 1365.18 | 1321 | 1365.00 | 1320 | -682.50 | 1369.00 |
| Chose a humanities GCSE vs nav.g | 1322 | 1.01 | 0.88 | 1.15 | 0.07 | 0.11 | 9.14E-01 | 9.37E-01 | | 0.00 | 1365.18 | 1321 | 1365.17 | 1320 | -682.59 | 1369.17 |
| Chose a humanities GCSE vs spatab.g | 1322 | 1.03 | 0.90 | 1.17 | 0.07 | 0.43 | 6.71E-01 | 7.13E-01 | | 0.00 | 1365.18 | 1321 | 1365.00 | 1320 | -682.50 | 1369.00 |
| Chose a humanities GCSE vs objmanip.v | 1324 | 1.13 | 0.99 | 1.29 | 0.07 | 1.84 | 6.60E-02 | 8.28E-02 | | 0.00 | 1366.13 | 1323 | 1362.76 | 1322 | -681.38 | 1366.76 |
| Chose a humanities GCSE vs vis.v | 1324 | 1.15 | 1.01 | 1.31 | 0.07 | 2.09 | 3.70E-02 | 4.70E-02 | * | 0.01 | 1366.13 | 1323 | 1361.80 | 1322 | -680.90 | 1365.80 |
| Chose a humanities GCSE vs nav.v | 1324 | 1.11 | 0.98 | 1.27 | 0.07 | 1.61 | 1.08E-01 | 1.33E-01 | | 0.00 | 1366.13 | 1323 | 1363.56 | 1322 | -681.78 | 1367.56 |
| Chose a humanities GCSE vs spatab.v | 1324 | 1.15 | 1.01 | 1.31 | 0.07 | 2.09 | 3.70E-02 | 4.70E-02 | * | 0.01 | 1366.13 | 1323 | 1361.80 | 1322 | -680.90 | 1365.80 |
| Chose a humanities A(S)-Level vs objmanip | 1710 | 1.42 | 1.28 | 1.56 | 0.05 | 6.86 | 7.13E-12 | 2.30E-11 | *** | 0.04 | 2277.07 | 1709 | 2228.62 | 1708 | -1114.31 | 2232.62 |
| Chose a humanities A(S)-Level vs vis | 1710 | 1.42 | 1.28 | 1.57 | 0.05 | 6.88 | 5.79E-12 | 2.03E-11 | *** | 0.04 | 2277.07 | 1709 | 2228.24 | 1708 | -1114.12 | 2232.24 |
| Chose a humanities A(S)-Level vs nav | 1710 | 1.41 | 1.28 | 1.56 | 0.05 | 6.78 | 1.21E-11 | 3.78E-11 | *** | 0.04 | 2277.07 | 1709 | 2229.74 | 1708 | -1114.87 | 2233.74 |
| Chose a humanities A(S)-Level vs spatab | 1710 | 1.42 | 1.28 | 1.57 | 0.05 | 6.88 | 5.79E-12 | 2.03E-11 | *** | 0.04 | 2277.07 | 1709 | 2228.24 | 1708 | -1114.12 | 2232.24 |
| Chose a humanities A(S)-Level vs objmanip.g | 1296 | 0.97 | 0.86 | 1.08 | 0.06 | -0.58 | 5.60E-01 | 6.18E-01 | | 0.00 | 1700.93 | 1295 | 1700.59 | 1294 | -850.30 | 1704.59 |
| Chose a humanities A(S)-Level vs vis.g | 1296 | 0.98 | 0.88 | 1.10 | 0.06 | -0.34 | 7.32E-01 | 7.59E-01 | | 0.00 | 1700.93 | 1295 | 1700.81 | 1294 | -850.41 | 1704.81 |
| Chose a humanities A(S)-Level vs nav.g | 1296 | 1.00 | 0.89 | 1.12 | 0.06 | -0.07 | 9.43E-01 | 9.54E-01 | | 0.00 | 1700.93 | 1295 | 1700.93 | 1294 | -850.46 | 1704.93 |
| Chose a humanities A(S)-Level vs spatab.g | 1296 | 0.98 | 0.88 | 1.10 | 0.06 | -0.34 | 7.32E-01 | 7.59E-01 | | 0.00 | 1700.93 | 1295 | 1700.81 | 1294 | -850.41 | 1704.81 |
| Chose a humanities A(S)-Level vs objmanip.v | 1297 | 1.06 | 0.95 | 1.19 | 0.06 | 1.09 | 2.78E-01 | 3.24E-01 | | 0.00 | 1701.84 | 1296 | 1700.66 | 1295 | -850.33 | 1704.66 |
| Chose a humanities A(S)-Level vs vis.v | 1297 | 1.07 | 0.96 | 1.20 | 0.06 | 1.19 | 2.34E-01 | 2.76E-01 | | 0.00 | 1701.84 | 1296 | 1700.42 | 1295 | -850.21 | 1704.42 |
| Chose a humanities A(S)-Level vs nav.v | 1297 | 1.07 | 0.96 | 1.20 | 0.06 | 1.26 | 2.09E-01 | 2.55E-01 | | 0.00 | 1701.84 | 1296 | 1700.27 | 1295 | -850.13 | 1704.27 |
| Chose a humanities A(S)-Level vs spatab.v | 1297 | 1.07 | 0.96 | 1.20 | 0.06 | 1.19 | 2.34E-01 | 2.76E-01 | | 0.00 | 1701.84 | 1296 | 1700.42 | 1295 | -850.21 | 1704.42 |
| Chose a humanities degree vs objmanip | 1659 | 1.03 | 0.91 | 1.16 | 0.06 | 0.44 | 6.62E-01 | 7.13E-01 | | 0.00 | 1709.57 | 1658 | 1709.37 | 1657 | -854.69 | 1713.37 |
| Chose a humanities degree vs vis | 1659 | 1.05 | 0.93 | 1.18 | 0.06 | 0.73 | 4.64E-01 | 5.19E-01 | | 0.00 | 1709.57 | 1658 | 1709.03 | 1657 | -854.51 | 1713.03 |
| Chose a humanities degree vs nav | 1659 | 1.05 | 0.93 | 1.18 | 0.06 | 0.79 | 4.32E-01 | 4.97E-01 | | 0.00 | 1709.57 | 1658 | 1708.94 | 1657 | -854.47 | 1712.94 |
| Chose a humanities degree vs spatab | 1659 | 1.05 | 0.93 | 1.18 | 0.06 | 0.73 | 4.64E-01 | 5.19E-01 | | 0.00 | 1709.57 | 1658 | 1709.03 | 1657 | -854.51 | 1713.03 |
| Chose a humanities degree vs objmanip.g | 1246 | 0.77 | 0.68 | 0.89 | 0.07 | -3.68 | 2.34E-04 | 4.09E-04 | *** | 0.02 | 1322.81 | 1245 | 1309.18 | 1244 | -654.59 | 1313.18 |
| Chose a humanities degree vs vis.g | 1246 | 0.80 | 0.70 | 0.92 | 0.07 | -3.17 | 1.50E-03 | 2.47E-03 | ** | 0.01 | 1322.81 | 1245 | 1312.73 | 1244 | -656.36 | 1316.73 |
| Chose a humanities degree vs nav.g | 1246 | 0.82 | 0.72 | 0.94 | 0.07 | -2.90 | 3.79E-03 | 5.79E-03 | ** | 0.01 | 1322.81 | 1245 | 1314.45 | 1244 | -657.22 | 1318.45 |
| Chose a humanities degree vs spatab.g | 1246 | 0.80 | 0.70 | 0.92 | 0.07 | -3.17 | 1.50E-03 | 2.47E-03 | ** | 0.01 | 1322.81 | 1245 | 1312.73 | 1244 | -656.36 | 1316.73 |
| Chose a humanities degree vs objmanip.v | 1248 | 0.81 | 0.71 | 0.93 | 0.07 | -3.01 | 2.57E-03 | 4.16E-03 | ** | 0.01 | 1326.31 | 1247 | 1317.20 | 1246 | -658.60 | 1321.20 |
| Chose a humanities degree vs vis.v | 1248 | 0.84 | 0.73 | 0.95 | 0.07 | -2.64 | 8.25E-03 | 1.14E-02 | * | 0.01 | 1326.31 | 1247 | 1319.35 | 1246 | -659.67 | 1323.35 |
| Chose a humanities degree vs nav.v | 1248 | 0.84 | 0.74 | 0.96 | 0.07 | -2.52 | 1.19E-02 | 1.56E-02 | * | 0.01 | 1326.31 | 1247 | 1320.01 | 1246 | -660.00 | 1324.01 |
| Chose a humanities degree vs spatab.v | 1248 | 0.84 | 0.73 | 0.95 | 0.07 | -2.64 | 8.25E-03 | 1.14E-02 | * | 0.01 | 1326.31 | 1247 | 1319.35 | 1246 | -659.67 | 1323.35 |

**Table S11**. Univariate twin model ACE estimates and fit indices. A = heritability, C = shared environment, E = non-shared environment. Figures in brackets indicate 95% confidence intervals. Comparative fit statistics (i.e., diffLL, diffdf, and p) indicate model fit comparisons with ACE model.

| Model | A | C | E | ep | -2LL | df | AIC | diffLL | diffdf | p |
|---|---|---|---|---|---|---|---|---|---|---|
| Spatial ability (AE) | 0.71 (0.67, 0.74) | | 0.29 (0.26, 0.33) | 3 | 10631.16 | 3935 | 10637.16 | 0.6403745 | 1 | 0.4235752 |
| Object manipulation (AE) | 0.72 (0.69, 0.75) | | 0.28 (0.25, 0.31) | 3 | 10583.37 | 3935 | 10589.37 | 0.5729981 | 1 | 0.44907 |
| Nav (AE) | 0.69 (0.66, 0.73) | | 0.31 (0.27, 0.34) | 3 | 10665.13 | 3935 | 10671.13 | 0.887608 | 1 | 0.3461263 |
| Visualization (AE) | 0.71 (0.67, 0.74) | | 0.29 (0.26, 0.33) | 3 | 10631.16 | 3935 | 10637.16 | 0.6403745 | 1 | 0.4235752 |
| g (ACE) | 0.67 (0.56, 0.80) | 0.16 (0.05, 0.27) | 0.17 (0.15, 0.19) | 4 | 7344.918 | 2833 | 7352.918 | | | |
| Verbal ability (ACE) | 0.67 (0.55, 0.80) | 0.14 (0.02, 0.25) | 0.19 (0.16, 0.22) | 4 | 7430.288 | 2844 | 7438.388 | | | |
| GCSE STEM + humanities mean grade (ACE) | 0.62 (0.58, 0.66) | 0.27 (0.23, 0.31) | 0.12 (0.11, 0.12) | 4 | 31759.13 | 12982 | 31767.13 | | | |
| GCSE STEM mean grade (ACE) | 0.63 (0.59, 0.67) | 0.23 (0.19, 0.27) | 0.14 (0.13, 0.15) | 4 | 32268.14 | 12947 | 32276.14 | | | |
| GCSE humanities mean grade (ACE) | 0.60 (0.56, 0.65) | 0.25 (0.20, 0.29) | 0.15 (0.14, 0.16) | 4 | 32393.44 | 12946 | 32401.44 | | | |
| GCSE overall mean grade (ACE) | 0.62 (0.58, 0.66) | 0.27 (0.23, 0.31) | 0.11 (0.11, 0.12) | 4 | 31693.96 | 12982 | 31701.96 | | | |
| GCSE Core mean grade (ACE) | 0.63 (0.60, 0.68) | 0.25 (0.21, 0.29) | 0.12 (0.11, 0.12) | 4 | 31803.25 | 12965 | 31811.25 | | | |
| GCSE Maths mean grade (ACE) | 0.65 (0.61, 0.70) | 0.18 (0.13, 0.22) | 0.17 (0.16, 0.18) | 4 | 32621 | 12818 | 32629 | | | |
| GCSE English mean grade (ACE) | 0.60 (0.56, 0.65) | 0.23 (0.18, 0.27) | 0.17 (0.16, 0.18) | 4 | 32705.55 | 12897 | 32713.55 | | | |
| GCSE Science mean grade (ACE) | 0.62 (0.58, 0.67) | 0.21 (0.17, 0.26) | 0.16 (0.15, 0.18) | 4 | 30449.35 | 11991 | 30457.35 | | | |
| GCSE Technology mean grade (ACE) | 0.66 (0.57, 0.74) | 0.1 (0.02, 0.18) | 0.24 (0.22, 0.27) | 4 | 18937.88 | 7076 | 18945.88 | | | |
| GCSE Humanities mean grade (ACE) | 0.56 (0.51, 0.61) | 0.22 (0.17, 0.27) | 0.22 (0.21, 0.24) | 4 | 31967.63 | 12309 | 31975.63 | | | |
| GCSE Languages mean grade (ACE) | 0.53 (0.46, 0.60) | 0.25 (0.19, 0.31) | 0.22 (0.20, 0.24) | 4 | 20183.37 | 7707 | 20191.37 | | | |
| GCSE Maths grade (ACE) | 0.65 (0.55, 0.75) | 0.14 (0.04, 0.23) | 0.22 (0.19, 0.24) | 4 | 13916.11 | 5215 | 13924.11 | | | |
| GCSE Science Core grade (ACE) | 0.63 (0.55, 0.72) | 0.14 (0.06, 0.21) | 0.23 (0.21, 0.25) | 4 | 17839.17 | 6735 | 17847.17 | | | |
| GCSE Statistics grade (ACE) | 0.55 (0.39, 0.72) | 0.27 (0.09, 0.42) | 0.18 (0.15, 0.22) | 4 | 3765.973 | 1440 | 3773.973 | | | |
| GCSE ICT grade (ACE) | 0.59 (0.45, 0.74) | 0.16 (0.02, 0.28) | 0.25 (0.22, 0.30) | 4 | 7290.607 | 2708 | 7298.607 | | | |
| GCSE Science Additional grade (ACE) | 0.64 (0.54, 0.74) | 0.13 (0.07, 0.22) | 0.23 (0.20, 0.25) | 4 | 14314.69 | 5400 | 14322.69 | | | |
| GCSE Physics grade (ACE) | 0.50 (0.41, 0.59) | 0.27 (0.18, 0.35) | 0.23 (0.21, 0.26) | 4 | 13694.61 | 5167 | 13702.61 | | | |
| GCSE Chemistry grade (ACE) | 0.55 (0.46, 0.64) | 0.25 (0.22, 0.33) | 0.20 (0.18, 0.22) | 4 | 13640.04 | 5181 | 13648.04 | | | |
| GCSE Biology grade (ACE) | 0.41 (0.33, 0.49) | 0.37 (0.30, 0.45) | 0.22 (0.19, 0.24) | 4 | 13701.48 | 5209 | 13709.48 | | | |
| GCSE English Language grade (ACE) | 0.57 (0.52, 0.62) | 0.22 (0.18, 0.27) | 0.20 (0.19, 0.22) | 4 | 32940.01 | 12803 | 32948.01 | | | |
| GCSE English Literature grade (ACE) | 0.55 (0.49, 0.61) | 0.20 (0.15, 0.25) | 0.25 (0.23, 0.27) | 4 | 29939.66 | 11383 | 29947.66 | | | |
| GCSE French grade (ACE) | 0.52 (0.43, 0.61) | 0.29 (0.20, 0.37) | 0.19 (0.17, 0.22) | 4 | 11665.75 | 4427 | 11673.75 | | | |
| GCSE History grade (ACE) | 0.52 (0.43, 0.61) | 0.27 (0.19, 0.35) | 0.22 (0.19, 0.24) | 4 | 15134.7 | 5677 | 15142.7 | | | |
| GCSE Spanish grade (ACE) | 0.55 (0.38, 0.73) | 0.27 (0.10, 0.42) | 0.18 (0.15, 0.23) | 4 | 4835.102 | 1796 | 4843.102 | | | |
| GCSE German grade (ACE) | 0.36 (0.22, 0.50) | 0.47 (0.33, 0.58) | 0.18 (0.15, 0.22) | 4 | 4949.238 | 1874 | 4957.238 | | | |
| Number of non-compulsory STEM GCSEs taken (ACE) | 0.58 (0.52, 0.63) | 0.16 (0.13, 0.21) | 0.26 (0.24, 0.28) | 4 | 34266.06 | 13044 | 34274.06 | | | |
| Total number of STEM GCSEs taken (ACE) | 0.59 (0.54, 0.64) | 0.18 (0.13, 0.23) | 0.23 (0.22, 0.25) | 4 | 33973.24 | 13044 | 33981.24 | | | |
| Number of non-compulsory humanities GCSEs taken (ACE) | 0.52 (0.46, 0.57) | 0.18 (0.13, 0.23) | 0.30 (0.28, 0.32) | 4 | 34589.23 | 13044 | 34597.23 | | | |
| Total number of humanities GCSEs taken (ACE) | 0.53 (0.48, 0.58) | 0.21 (0.16, 0.25) | 0.26 (0.25, 0.28) | 4 | 34244.66 | 13044 | 34252.66 | | | |
| A(S)-Level English mean grade (AE) | 0.71 (0.65, 0.76) | | 0.29 (0.24, 0.35) | 3 | 6958.571 | 2517 | 6964.571 | 0.08974957 | 1 | 0.7644958 |
| A(S)-Level Maths mean grade (AE) | 0.67 (0.62, 0.72) | | 0.33 (0.28, 0.38) | 3 | 7795.677 | 2824 | 7801.677 | -1.55E-11 | 1 | 1 |

| | | | | | | | | | | |
|---|---|---|---|---|---|---|---|---|---|---|
| A(S)-Level Science mean grade (AE) | 0.69 (0.65, 0.73) | | 0.31 (0.27, 0.35) | 3 | 9910.43 | 3609 | 9916.43 | -1.52E-09 | 1 | 1 |
| A(S)-Level Technology mean grade (AE) | 0.69 (0.57, 0.74) | | 0.33 (0.26, 0.41) | 3 | 3923.945 | 1414 | 3929.945 | 2.541468 | 1 | 0.1108917 |
| A(S)-Level Humanities mean grade (AE) | 0.61 (0.58, 0.65) | | 0.39 (0.35, 0.42) | 3 | 17476.05 | 6345 | 17482.05 | 1.408072 | 1 | 0.2353768 |
| A(S)-Level Languages mean grade (ACE) | 0.35 (0.07, 0.65) | 0.38 (0.07, 0.61) | 0.27 (0.21, 0.37) | 3 | 2587.563 | 942 | 2595.563 | | | |
| A(S)-Level Vocational mean grade (CE) | | 0.52 (0.45, 0.59) | 0.48 (0.41, 0.55) | 3 | 5501.656 | 1977 | 5507.656 | 1.720206 | 1 | 0.1896666 |
| A(S)-Level overall mean grade (ACE) | 0.64 (0.56, 0.72) | 0.09 (0.02, 0.16) | 0.27 (0.25, 0.29) | 4 | 21252.29 | 7933 | 21260.29 | | | |
| A-Level overall mean grade (ACE) | 0.59 (0.50, 0.67) | 0.13 (0.06, 0.20) | 0.29 (0.27, 0.31) | 4 | 20500.43 | 7624 | 20508.43 | | | |
| A(S)-Level STEM mean grade (AE) | 0.65 (0.61, 0.69) | | 0.35 (0.31, 0.39) | 3 | 14194.16 | 5161 | 14200.16 | -1.91E-08 | 1 | 1 |
| A(S)-Level humanities mean grade (AE) | 0.61 (0.58, 0.65) | | 0.39 (0.35, 0.42) | 3 | 18619.92 | 6776 | 18625.92 | 3.379699 | 1 | 0.06600412 |
| A(S)-Level STEM + humanities mean grade (AE) | 0.69 (0.66, 0.71) | | 0.31 (0.29, 0.34) | 3 | 21193.74 | 7826 | 21199.74 | 2.299048 | 1 | 0.1294533 |
| Number of STEM A(S)-Levels taken (AE) | 0.70 (0.68, 0.72) | | 0.30 (0.28, 0.22) | 3 | 36628.22 | 13683 | 36634.22 | 1.819078 | 1 | 0.1774231 |
| Number of humanities A(S)-Levels taken (AE) | 0.63 (0.61, 0.65) | | 0.37 (0.35, 0.39) | 3 | 37160.28 | 13683 | 37166.28 | 1.840659 | 1 | 0.1748738 |

**Table S12**. Bivariate twin model ACE path coefficients, correlations, and fit indices. a = heritability, c = shared environment, e = non-shared environment. a/c/e11 = estimate of variance unique to variable 1, a/c/e21 = estimate of variance shared between variable 1 and 2, a/c/e22 = estimate of variance unique to variable 2. rPh = phenotypic correlation, rA = genetic correlation, rC = shared environment correlation, rE = non-shared environment correlation. Number suffixes indicate the variable the estimate or coefficient corresponds to, as denoted by the model column.

| Model | | Standardized squared path coefficients | | | | | | | | | Correlations | | | | Fit indices | | |
|---|---|---|---|---|---|---|---|---|---|---|---|---|---|---|---|---|---|
| 1 | 2 | a11 | a21 | a22 | c11 | c21 | c22 | e11 | e21 | e22 | rPh | rA | rC | rE | ep | -2LL | df |
| Spatial ability | GCSE STEM mean grade | 0.65 | 0.26 | 0.36 | 0.04 | 0.03 | 0.2 | 0.28 | 0.01 | 0.13 | 0.51 | 0.65 | 0.37 | 0.28 | 11 | 42080.43 | 16878 |
| Spatial ability | GCSE Maths mean grade | 0.64 | 0.29 | 0.35 | 0.05 | 0 | 0.17 | 0.29 | 0.01 | 0.16 | 0.52 | 0.68 | 0.21 | 0.25 | 11 | 42418.67 | 16749 |
| Spatial ability | GCSE Science mean grade | 0.66 | 0.25 | 0.36 | 0.03 | 0.01 | 0.21 | 0.29 | 0.01 | 0.15 | 0.48 | 0.64 | 0.18 | 0.22 | 11 | 40416.97 | 15922 |
| Spatial ability | GCSE Physics grade | 0.66 | 0.12 | 0.37 | 0.04 | 0.17 | 0.1 | 0.3 | 0.01 | 0.22 | 0.42 | 0.5 | 0.8 | 0.2 | 11 | 24126.85 | 9098 |
| Spatial ability | GCSE Chemistry grade | 0.67 | 0.1 | 0.44 | 0.04 | 0.1 | 0.15 | 0.3 | 0.01 | 0.19 | 0.37 | 0.43 | 0.63 | 0.22 | 11 | 24111.1 | 9112 |
| Spatial ability | GCSE Biology grade | 0.66 | 0.06 | 0.34 | 0.04 | 0.37 | 0 | 0.3 | 0.01 | 0.2 | 0.39 | 0.39 | 1 | 0.25 | 11 | 24153.42 | 9140 |
| Spatial ability | A(S)-Level STEM mean grade | 0.65 | 0.04 | 0.59 | 0.06 | 0.02 | 0 | 0.3 | 0.01 | 0.33 | 0.25 | 0.25 | 1 | 0.2 | 11 | 24734.46 | 9091 |
| Spatial ability | A(S)-Level Maths mean grade | 0.64 | 0.03 | 0.61 | 0.06 | 0.03 | 0 | 0.29 | 0.01 | 0.31 | 0.23 | 0.2 | 1 | 0.16 | 11 | 18385.19 | 6754 |
| Spatial ability | A(S)-Level Sciences mean grade | 0.65 | 0.02 | 0.65 | 0.05 | 0.02 | 0 | 0.3 | 0.04 | 0.28 | 0.24 | 0.16 | 1 | 0.34 | 11 | 20472.98 | 7539 |
| Spatial ability | STEM pipeline | 0.65 | 0.15 | 0.44 | 0.05 | 0.02 | 0.04 | 0.3 | 0.01 | 0.34 | 0.39 | 0.51 | 0.53 | 0.14 | 11 | 29263.42 | 10928 |
| Spatial ability | Total number of STEM GCSEs taken | 0.63 | 0.11 | 0.48 | 0.07 | 0 | 0.18 | 0.29 | 0 | 0.22 | 0.31 | 0.43 | 0.1 | 0.13 | 11 | 44344.74 | 16975 |
| Spatial ability | Number of STEM A(S)-Levels taken | 0.67 | 0.13 | 0.54 | 0.04 | 0 | 0.03 | 0.3 | 0.01 | 0.3 | 0.35 | 0.43 | 0.34 | 0.16 | 11 | 46799.69 | 17613 |
| Spatial ability | GCSE humanities mean grade | 0.64 | 0.12 | 0.48 | 0.06 | 0.08 | 0.17 | 0.29 | 0.01 | 0.14 | 0.39 | 0.45 | 0.57 | 0.21 | 11 | 42595.06 | 16877 |
| Spatial ability | A(S)-Level humanities mean grade | 0.64 | 0 | 0.51 | 0.06 | 0.09 | 0 | 0.29 | 0 | 0.39 | 0.12 | 0.08 | 1 | 0 | 11 | 29221.84 | 10706 |
| Spatial ability | Total number of humanities GCSES taken | 0.63 | 0.02 | 0.5 | 0.07 | 0.21 | 0 | 0.29 | 0 | 0.26 | 0.26 | 0.21 | 1 | 0.03 | 11 | 44707.51 | 16975 |
| Spatial ability | Number of humanities A(S)-Levels taken | 0.62 | 0 | 0.58 | 0.08 | 0.05 | 0 | 0.3 | 0 | 0.37 | 0.1 | 0.07 | 1 | 0.02 | 11 | 47753.29 | 17613 |
| Spatial ability | Chose a STEM GCSE | 0.63 | 0.03 | 0.49 | 0.08 | 0.3 | 0 | 0.3 | 0 | 0.18 | | | | | 16 | 24924.61 | 16971 |
| Spatial ability | Chose a STEM A(S)-Level | 0.65 | 0.1 | 0.52 | 0.05 | 0.16 | 0.04 | 0.3 | 0.01 | 0.16 | | | | | 16 | 27059.3 | 17690 |
| Spatial ability | Chose a STEM degree | 0.69 | 0.14 | 0.39 | 0 | 0 | 0 | 0.29 | 0.01 | 0.46 | | | | | 13 | 17949.81 | 13676 |
| Spatial ability | Chose a humanities GCSE | 0.64 | 0.03 | 0.49 | 0.08 | 0.3 | 0 | 0.3 | 0 | 0.18 | | | | | 16 | 24924.61 | 16971 |
| Spatial ability | Chose a humanities A(S)-Level | 0.64 | 0.05 | 0.57 | 0.07 | 0.04 | 0.18 | 0.3 | 0 | 0.15 | | | | | 16 | 27886.99 | 17690 |
| Spatial ability | Chose a humanities degree | 0.71 | 0 | 0.57 | 0 | 0 | 0 | 0.29 | 0 | 0.42 | | | | | 13 | 18347.93 | 13676 |

**Table S13**. Trivariate twin model ACE path coefficients and fit indices. a = heritability, c = shared environment, e = non-shared environment. a/c/e11 = estimate of variance unique to variable 1, a/c/e21 = estimate of variance shared between variables 1 and 2, a/c/e31 = estimate of variance shared between variables 1 and 3, a/c/e22 = estimate of variance unique to variable 2 after accounting for shared variance with variable 1, a/c/e32 = estimate of variance shared between variables 2 and 3 after accounting for variance shared with variable 1, a/c/e33 = estimate of variance unique to variable 3 after accounting for variance shared with variables 1 and 2. Number suffixes indicate the variable the estimate or coefficient corresponds to, as denoted by the model column.

| Model | | | Standardized squared path coefficients | | | | | | | | | | | | | | | | | | Fit indices | | |
|---|---|---|---|---|---|---|---|---|---|---|---|---|---|---|---|---|---|---|---|---|---|---|---|
| 1 | 2 | 3 | a11 | a21 | a31 | a22 | a32 | a33 | c11 | c21 | c31 | c22 | c32 | c33 | e11 | e21 | e31 | e22 | e32 | e33 | ep | -2LL | df |
| g | Spatial ability | GCSE STEM mean grade | 0.72 | 0.41 | 0.38 | 0.23 | 0 | 0.23 | 0.09 | 0 | 0.09 | 0.05 | 0.05 | 0.09 | 0.16 | 0.02 | 0.02 | 0.27 | 0 | 0.11 | 21 | 47537.85 | 19705 |
| Verbal ability | | | 0.67 | 0.26 | 0.32 | 0.38 | 0.04 | 0.26 | 0.12 | 0 | 0.1 | 0.04 | 0.07 | 0.06 | 0.18 | 0.01 | 0.01 | 0.27 | 0.01 | 0.12 | 21 | 48233.81 | 19716 |
| g | Spatial ability | GCSE Maths mean grade | 0.72 | 0.4 | 0.36 | 0.23 | 0.01 | 0.27 | 0.09 | 0 | 0.08 | 0.06 | 0.01 | 0.09 | 0.17 | 0.02 | 0.02 | 0.27 | 0 | 0.15 | 21 | 48030.17 | 19576 |
| Verbal ability | | | 0.66 | 0.25 | 0.29 | 0.38 | 0.07 | 0.29 | 0.13 | 0 | 0.09 | 0.06 | 0.01 | 0.07 | 0.18 | 0.01 | 0.01 | 0.28 | 0.01 | 0.15 | 21 | 48751.65 | 19587 |
| g | Spatial ability | GCSE Science mean grade | 0.72 | 0.41 | 0.38 | 0.24 | 0 | 0.23 | 0.11 | 0 | 0.05 | 0.04 | 0.01 | 0.15 | 0.17 | 0.02 | 0.02 | 0.27 | 0 | 0.14 | 21 | 45948.58 | 18749 |
| Verbal ability | | | 0.67 | 0.26 | 0.36 | 0.4 | 0.03 | 0.23 | 0.13 | 0 | 0.05 | 0.04 | 0.01 | 0.15 | 0.19 | 0.01 | 0.01 | 0.28 | 0 | 0.14 | 21 | 46590.56 | 18760 |
| g | Spatial ability | GCSE Physics grade | 0.69 | 0.41 | 0.23 | 0.25 | 0 | 0.26 | 0.13 | 0 | 0.18 | 0.04 | 0.04 | 0.04 | 0.17 | 0.02 | 0.02 | 0.27 | 0 | 0.2 | 21 | 30102.53 | 11925 |
| Verbal ability | | | 0.67 | 0.25 | 0.17 | 0.4 | 0.04 | 0.28 | 0.14 | 0 | 0.2 | 0.05 | 0.07 | 0 | 0.19 | 0.01 | 0.01 | 0.28 | 0 | 0.2 | 21 | 30772.72 | 11936 |
| g | Spatial ability | GCSE Chemistry grade | 0.7 | 0.42 | 0.18 | 0.25 | 0.01 | 0.35 | 0.13 | 0 | 0.18 | 0.04 | 0.01 | 0.06 | 0.17 | 0.02 | 0.02 | 0.27 | 0 | 0.17 | 21 | 30100.92 | 11939 |
| Verbal ability | | | 0.67 | 0.26 | 0.21 | 0.4 | 0.02 | 0.32 | 0.14 | 0 | 0.09 | 0.04 | 0.03 | 0.13 | 0.19 | 0.01 | 0.01 | 0.28 | 0.01 | 0.18 | 21 | 30765.21 | 11950 |
| g | Spatial ability | GCSE Biology grade | 0.7 | 0.41 | 0.2 | 0.25 | 0 | 0.2 | 0.13 | 0 | 0.16 | 0.04 | 0.12 | 0.09 | 0.17 | 0.02 | 0.03 | 0.27 | 0 | 0.18 | 21 | 30129.8 | 11967 |
| Verbal ability | | | 0.67 | 0.25 | 0.2 | 0.41 | 0 | 0.2 | 0.14 | 0 | 0.1 | 0.04 | 0.17 | 0.1 | 0.19 | 0.01 | 0.02 | 0.28 | 0.01 | 0.18 | 21 | 30782.32 | 11978 |
| g | Spatial ability | A(S)-Level STEM mean grade | 0.68 | 0.4 | 0.16 | 0.25 | 0.03 | 0.43 | 0.15 | 0 | 0.02 | 0.05 | 0.01 | 0 | 0.17 | 0.02 | 0 | 0.27 | 0.01 | 0.33 | 21 | 30827.06 | 11918 |
| Verbal ability | | | 0.67 | 0.24 | 0.15 | 0.41 | 0 | 0.47 | 0.14 | 0 | 0.02 | 0.06 | 0.01 | 0 | 0.19 | 0.01 | 0 | 0.28 | 0.01 | 0.33 | 21 | 31469.76 | 11929 |
| g | Spatial ability | A(S)-Level Maths mean grade | 0.68 | 0.39 | 0.13 | 0.25 | 0.03 | 0.46 | 0.15 | 0.01 | 0.04 | 0.06 | 0.01 | 0 | 0.17 | 0.02 | 0 | 0.27 | 0.01 | 0.31 | 21 | 24530.96 | 9581 |
| Verbal ability | | | 0.66 | 0.23 | 0.15 | 0.41 | 0 | 0.48 | 0.15 | 0 | 0.02 | 0.06 | 0.02 | 0 | 0.19 | 0.01 | 0 | 0.28 | 0.01 | 0.21 | 21 | 25175.67 | 9592 |
| g | Spatial ability | A(S)-Level Sciences mean grade | 0.69 | 0.41 | 0.15 | 0.25 | 0.06 | 0.43 | 0.14 | 0 | 0.04 | 0.05 | 0.01 | 0 | 0.17 | 0.02 | 0.01 | 0.27 | 0.03 | 0.27 | 21 | 26561.38 | 10366 |
| Verbal ability | | | 0.67 | 0.23 | 0.12 | 0.41 | 0 | 0.51 | 0.14 | 0 | 0.04 | 0.05 | 0.01 | 0 | 0.19 | 0.01 | 0.02 | 0.28 | 0.02 | 0.26 | 21 | 27207.59 | 10377 |
| g | Spatial ability | STEM pipeline | 0.71 | 0.4 | 0.15 | 0.25 | 0.03 | 0.42 | 0.13 | 0 | 0.05 | 0.06 | 0 | 0 | 0.18 | 0.02 | 0 | 0.28 | 0 | 0.34 | 21 | 35352.1 | 13755 |
| Verbal ability | | | 0.68 | 0.24 | 0.11 | 0.41 | 0.07 | 0.41 | 0.13 | 0 | 0.06 | 0.06 | 0.01 | 0 | 0.19 | 0.01 | 0 | 0.29 | 0.01 | 0.34 | 21 | 35994.86 | 13766 |
| g | Spatial ability | Total number of STEM GCSEs taken | 0.68 | 0.39 | 0.15 | 0.24 | 0 | 0.43 | 0.14 | 0.01 | 0 | 0.06 | 0 | 0.17 | 0.17 | 0.02 | 0 | 0.27 | 0 | 0.22 | 21 | 50442.08 | 19802 |
| Verbal ability | | | 0.66 | 0.24 | 0.12 | 0.4 | 0.02 | 0.44 | 0.14 | 0 | 0.02 | 0.06 | 0 | 0.17 | 0.19 | 0.01 | 0 | 0.27 | 0 | 0.22 | 21 | 51073.85 | 19813 |
| g | Spatial ability | Number of STEM A(S)-Levels taken | 0.71 | 0.41 | 0.11 | 0.26 | 0.03 | 0.51 | 0.14 | 0 | 0.05 | 0.05 | 0 | 0 | 0.18 | 0.02 | 0 | 0.28 | 0.01 | 0.29 | 21 | 52868.93 | 20440 |
| Verbal ability | | | 0.68 | 0.25 | 0.08 | 0.42 | 0.06 | 0.51 | 0.15 | 0 | 0.05 | 0.04 | 0 | 0 | 0.2 | 0.01 | 0 | 0.29 | 0.01 | 0.29 | 21 | 53528.93 | 20451 |
| g | Spatial ability | GCSE humanities mean grade | 0.69 | 0.39 | 0.31 | 0.24 | 0.03 | 0.26 | 0.11 | 0 | 0.13 | 0.06 | 0.05 | 0.06 | 0.17 | 0.02 | 0.01 | 0.27 | 0 | 0.13 | 21 | 48041.99 | 19704 |
| Verbal ability | | | 0.66 | 0.24 | 0.36 | 0.39 | 0 | 0.23 | 0.13 | 0 | 0.1 | 0.06 | 0.08 | 0.07 | 0.18 | 0.01 | 0.01 | 0.28 | 0 | 0.14 | 21 | 48603.97 | 19715 |
| g | Spatial ability | A(S)-Level humanities mean grade | 0.69 | 0.39 | 0.07 | 0.25 | 0.03 | 0.38 | 0.14 | 0 | 0.1 | 0.06 | 0.02 | 0 | 0.17 | 0.02 | 0.01 | 0.27 | 0 | 0.38 | 21 | 35235.87 | 13533 |
| Verbal ability | | | 0.67 | 0.24 | 0.11 | 0.4 | 0.01 | 0.36 | 0.14 | 0 | 0.08 | 0.06 | 0.04 | 0 | 0.19 | 0.01 | 0.01 | 0.28 | 0 | 0.38 | 21 | 35854.55 | 13544 |
| g | Spatial ability | Total number of humanities GCSEs taken | 0.66 | 0.37 | 0.12 | 0.25 | 0.03 | 0.37 | 0.15 | 0.01 | 0.12 | 0.07 | 0.08 | 0 | 0.17 | 0.02 | 0 | 0.27 | 0 | 0.26 | 21 | 50646.71 | 19802 |
| Verbal ability | | | 0.63 | 0.21 | 0.14 | 0.4 | 0 | 0.38 | 0.16 | 0.01 | 0.13 | 0.08 | 0.07 | 0 | 0.18 | 0.01 | 0 | 0.28 | 0 | 0.26 | 21 | 51206.73 | 19813 |
| g | Spatial ability | Number of humanities | 0.68 | 0.39 | 0.05 | 0.23 | 0.03 | 0.49 | 0.15 | 0.01 | 0.03 | 0.07 | 0.02 | 0 | 0.18 | 0.02 | 0 | 0.27 | 0 | 0.38 | 21 | 53821.67 | 20440 |

| | | | | | | | | | | | | | | | | | | | | | | | |
|---|---|---|---|---|---|---|---|---|---|---|---|---|---|---|---|---|---|---|---|---|---|---|---|
| Verbal ability | | A(S)-Levels taken | 0.67 | 0.24 | 0.09 | 0.38 | 0.02 | 0.46 | 0.15 | 0 | 0.01 | 0.08 | 0.03 | 0 | 0.19 | 0.01 | 0 | 0.28 | 0 | 0.38 | 21 | 54382.28 | 20451 |
| g | Spatial ability | Chose a STEM GCSE | 0.71 | 0.41 | 0.29 | 0.25 | 0 | 0.31 | 0.15 | 0.01 | 0.04 | 0.06 | 0.02 | 0.23 | 0.18 | 0.02 | 0 | 0.27 | 0 | 0.11 | 21 | 26138.615 | 19804 |
| Verbal ability | | | 0.69 | 0.25 | 0.21 | 0.41 | 0.04 | 0.35 | 0.15 | 0.00 | 0.10 | 0.06 | 0.04 | 0.16 | 0.19 | 0.01 | 0.00 | 0.28 | 0.00 | 0.10 | 21 | 26801.627 | 19815 |
| g | Spatial ability | Chose a STEM A(S)-Level | 0.67 | 0.38 | 0.12 | 0.25 | 0.02 | 0.49 | 0.14 | 0.00 | 0.16 | 0.06 | 0.03 | 0.00 | 0.18 | 0.02 | 0.00 | 0.27 | 0.01 | 0.16 | 21 | 33155.352 | 20523 |
| Verbal ability | | | 0.66 | 0.22 | 0.08 | 0.41 | 0.06 | 0.49 | 0.16 | 0.00 | 0.16 | 0.07 | 0.04 | 0.00 | 0.19 | 0.01 | 0.00 | 0.28 | 0.01 | 0.16 | 21 | 33796.575 | 20534 |
| g | Spatial ability | Chose a STEM degree | 0.69 | 0.40 | 0.12 | 0.25 | 0.02 | 0.32 | 0.12 | 0.00 | 0.00 | 0.04 | 0.01 | 0.04 | 0.18 | 0.02 | 0.01 | 0.27 | 0.00 | 0.46 | 21 | 24116.098 | 16506 |
| Verbal ability | | | 0.67 | 0.24 | 0.05 | 0.41 | 0.08 | 0.32 | 0.12 | 0.00 | 0.04 | 0.04 | 0.02 | 0.00 | 0.19 | 0.01 | 0.01 | 0.28 | 0.00 | 0.46 | 21 | 24763.476 | 16517 |
| g | Spatial ability | Chose a humanities GCSEs | 0.69 | 0.38 | 0.20 | 0.25 | 0.07 | 0.25 | 0.24 | 0.01 | 0.07 | 0.07 | 0.22 | 0.00 | 0.18 | 0.02 | 0.00 | 0.28 | 0.00 | 0.18 | 21 | 30992.373 | 19804 |
| Verbal ability | | | 0.67 | 0.23 | 0.25 | 0.40 | 0.02 | 0.26 | 0.15 | 0.00 | 0.06 | 0.08 | 0.23 | 0.00 | 0.19 | 0.01 | 0.00 | 0.28 | 0.00 | 0.18 | 21 | 31574.957 | 19815 |
| g | Spatial ability | Chose a humanities A(S)-Level | 0.69 | 0.40 | 0.15 | 0.25 | 0.01 | 0.46 | 0.14 | 0 | 0.07 | 0.06 | 0.02 | 0.14 | 0.18 | 0.02 | 0 | 0.27 | 0 | 0.14 | 21 | 33927.132 | 20523 |
| Verbal ability | | | 0.67 | 0.24 | 0.22 | 0.40 | 0 | 0.41 | 0.15 | 0 | 0.03 | 0.07 | 0.03 | 0.16 | 0.19 | 0.01 | 0 | 0.28 | 0 | 0.14 | 21 | 34517.15 | 20534 |
| g | Spatial ability | Chose a humanities degree | 0.69 | 0.41 | 0.08 | 0.24 | 0.1 | 0.38 | 0.13 | 0 | 0 | 0.06 | 0.01 | 0 | 0.18 | 0.02 | 0.01 | 0.27 | 0 | 0.42 | 21 | 24483.225 | 16506 |
| Verbal ability | | | 0.67 | 0.26 | 0.12 | 0.40 | 0.05 | 0.40 | 0.14 | 0 | 0 | 0.05 | 0 | 0 | 0.19 | 0.01 | 0 | 0.28 | 0 | 0.42 | 21 | 25081.14 | 16517 |

**Fig. S1.** Item and factor loadings for the hierarchical model of spatial ability. This model is a replication of the best-fitting model discovered using exploratory factor analysis in our previous work (see reference 17 in main text). CS = cross sections, 2D = 2D drawing, PA = pattern assembly, PT = perspective taking, MR = mechanical reasoning, PF = paper folding, 3D = 3D drawing, SR = shape rotation, EM = Elithorn mazes, M = mazes, Sc = scanning, LPT = large-scale perspective taking, OD = orientation directions, OL = orientation landmarks, MRm = map reading (memory), MRnm = map reading (no memory). Model fit indices: AIC = 52396.81; $\chi 2$ = 435.097 (102), $p < 0.001$; CFI = 0.946; TLI = 0.936; RMSEA = 0.041; SRMR = 0.049.

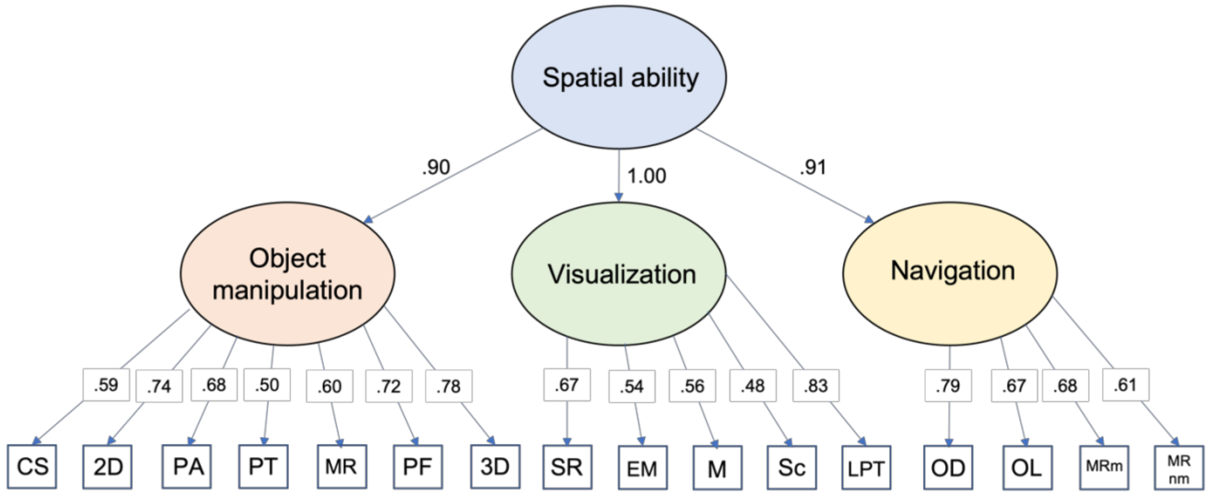

**Fig. S2.** Item loadings for the unifactorial model of *g*. Model fit indices: AIC = 17748.64; $\chi^2$ = 107.895 (14), $p < 0.001$; CFI = 0.961; TLI = 0.944; RMSEA = 0.068; SRMR = 0.036.

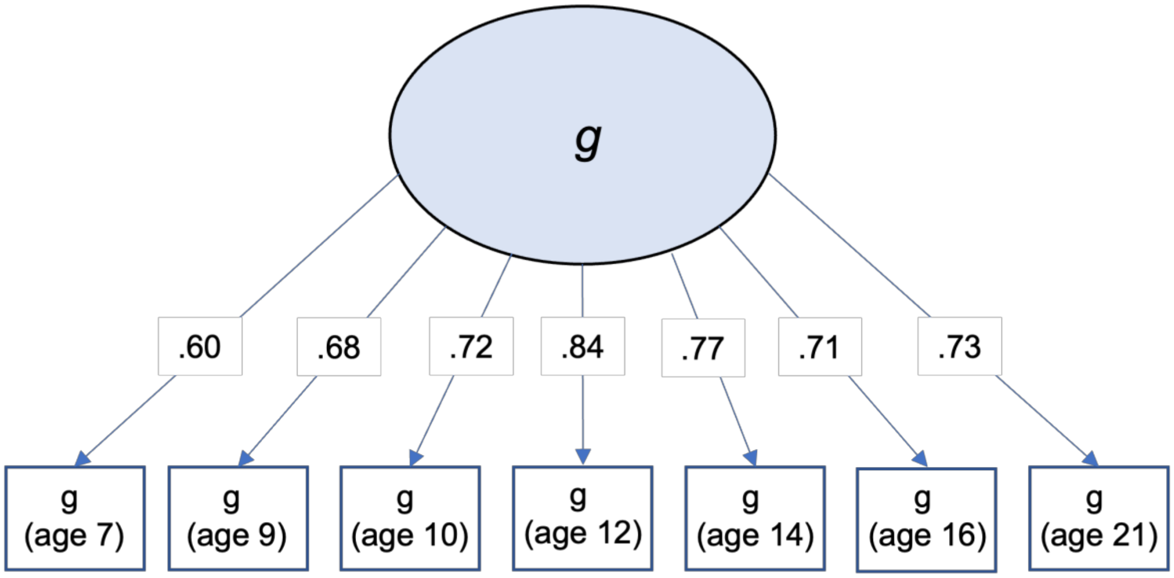

**Fig. S3.** Item loadings for the unifactorial model of verbal ability. Model fit indices: AIC = 18921.655; $\chi^2$ = 71.641 (14), p < 0.001; CFI = 0.970; TLI = 0.955; RMSEA = 0.055; SRMR = 0.034.

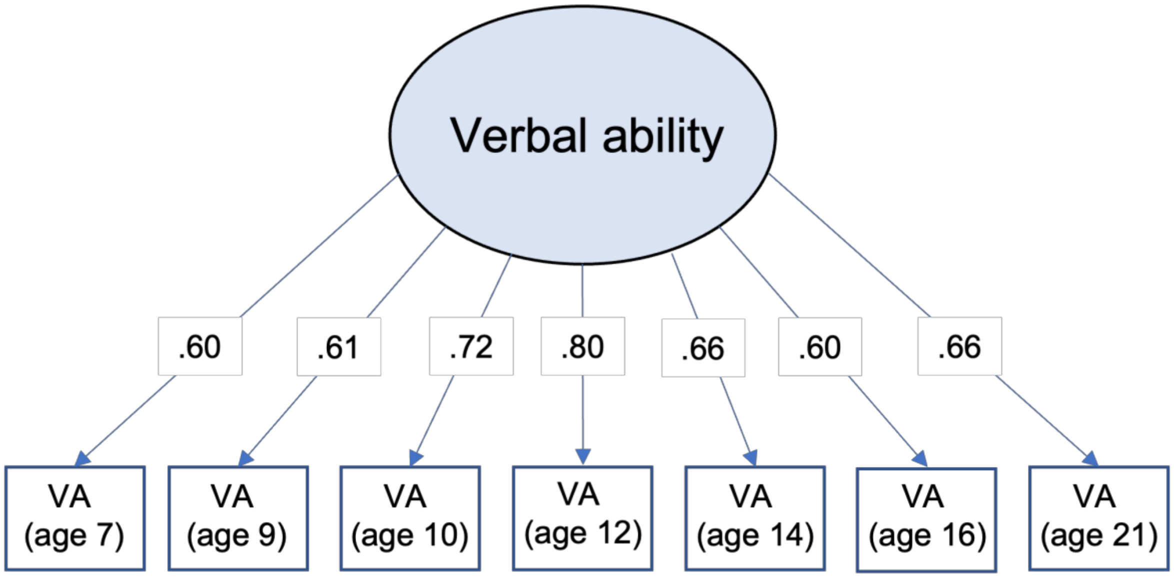

**Fig S4.** Correlation matrix denoting correlations between all cognitive predictor variables. spatab = spatial ability; nav = navigation; vis = vis, objmanip = object manipulation, v = verbal ability. .g and .v suffixes denote *g*- and v-corrected predictors, respectively.

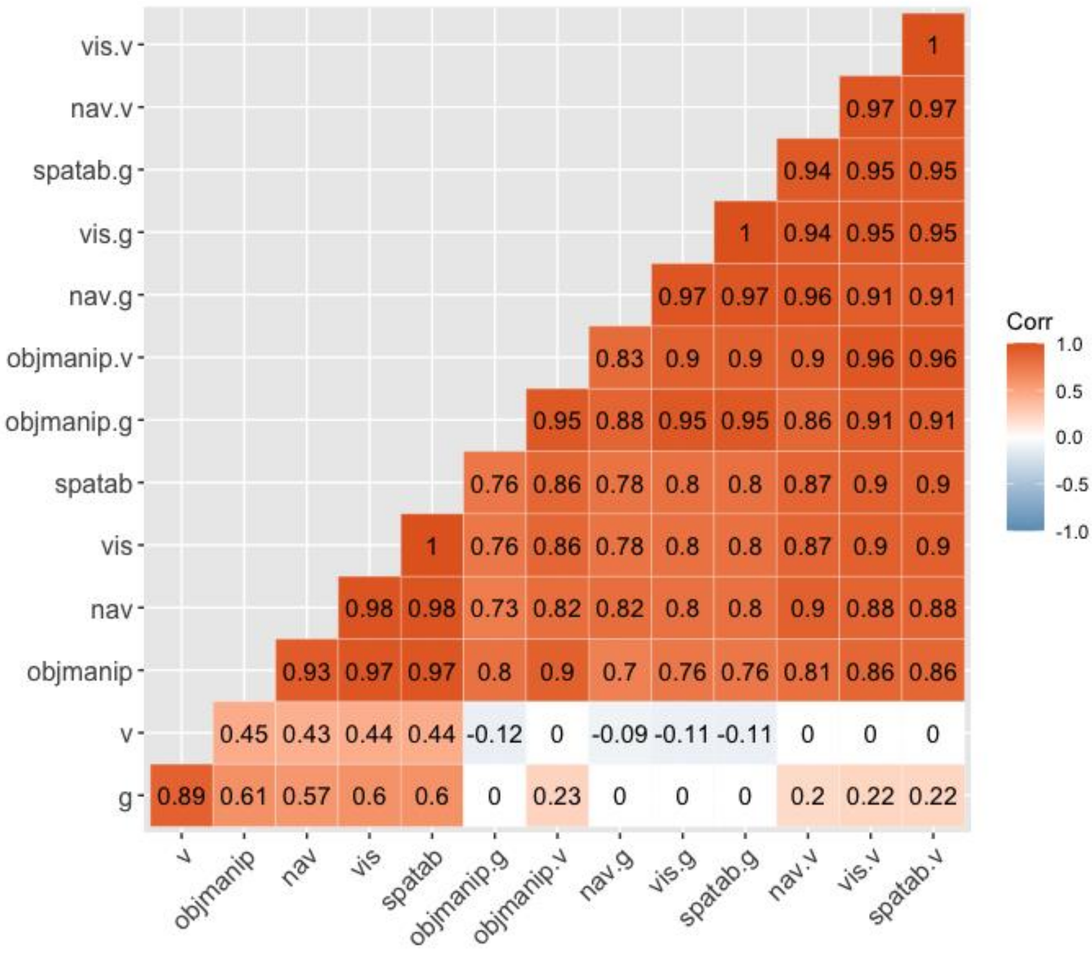

**Fig S5.** Spaghetti plot visualizing progression through the STEM pipeline. Each line represents an individual, with its position along the y-axis representing the individual's spatial ability score and the length of the line denoting their STEM pipeline score, where 0 = completed compulsory schooling (i.e., all individuals with complete data), 1 = did not choose a non-compulsory STEM GCSE, 2 = chose a non-compulsory STEM GCSE, 3 = chose a STEM A(S)-Level, and 4 = chose a STEM Bachelor's degree (or higher).

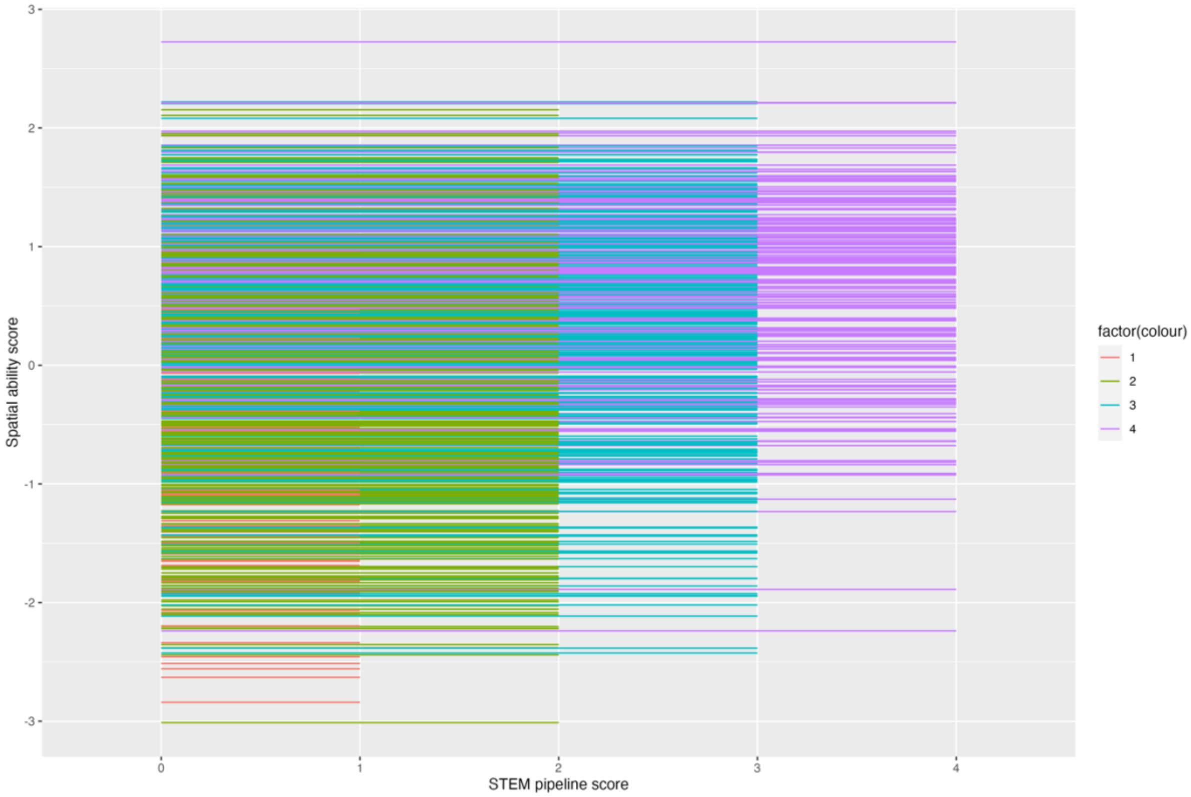

**Fig S6.** Standardized squared path estimates from bivariate Cholesky decompositions examining the shared environmental (C) overlap between spatial ability and educational outcomes. The total bar length represents the total shared environmental contribution to variance in the outcome, which is decomposed into that which is unique to the outcome and that which is shared with spatial ability.

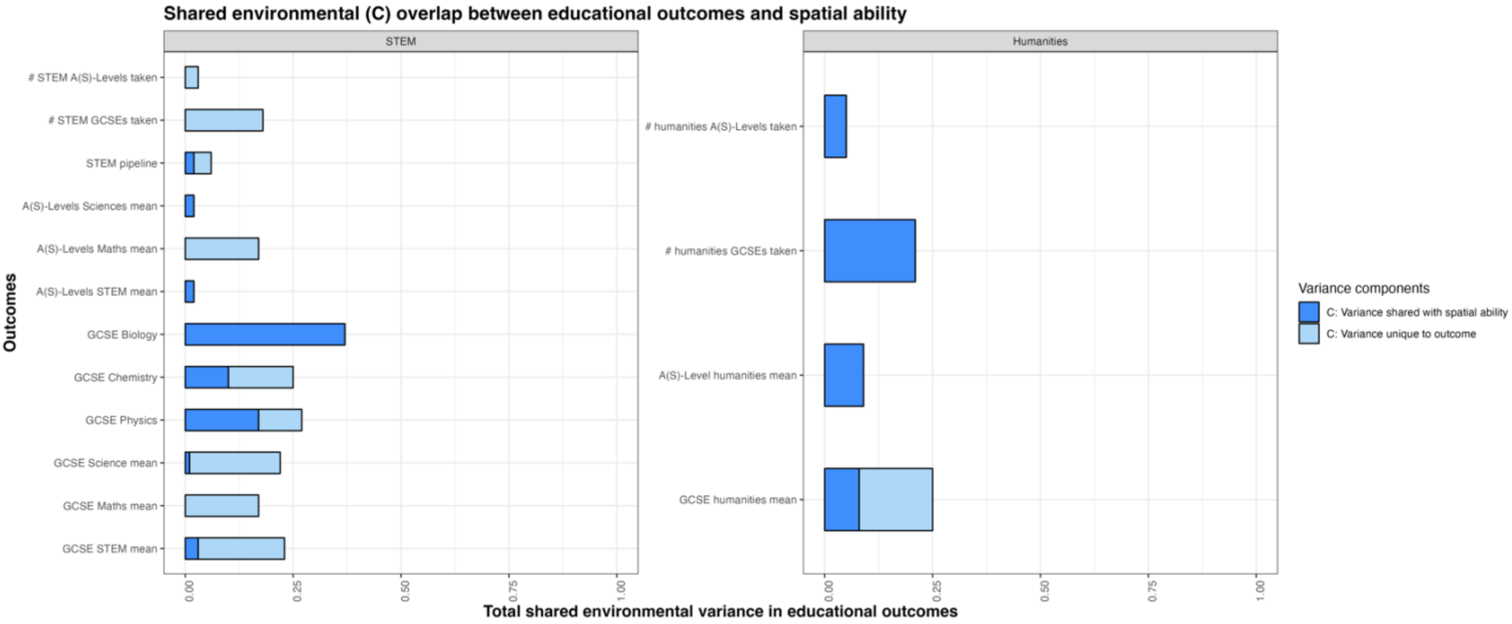

**Fig S7.** Standardized squared path estimates from bivariate Cholesky decompositions examining the unique environmental overlap between spatial ability and educational outcomes. The total bar length represents the total unique environmental contribution to variance in the outcome, which is decomposed into that which is unique to the outcome and that which is shared with spatial ability.

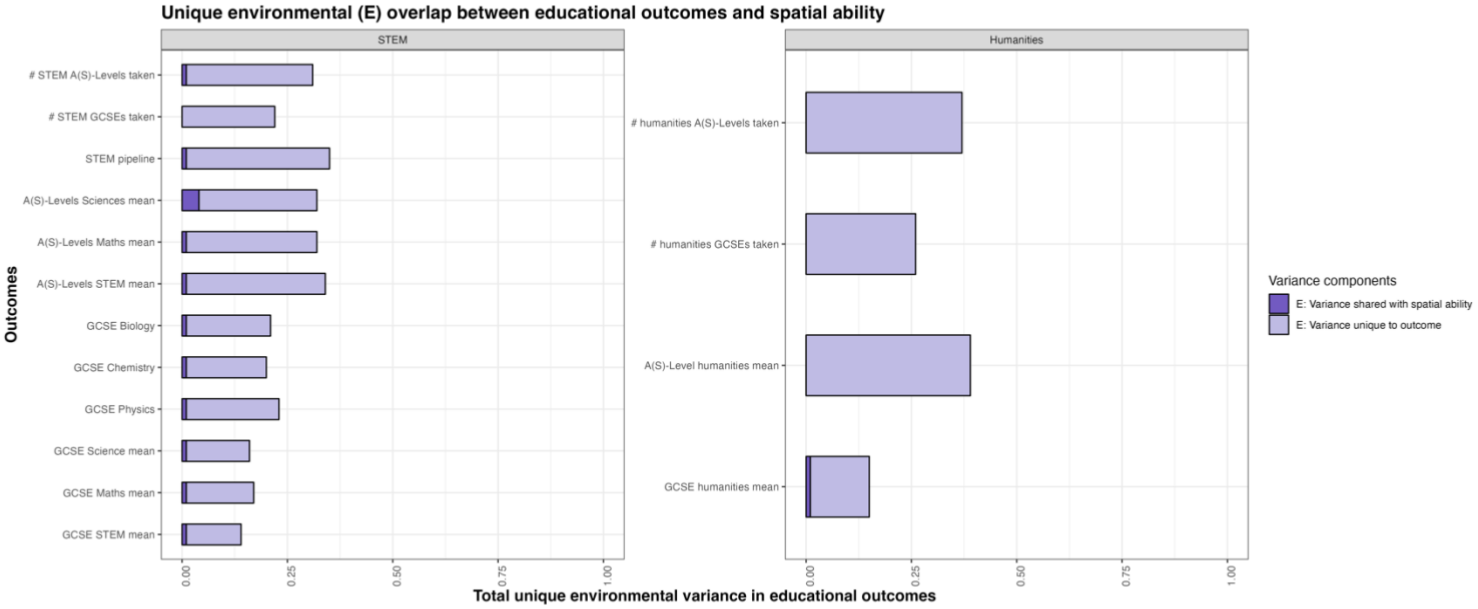

**Fig S8.** Standardized squared path estimates from trivariate Cholesky decompositions examining the aetiological overlap between verbal ability, spatial ability, and educational. The total bar length represents the total variance in the outcome, which is decomposed into A, C, and E components that are unique to the outcome, shared with verbal ability, and uniquely shared with spatial ability after accounting for verbal ability.

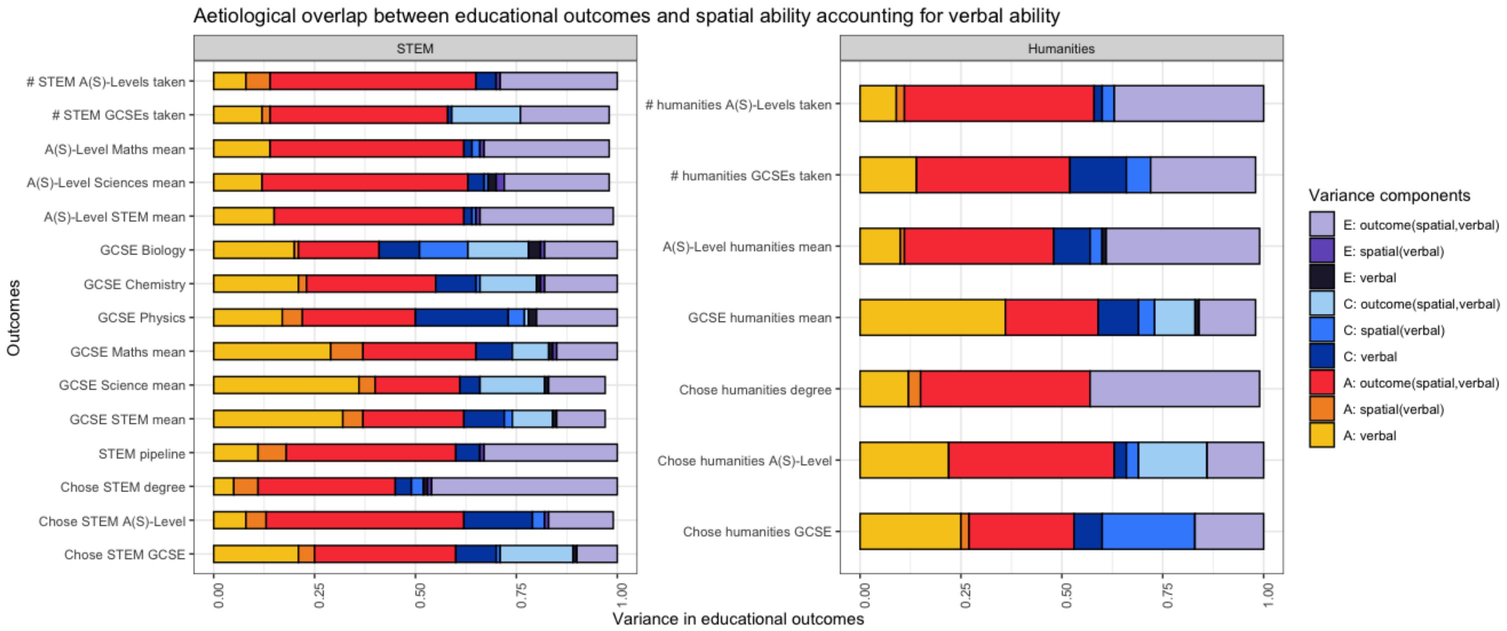

**Fig S9.** Standardized squared path estimates from trivariate Cholesky decompositions examining the aetiological overlap between g, spatial ability, and educational. The total bar length represents the total variance in the outcome, which is decomposed into A, C, and E components that are unique to the outcome, shared with *g*, and uniquely shared with spatial ability after accounting for *g*.

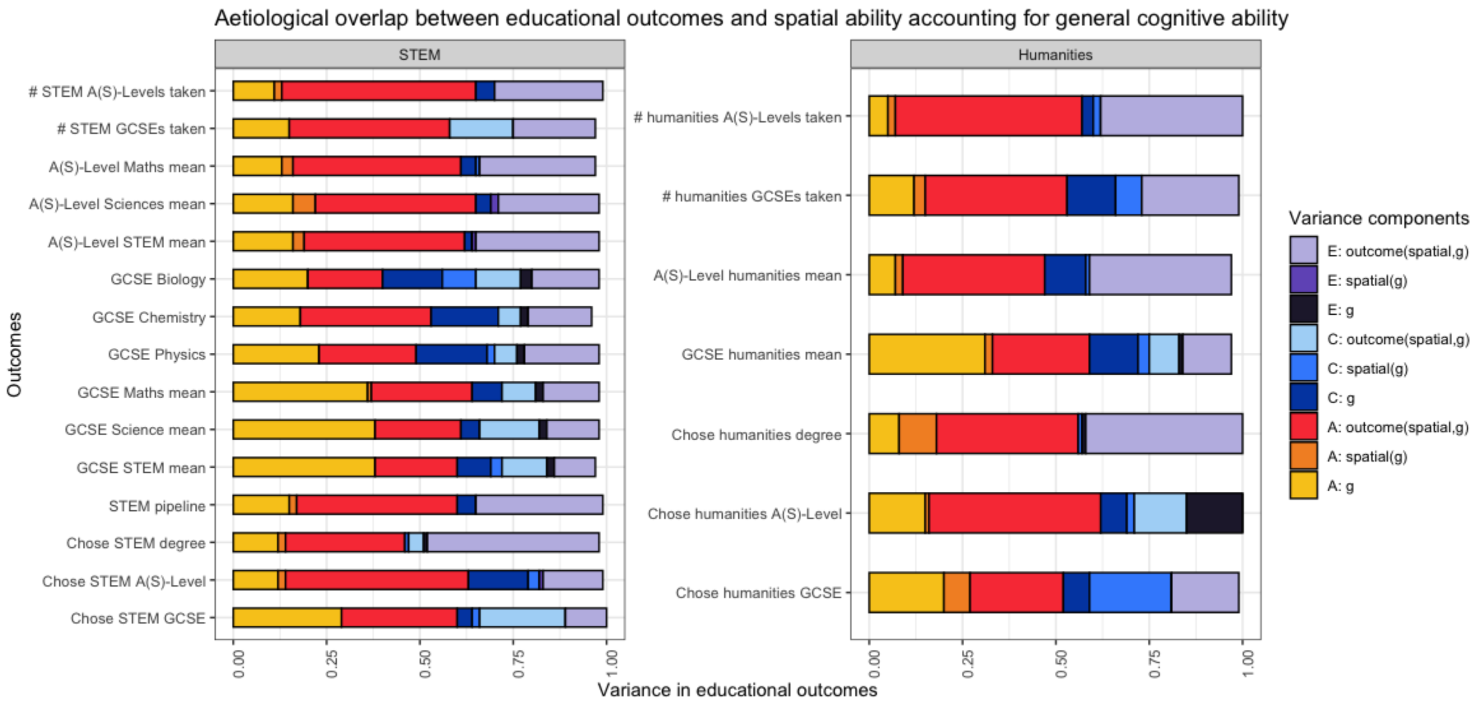
